# Promoter G-quadruplexes and transcription factors cooperate to shape the cell type-specific transcriptome

**DOI:** 10.1101/2020.08.27.236778

**Authors:** Sara Lago, Filippo M. Cernilogar, Maryam Kazerani, Helena Domíniguez Moreno, Matteo Nadai, Gunnar Schotta, Sara N. Richter

## Abstract

Cell identity is maintained by activation of cell-specific gene programs, regulated by epigenetic marks, transcription factors and chromatin organization^1-3^. DNA G-quadruplex (G4)-folded regions in cells were reported to be associated with either increased or decreased transcriptional activity^4,5^. By G4 ChIP-seq/RNA-seq analysis on liposarcoma cells we confirmed that G4s in promoters are invariably associated with high transcription levels in open chromatin. Comparing G4 presence, location and transcript levels in liposarcoma cells to available data on keratinocytes, we showed that the same promoter sequences of the same genes in the two cell lines had different G4-folding state: high transcript levels consistently associated with high G4-folding. Transcription factors AP-1 and SP1, whose binding sites were the most significantly represented in G4-folded sequences, coimmunoprecipitated with their G4-folded promoters. Thus G4s and their associated transcription factors cooperate to determine cell-specific transcriptional programs, making G4s strongly emerge as new epigenetic regulators of the transcription machinery.

## Main

DNA G4s have been indicated as physical obstacles for RNA polymerase processivity, and thus transcriptional repressors in vitro^6-9^. However, recent evidence in cells suggests that several factors are involved in the outcome of G4-mediated transcription regulation: the template/coding strand harbouring the G4 sequence^10^, transcription factors (TFs)^11-13^, chromatin remodelling proteins^14^, interaction with distal genomic loci^15^; in this context, G4s where rather linked to high gene expression^4,16^. Since G4s are dynamic structures that can be induced or unfolded upon interaction with G4-binding proteins, we reasoned that the G4 landscape could be unique to each transcriptional program, hence cell type. We thus compared the G4 landscape in two cell types and explored the specific interactions at the G4-folded and transcriptionally active sequences to ascertain the mechanism behind G4-mediated transcription regulation.

We investigated by G4 ChIP-seq^17^ the *in vivo* DNA G4-folding in a cancer line, the human well-differentiated liposarcoma (WDLPS) 93T449 cell line, and compared data to those previously reported for the spontaneously immortalized keratinocyte HaCaT cell line from adult human skin^4^. G4 ChIP in WDLPS cells was first validated by quantitative real-time PCR (qPCR) on control sequences (**Supplementary Figure 1A**), then subjected to Next-Generation Sequencing yielding about 5,000 peaks that described the WDLPS genome-wide G4 landscape **(Supplementary File 1)**. GC content was significantly enriched in ChIP sample (~55%) *vs* the whole genome^18^ (~41%) **(Supplementary Figure 1B)**, with the highest GC frequency increasing towards the ChIP peak centre **(Supplementary Figure 1C)**. The presence of G4s within ChIP peaks was assessed through Quadparser^19^ and G4Hunter^20^. The mean length of the 7,400-10,000 identified G4s was 21-36 bp **(Supplementary Table 1)**, corresponding to that of the most stable G4s *in vitro*^21^. An average of 77% ChIP peaks contained at least one predicted G4 **(Supplementary Figure 2A-B)**, most contained 1-6 G4s, reaching 15 G4s in some sequences, with equivalent distribution between DNA strands **(Supplementary Figure 2C-D and Supplementary Table 1)**.

Non-random distribution of G4s in the human genome has been previously reported^22^. In line with this, ChIP-G4s in 93T449 cells were strikingly prevalent in promoter regions (79%) **(Figure 1A)**, fact that strongly supports G4 involvement in transcription regulation. RNA-seq data (**Supplementary File 2**) and subsequent analysis of the percentage of expressed genes that contained at least one ChIP-G4 *vs* G4-location showed that: i) G4-containing genes were more prone to be actively expressed with respect to G4-depleted genes; ii) genes with G4s in promoter regions (annotated as promoter-TSS and 5’-UTR) were almost 100% expressed **(Figure 1B)** and iii) produced significantly higher amounts of transcripts **(Figure 1C)**. In particular, there was positive correlation between higher gene expression and the presence of G4s, especially when G4s were within 1 kb from the TSS (**Supplementary Figure 3A and B**); this trend was smoothed as the distance from the TSS increased **(Figure 1D, top)**. Moreover, G4s mostly clustered within 250 bp from the TSS and the closer they were to the TSS, the higher the expression of the corresponding gene **(Figure 1D, bottom)**. Representative regions showing the relative position of G4s with respect to gene TSS are reported in **Figure 1F**.

**Figure 1.**
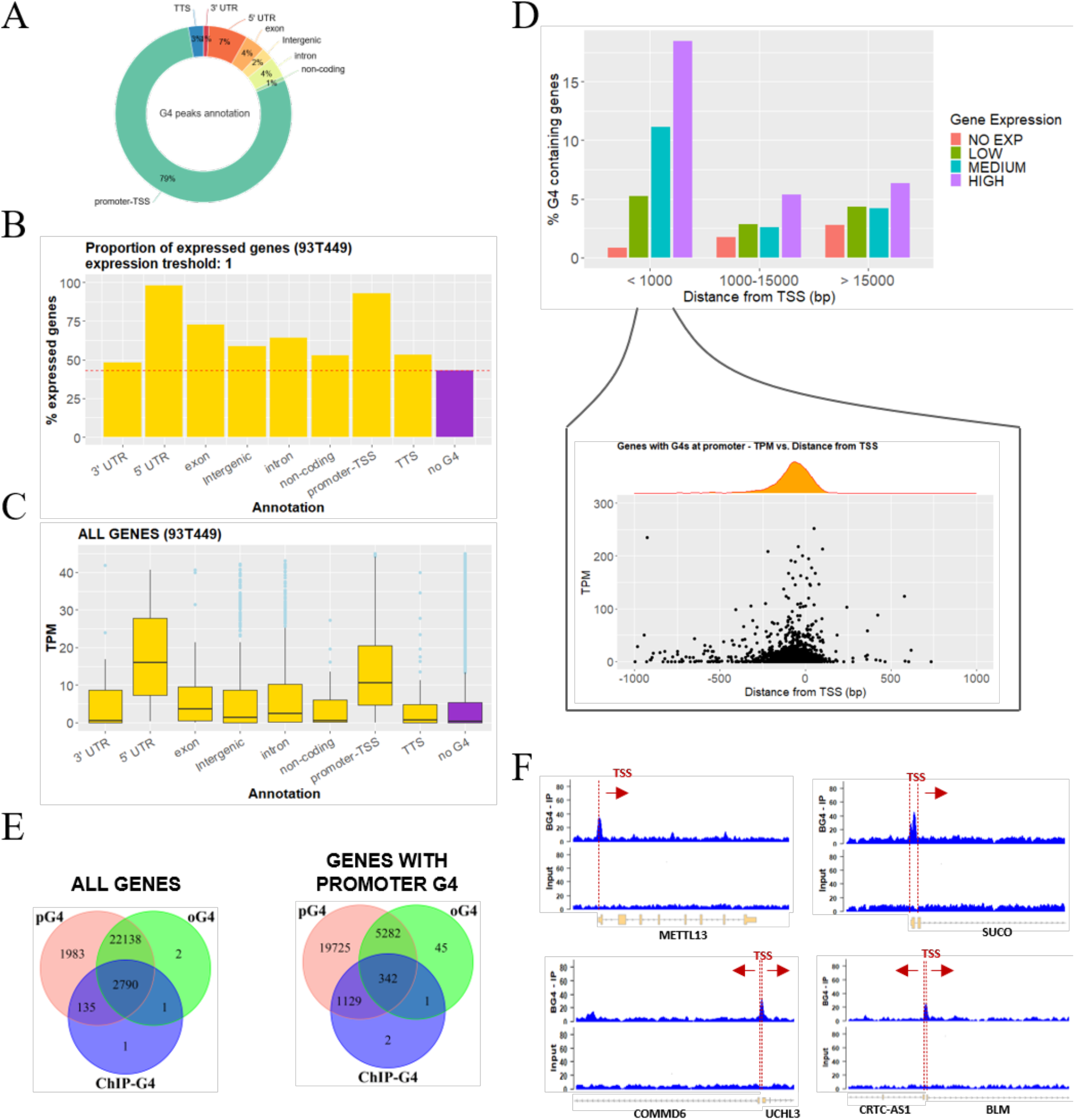
Genomic position of G4s and association to gene expression. **A)** Percentage distribution of G4 peaks in functional genomic regions according to HOMER gene annotation. Percentages are normalized over the genomic abundance of each functional region. **B)** Percentage proportion of expressed genes among the G4-containing genes (yellow). G4 depleted genes (no G4, violet) are reported as reference. One transcript per gene was considered as threshold. **C)** Gene expression distribution expressed in transcripts per million (TPM) of all the G4-containing genes (yellow). Genes were grouped according to the functional annotation of the immunoprecipitated G4 region. G4 depleted genes (no G4, violet) are reported as reference. **D)** Upper panel: Percentage of G4 containing genes, in genes grouped according to their expression level (no expression, low, medium or high) and their distance from the TSS of the closest gene (< 1000 bp, between 1000-15000 bp and > 15000 bp). Lower panel: detailed view of gene expression level (TPM) and density distribution of genes with folded G4s within 1000 bp from TSS in function of the G4 distance from the TSS. **E)** Venn diagram showing the intersection of genes which harbor pG4s, oG4s and ChIP-G4s in any positions (left) or in their promoter (right). **F)** Genomic view of representative regions showing the G4-ChIP peak position with respect to the TSS: G4-ChIP peaks in two gene promoters with non-coding upstream regions are displayed in the upper panels (METTL13 and SUCO); G4-ChIP peaks embedded in the coding regions of two adjacent genes with opposite transcription direction are shown in the lower panels (COMMD6 and UCHL3 – left; CRTC-AS1 and BLM – right).

We next compared the amount of ChIP-G4s to that of putative G4s (pG4s) calculated by Quadparser and of “observed G4s” (oG4s), i.e. genomic regions previously observed to stop polymerase progression *in vitro*^23^. Almost all ChIP-G4s were also found as pG4s (~100%) and oG4s (~95%) when considering all genes; in gene promoters a similar percentage was found for ChIP-G4s matching pG4s (~100%), while a much lower percentage was found for oG4s (~23%) (**Figure 1E**). This difference can be explained by the fact that the majority of ChIP-G4s correspond to canonical G4 motifs (**Supplementary Figure 2A**), while more than 60% of oG4s contained long loops or non-canonical patterns^23^. When analyzing the expression level of promoters in these three G4 categories, we found a significant increase of transcripts in ChIP-G4s. The other categories yielded significantly lower levels (**Supplementary Figure 4**). These data indicate that only the sequences that are actually folded into G4 are associated to high transcript levels, while pG4s or unfolded G4-forming sequences are not.

Analysis of accessible chromatin regions by Omni-ATAC-seq^24^ showed that 93% of promoter ChIP-G4s (**Figure 2B**) and 70% of all ChIP-G4s (**Supplementary Figure 5A**) overlapped with open chromatin (**Figure 2A, Supplementary File 3**); 83% of open chromatin regions were embedded in gene promoters (**Supplementary Figure 5B**), however, only 12% of these overlapped with G4s (**Figure 2B**), suggesting that the G4 regulatory role is limited to specific DNA region.

**Figure 2.**
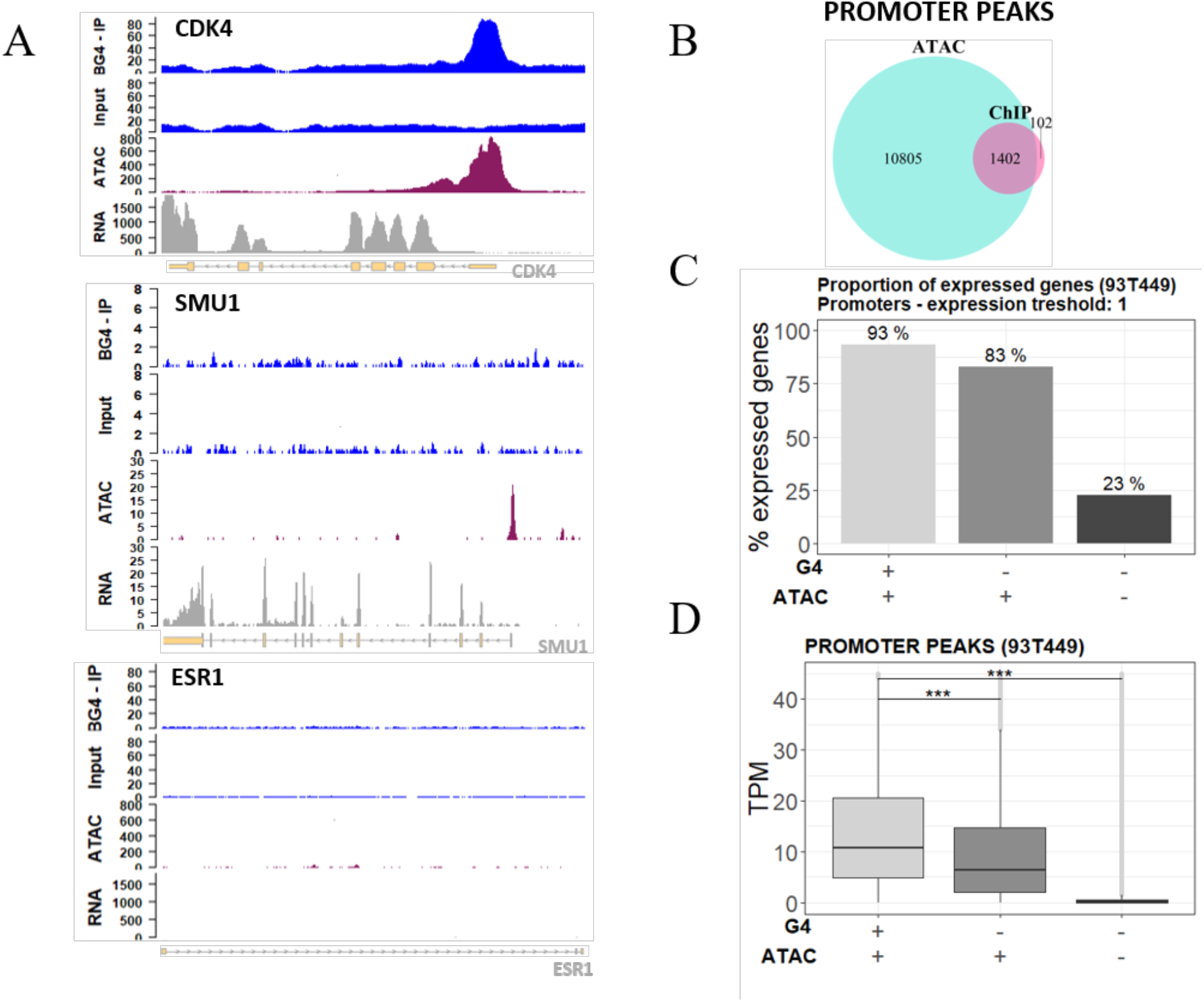
Relationship between G4s and open chromatin. **A)** Genomic view showing input and G4 IP samples, ATAC-seq and RNA-seq peaks in the promoter of representative genes: CDK4 (upper panel), SMU1 (mid panel), ESR-1 as negative control gene (lower panel). **B)** Venn diagram displaying the intersection between peak regions corresponding to IP G4s (light blue) and open chromatin regions (violet) mapped by ATAC-seq in promoters. **C)** Percentage proportion of expressed genes grouped according to the presence of G4s and open chromatin signal in their promoter region. One transcript per gene was considered as expression threshold. **D)** Expression distribution of all genes grouped according to the presence of ChIP-seq G4s and ATAC-seq signals in their promoter region. Gene expression is reported as TPM. In C) and D) the presence and absence of G4 and ATAC-seq signals are indicated below the graphs.

In 93T449 cells, 90% of genes with ChIP-G4s in their promoter (i.e. G4-folded promoters) and embedded in open chromatin were actively expressed; around 80% of genes with G4-depleted promoters in open chromatin and 25% of genes with G4-depleted promoters in closed chromatin were also actively expressed (**Figure 2C**). When analysing transcript abundance, we observed that the G4-folded promoters displayed higher transcriptional activity in open chromatin (**Figure 2D**). The same trend was maintained in the other functional regions (**Supplementary Figure 5C-D**). These results suggest that G4s stimulate recruitment of the transcription machinery/chromatin remodelling factors; alternatively, G4-folding could be a control mechanism to restrain/reinitiate transcription in highly active genes.

Specific gene programs are activated in cells to maintain their diversity and identity. Chromatin organization and gene expression are modulated by a complex interplay between epigenetic marks (DNA methylation, histone modification, nucleosome positioning) and binding of core TFs to regulatory DNAs^1-3^. We hypothesized that G4s cooperated with TFs in the maintenance of cell-specific gene expression programs, contributing to the establishment of the transcriptome. To verify this issue, we compared transcriptome, chromatin state and ChIP-G4s in 93T449 cells *vs* the immortalized keratynocytes cell line HaCaT (**Supplementary File 4, 5, 6**)^4^. The number of ChIP-G4s detected in the two cell lines was highly different (~5,000 in 93T449 *vs* ~30,000 in HaCaT). Based on the above collected data, i.e. that G4s are associated with high transcription levels, we reasoned that cell growth rate could be a determining factor: indeed 93T449 cells displayed doubling time twofold lower (3.3 ± 0.3 days) than that of HaCaT cells (1.5 ± 0.2 days) (**Supplementary Figure S6**). Notwithstanding the substantially lower amount of 93T449 G4s, the identity and distribution of the majority of them was cell-specific, i.e. not shared with HaCaT cells (**Figure 3D**) and independent of chromosome length and number of encoded genes (**Supplementary Figure S7A**).

**Figure 3.**
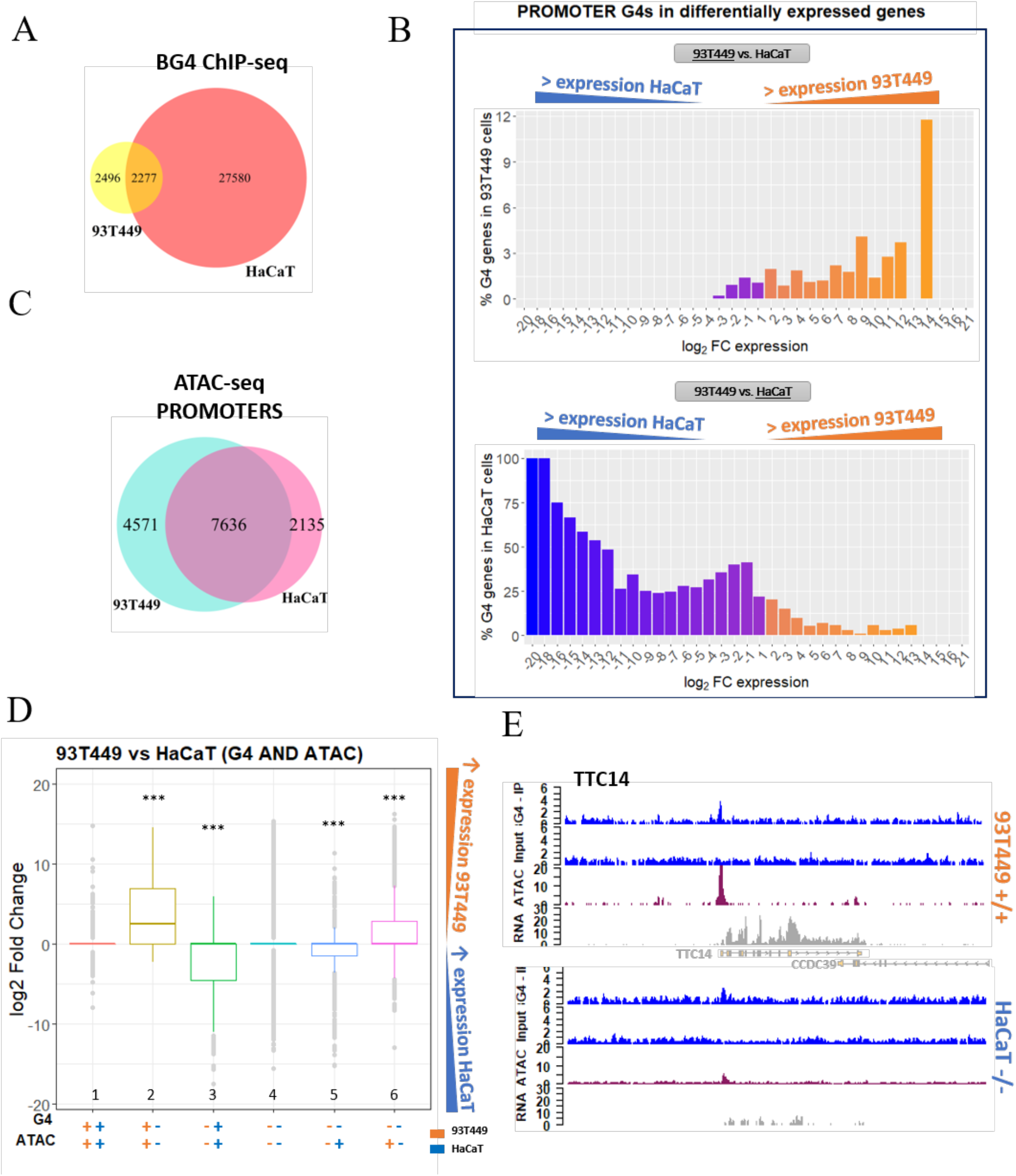
Comparison of 93T449 and HaCaT cell lines. **A)** Venn diagram showing the intersection between G4 peaks found in 93T449 and HaCaT cell lines. **B)** Percentage of genes containing at least one G4 in their promoter in 93T449 (upper panel) or HaCaT (lower panel) cells, in function of their differential expression in the two cell lines. Orange and blue-violet bars in both panels correspond to genes that have higher expression in 93T449 cells and HaCaT cells, respectively. **C)** Venn diagram showing the intersection between the ATAC-seq peaks found in promoters of 93T449 and HaCaT cell lines. **D)** Differential gene expression comparison of the same genes in 93T449 and HaCaT cells, based on the presence of G4s and open chromatin combinations. Orange and blue symbols indicate data for 93T449 and HaCaT cells, respectively. The presence (+) or absence (-) of G4s or ATAC signals are reported. Bars indicate gene expression distribution of the differentially expressed genes in 93T449 vs HaCaT cells. (*** = p-value < 0.001). **E)** Genomic view showing representative regions of 93T449 and HaCaT cell lines gene expression (RNA-seq track) with respect to the presence of G4 peaks (ChIP-seq) and open chromatin (ATAc-seq). In particular, TTC14 gene is displayed, which shows both G4 and ATAC signals in 93T449 cells, while it shows limited accessibility and no G4 in HaCaT cells. These differences are reflected in the corresponding RNA amount, which is much lower in HaCaT cells.

The two cell lines showed a largely different expression pattern^4,25^: in 93T449 cells, 5541 and 3560 genes were expressed to a significantly higher and lower extent, respectively, compared to the same genes in HaCaT (**Supplementary File 7**). Importantly, promoters contained folded G4s only in genes expressed at higher rates, while they were mainly unfolded when the same genes were downregulated (**Figure 3B**). Despite associated to lower transcriptional output with respect to genes with promoter G4s, when G4s folded outside the promoter of the respective gene, we observed the same trend of differential expression when comparing the two cell lines (**Supplementary Figure 7C**). This suggests that G4s sustain transcription also when folded outside promoters, by facilitating polymerase progression, double helix opening in gene coding sequence, enhancer/promoter interaction in intergenic regions. Together, these data indicate that different cell types modulate G4 folding to establish or maintain their transcriptional program.

Promoters (**Figure 3A)** and other genetic regions **(Supplementary Figure 7B**) associated with open chromatin were also different between the two cell lines. Analysis of the extent of different transcription levels in 93T449 *vs* HaCaT cells, evaluated according to the presence of folded G4s and open chromatin signals in promoters, indicated that the presence of both folded G4s and open chromatin strongly contributes to enhanced transcription levels (**Figure 3D and E,** bars 2 and 3), to a higher extent with respect to regions with open chromatin only (**Figure 3D,** bars 5 and 6; **Supplementary File 5**). Example regions for each of the described condition are reported in **Supplementary Figure 8.** These data indicate that G4s are involved in the recruitment of the transcriptional machinery or facilitate its function in open chromatin regions^26^.

To test the above hypothesis that G4s in promoters stimulate TF binding, we first predicted the presence of putative TF binding sites (TFBS) within the ChIP-G4s in 93T449 cells using HOMER tool (http://homer.ucsd.edu/homer/). The most relevant TFBS corresponded to AP-1 (24.08% in ChIP *vs* 3.36% in background sequences) and Sp1 (24.98% in ChIP vs 9.45% in background sequences) (**Figure 4A, Supplementary Files 8.1.1, 8.2.1**). AP-1 TFBS were mostly centred at the G4 peak, while Sp1 TFBS were more broadly distributed within the peak, suggesting that they can either overlap or be adjacent to the G4 (**Figure 4B**). When TFBS were calculated on an extended promoter region (−1000 to +750 bp from TSS) of either ChIP-G4-enriched or depleted genes, different TFBS were found and with consistently lower representation **(Supplementary Figure 9**). When considering all the target genes of AP-1 and Sp1 (data from ENCODE database), those presenting G4s (ChIP-G4s, pG4s and oG4s) were highly enriched with respect to those lacking G4s; among G4-containing categories, ChIP-G4s were the most enriched, suggesting association between TF binding and G4-folding capacity (**Figure 4C).** As a further proof, AP-1 and SP1 were co-immunoprecipitated using anti-G4 primary antibody (**Figure 4D**, lanes 2-3) and, vice versa, primary antibodies against SP1 and AP-1 co-immunoprecipitated the anti-G4 antibody added to the samples as marker of G4 regions (**Figure 4D**, lanes 4-8), proof that these TFs interact with their binding sites in the presence of folded G4s (**Figure 4D)**. These data indicate that TFs AP-1 and SP1 are strictly linked to G4s and suggest that G4s are exploited by cells to facilitate TF interaction with the DNA: this effect is reached by G4-mediated exposure of the binding region or altered DNA methylation state, as G4s were reported to prevent CpG island methylation, condition which is required for transcription initiation ^3,14,27^. An alternative possibility is that TF binding to G4 regions stimulates G4 folding to maintain the underneath chromatin in a transcription permissive state.

**Figure 4.**
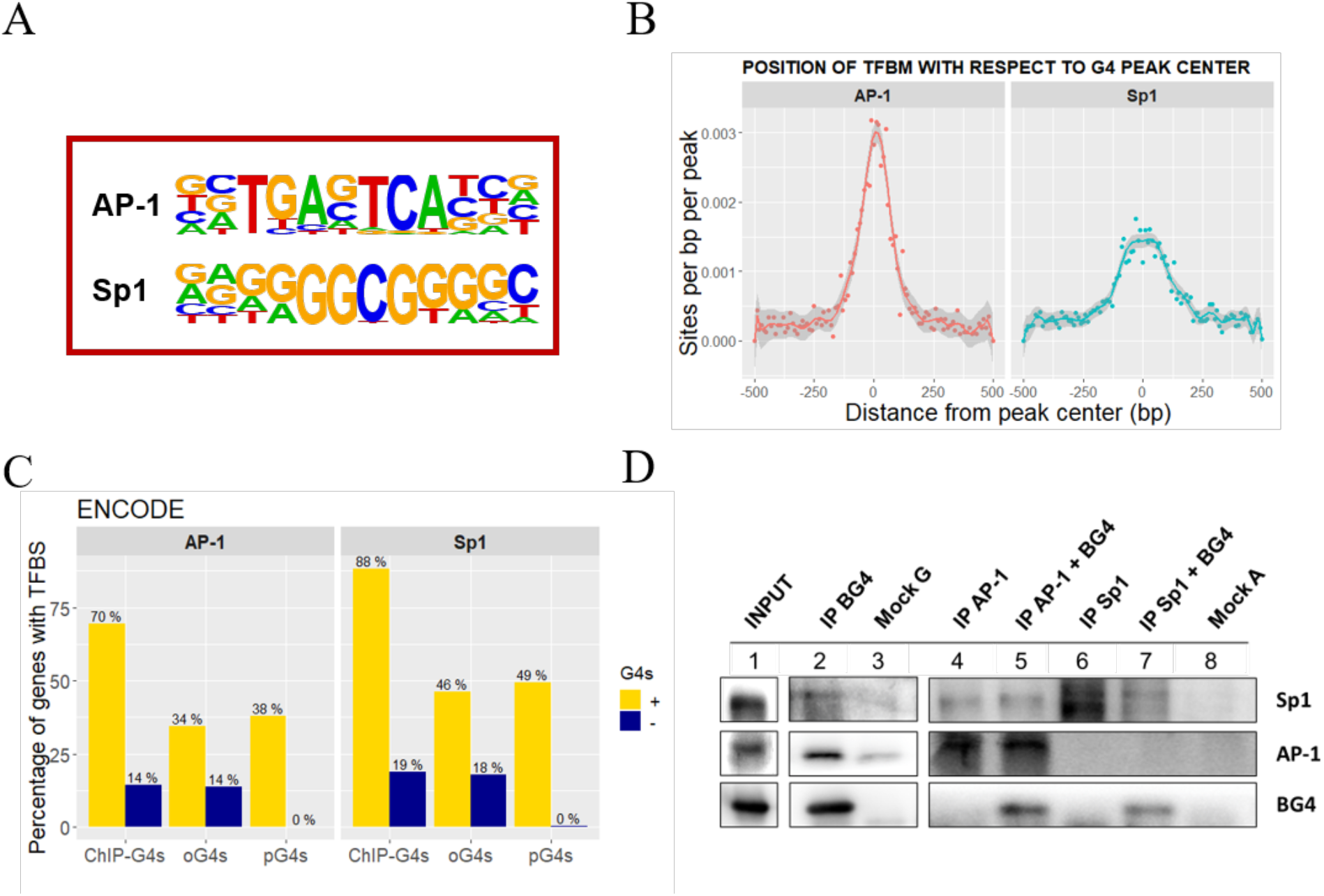
Identification of TFs binding to BG4-IP regions in 93T449 cells. **A)** Consensus sequences of TFs binding sites that are significantly enriched in BG4 ChIP peaks, as calculated by Homer software. **B)** Position and frequency of TF binding sites with respect to the BG4 ChIP peak centre. **C)** Percentage of genes with and without ChIP-G4s, oG4s and pG4s containing validated TFBS for AP-1 and SP1 according to ENCODE database. **D)** Western blot showing co-immunoprecipitation of G4s and the two TFs SP1 and AP-1. The INPUT lane (1) corresponds to the total fraction of the sheared chromatin used as starting material. G4s were immunoprecipitated by BG4 antibody (IP BG4) and AP-1 and SP1 were detected by immunoblotting (lane 2). AP-1 (lanes 4 and 5) and SP1 (lanes 6-7) TFs were immunoprecipitated from the sheared chromatin with or without previous incubation in the presence of BG4 antibody. Mock G (lane 3) and A (lane 8) are the negative controls immunoprecipitated in the absence of primary antibody using protein G or protein A coated beads, respectively.

Our data also indicate that the use of an anti-G4 antibody is a powerful tool to investigate the G4 landscape in cells. There have been doubts about the possible induction of G4 folding by this approach. However, our data help dissipate this doubt since G4s have been differently detected in the same sequence in different cell types, fact that indicates the actual recognition of a previously formed G4 rather than G4 induction.

Overall our results point to G4s as key epigenetic regulatory elements that, in association with transcription factors, activate gene expression.

## Materials and methods

### Primers and compounds

Desalted primers were purchased from Eurofins (Munich, Germany). A detailed list of primers name and sequence is available in the **Supplementary Table 1**.

### BG4 expression and purification

BG4-encoding plasmid (kindly provided by Professor Shankar Balasubramanian, University of Cambridge, UK) was transformed into BL21(DE3) competent cells (Stratagene) which were cultured in TY medium (1.6% tryptone peptone, 1% yeast extract and 0.5% NaCl) and 50 μg/ml kanamycin. Transformed cells were grown at 37°C 160 rpm to an OD_600_ of 0.7-0.8. BG4 antibody expression was induced with 0.85 mM isopropyl β-D-1-thiogalactopyranoside overnight at RT. The cells were pelleted for 25 min at 25,000g at 4 °C, resuspended in lysis buffer (20 mM Tris-Cl pH 8.0, 50 mM NaCl, 5% Glycerol, 1% Triton and 100 μM Phenylmethanesulfonylfluoride solution) and lysed through 5 cycles of freezing and thawing. After centrifugation at 10,000g at 4 °C for 20 min, the supernatant was filtered (0.45 μm) and purified on a Protino Ni-NTA-Agarose Affinity column (Machery-Nagel, Germany) according to the manufacturer instructions. The column was washed in 20 mM imidazole in 20 mM Tris HCl pH 8.0 and 300 mM NaCl, and BG4 antibody 1.5 ml fractions eluted in 250 mM imidazole in 20 mM Tris HCl pH 8.0 and 300 mM NaCl. BG4 antibody containing fractions were checked on a Coomassie-stained SDS-PAGE and concentrated in Amicon Ultra-3k Centrifugal Filter Unit (Millipore). The concentration of BG4 was determined using Thermo Scientific Pierce BCA Protein Assay kit, and the antibody was stored at −20 °C.

### G4 chromatin immunoprecipitation

The 93T449 (ATCC^®^ CRL-3043™) cells were grown to 80 % confluence in RPMI 1640 (Gibco, Thermo Fisher Scientific, Waltham, MA, USA) supplemented with 10% heat inactivated FBS. After trypsinization, 2 million cells were fixed in RPMI containing 1 % (v/v) formaldehyde and 10 % (v/v) FBS for 10 min at RT. After 5 min quenching with 125 mM glycine, cells were pelleted and washed twice with PBS containing 10 % FBS. The flash frozen pellets were lysed for 5 min on ice in 100 μl of 50 mM tris-HCl pH 8.0, 10 mM EDTA, 0.5 % SDS and protease inhibitor cocktail. Samples were sonicated using the Covaris E220 to shear chromatin to an average size of 100-500 bp (2 % duty cycle, 105 W peak incident power, 200 cycles per burst, 25 min). Sheared chromatin was diluted 1:5 in IP-buffer (10 mM Tris-HCl pH 7.5, 1mM EDTA, 0.5 mM EGTA, 1% Triton X-100, 0.1 % SDS, 0.1 % Na-deoxycholate and 140 mM NaCl) supplemented with protease inhibitor cocktail. After centrifuging 10 min at 13,000g at 4 °C, the supernatant containing soluble chromatin fraction was recovered and incubated with 0.7 mg/ml RNase A (ThermoFisher) for 30 min at 37 °C. For chromatin immunoprecipitation 10 μl protein-G magnetic beads (Pierce™ ThermoFisher) were washed in IP-buffer and incubated with 1 μg Anti-FLAG Ab (Sigma-Aldrich #F3165) for 1 h at 4 °C on a rotating wheel. 50 μl of RNA digested chromatin were incubated with 250 ng BG4 Ab (or without for the mock negative control) for 1h at 16 °C. The anti-FLAG coated beads were washed with IP-buffer and incubated with chromatin-BG4 complex for 3 h at 4 °C on a rotating wheel. Beads were washed 4 times with IP-buffer and once in wash buffer (10 mM Tris-HCl pH 8.0, 10 mM EDTA). Elution of immunoprecipitates and chromatin crosslink reversal were performed incubating beads with 70 μl elution buffer (10 mM Tris-HCl pH 8.0, 5 mM EDTA, 300 mM NaCl and 0.5 % SDS) containing 0.3 mg/ml RNase A (Thermo Fisher) for 30 min at 37 °C followed by the addition of 0.5 mg/ml proteinase K for 1h at 55 °C and 8 h at 65 °C shaking. Supernatant was then recovered and incubated for 1 additional h at 55 °C in the presence of 0.25 mg/ml proteinase K (Thermo Fisher). The eluate was finally purified with SPRI AMPure XP beads (Backman Coulter). For each technical replicate, eluted DNA from two ChIP reactions were combined and the pool subjected either to G4 enrichment quantification via qPCR or to to library preparation for sequencing.

### G-quadruplex ChIP-qPCR

The immunoprecipitated sample (IP and Mock) and the input were used to quantify G4 enrichment via qPCR, using Fast SYBR PCR mix (Applied Biosystems), with a LightCycler 480 (Roche) quantitative PCR machine. Cycling conditions were 95 °C for 20 s followed by 50 cycles of 3 s at 95 °C and 30 s at 60 °C. We employed primer pairs that target G4 ChIP positive and negative regions **(Supplementary Table 1)**. Relative enrichments were derived with respect to their inputs.

### G4 ChIP-seq library preparation

The immunoprecipitated sample and the input were subjected to Nextera library preparation as described by the manufacturer (NEBNext Ultra II DNA library Prep Kit for Illumina, NEB). The quality and size of libraries and chromatin shearing fragments were checked by Agilent Bioanalyzer using Agilent DNA High Sensitivity Chips (Agilent Technologies). Samples were sequenced on an Illumina HiSeq 1500 platform in single-end using 50-bp reads. The experiment was repeated twice.

### Putative G4s prediction

The presence of pG4s in the BG4 immunoprecipitated peaks was assessed by two different computational tools: i) Quadparser, based on a regular expression matching algorithm ^19^; ii) G4Hunter, based on a scored sliding window approach that considers also the G-richness and skewness of all bases in each window ^20,28^. A fasta file containing the sequences of BG4 immunoprecipitated regions was obtained from HOMER output by mean of bedtools ^29^ and used as input of both prediction algorithms. Quadparser script was downloaded from https://github.com/dariober/ as indicated by Puig Lombardi et al. ^28^, and applied with two different regular expressions, in order to allow the matching of loops with length 0-7 ([gG]{2,5}\w{0,7}){3,}[gG]{2,5} or 0-12 ([gG]{2,5}\w{0,12}){3,}[gG]{2,5}. G4Hunter algorithm was instead retrieved from https://github.com/AnimaTardeb/G4Hunter and applied using a window size of 15 bp and score threshold of 1.25, demonstrated to reliably discriminate G4s from non-G4s sequences ^28^. The obtained results were then evaluated by using R programming language.

### Cells doubling time

For the calculation of cells doubling time (DT), 93T449 and HaCaT were seeded at a density of 40000 cells/ml and 20000 cells/ml respectively in multiwell plates. The cells density was then monitored every 24 hours by counting cells in Trypan Blue (Sigma Aldrich) using a Bürker chamber, until the cells growth curve reached the plateau. The DT was then calculated as DT=T ln2/ln(Xe/Xb), where T is the incubation time, Xb is the cells number at the beginning of the incubation time and Xe is the cell number at the end of the incubation time.

### RNA extraction and cDNA library preparation

Total RNA for RNA-seq experiments was extracted from 80% confluent cells. Then, 1 million cells were trypsinized, pelleted and resuspended in 1 ml TRIreagent (Sigma Aldrich). Total RNA was extracted by phenolchloroform and purified through the RNA Clean and Concentrator-25 kit (Zymo Research), following the manufacturer’s instructions. Ribosomal RNA was depleted and purified using Ribo-Zero rRNARemoval Kit (Illumina) and RNA Clean and Concentrator-5 (Zymo Research). The quality of extracted RNA was checked by Agilent Bioanalyzer on Agilent RNA 6000 Pico Chips (Agilent technologies) both before and after rRNA depletion. RNA-seq libraries were generated using the NEBNext Ultra Directional RNA Library Prep kit for Illumina (NEB). Agilent DNA High Sensitivity Chips (Agilent Technologies) were used to check library size and quality. Samples were sequenced on an Illumina HiSeq 1500 platform in single-end using 50-bp reads. The experiment was done in three independent biological replicates.

### Comparison of pG4s, oG4s and ChIP-G4s

To obtain direct evidence of the association between G4s in the folded state and transcriptional level, three G4 categories were compared, namely: pG4s, oG4s and ChIP-G4s. This comparison was used to distinguish the transcriptional effect of a G4 permissive sequence versus the folded G4 structure. pG4s correspond to the Quadparser ^19^ predicted G4s with G-tracts of 2-5 Gs and loops length 0-12 nts; oG4s are G4s forming sequences detected by modified Illumina sequencing protocol from Chambers et al. ^23,30^; ChIP-G4s are the BG4 immunoprecipitated regions in 93T449 cells. Bed files containing the G4 regions of each category were annotated using HOMER software to retrieve the corresponding gene and then integrated with 93T449 RNA-seq data to evaluate their expression level.

### ATAC-seq (Assay for Transposase Accessible Chromatin followed by high throughput sequencing)

ATAC-seq of 93T449 cells was performed according to the Omni-ATAC protocol developed by Corces et al. 2017 ^24^. Briefly, 50000 viable cells were pelleted and lysed in Resuspension buffer (Tris-HCl pH 7.4 10 mM, NaCl 10 mM, MgCl_2_ 3 mM, NP40 0.1%, Tween-20 0.1%, digitonin 0.01% - Promega #G9441) for 3 min on ice. Next nuclei were pelleted and the transposition reaction was performed incubating the lysate for 30 min at 37 °C under agitation in the presence of Transposition mixture (Tris-HCl pH 7.6 10 mM, MgCl_2_ 5 mM, dimethyl formamide 10%, Tn5 enzyme 100 nM – Illumina #20018704, digitonin 0.01% - Promega #G9441, Tween-20 0.1%, PBS 33%). The transposition reaction was purified (Qiagen MinElute PCR Purification kit, Qiagen #28004) and pre-amplified by PCR in the presence of adapter primers (NEBNext 2x PCR Master Mix, New England Biolabs Inc. #M0541S). Amplification was monitored by RT-PCR using the PowerUp™ SYBR™ Green Master Mix (ThermoFisher Scientific, #A25741) to determine the number of additional amplification cycles ^31^. Finally, samples were purified with SPRI AMPure XP beads (ThermoFisher Scientific) and sequenced on an Illumina HiSeq 1500 platform in single-end using 50-bp reads.

### Identification of G4-associated transcription factors binding sites

Bed files of 93T449 ChIP-G4s, Quadparser predicted pG4s (G 2-5, loop 0-12) and oG4s from Chamber et al. ^23^ were used to annotate peaks and extract genes with promoter G4s by mean of HOMER software. The complete list of human genes annotated on the GRCh38-hg38 reference genome was retrieved from BioMart Ensembl database (http://www.ensembl.org/biomart/martview) and used to divide genes into promoter G4 containing/depleted genes according to the three categories of ChIP-G4s (**Supplementary Files 8.1** and **8.2** for known motifs, **Supplementary Files 8.1.1** and **8.2.1** for de novo motifs), pG4s and oG4s. The findMotifs.pl function of HOMER software was employed to predict the presence of TFBSs associated to either all the ChIP-G4s peaks and genes with/without promoter ChIP-G4s, pG4s and oG4s. Since promoter ChIP-G4s were strongly clustered within −1000 and +750 bp form the corresponding gene TSS we set this region for the prediction of TFBSs. To confirm G4 association with the predicted TFs, the validated target genes of AP-1 and Sp1 were retrieved from ENCODE database, accessed through Harmonizome ^32^.

### Co-immunoprecipitation of G4s and transcription factors and Western blotting

Co-immunoprecipitations to detect G4s interaction with TFs were performed by immunoprecipitating either BG4-G4complex, or AP-1 and Sp1 TFs. In the first case G4s immunoprecipitation was performed as described for the BG4 ChIP-qPCR procedure, but after the last wash, three replicates of BG4 captured material were collected and eluted in 35 μl Elution buffer (10 mM Tris-HCl pH 8.0, 5 mM EDTA, 300 mM NaCl and 0.5 % SDS) by incubating beads for 1 h at 55 °C to revert the cross-linking without degrading proteins. The collected proteins are expected to be G4s interacting proteins, since they were co-immunoprecipitated together with BG4. The presence of BG4 (internal positive control) and the TFs AP-1 and Sp1 was then checked by Western Blot as described below.

For the immunoprecipitation of AP-1 and Sp1, the fixed and sheared chromatin from 1.5 million cells was treated with 0.7 mg/ml RNase A (ThermoFisher) for 30 min at 37 °C, precleared with protein-A magnetic beads (Pierce™ ThermoFisher) for 30 min at 4°C under rotation to reduce the background due to the non specific adhesion of sample to the beads. 20 μl protein-A magnetic beads (Pierce™ ThermoFisher) were washed in IP-buffer and incubated with 4 μg anti-AP-1 (Thermo Scientific™ #MA5-15172)or anti-Sp1 Ab (ChIPAb+™ Merck #17-601) for 1 h at 4 °C on a rotating wheel. After incubation in the presence or absence of BG4 for 1h at 16 °C, the pre-cleared chromatin was incubated on the AP-1, Sp1 functionalized beads, or non functionalized protein-A beads as Mock for 4 h at 4 °C under rotation, washed 4 times with IP-buffer and once in Wash buffer (10 mM Tris-HCl pH 8.0, 10 mM EDTA). After the last wash, three replicates were collected and eluted in 35 μl Elution buffer (10 mM Tris-HCl pH 8.0, 5 mM EDTA, 300 mM NaCl and 0.5 % SDS) by incubating beads for 1 h at 55 °C to revert the cross-linking without degrading proteins. If AP-1 and Sp1 TFs interact with G4s, BG4 antibody is supposed to be co-immunoprecipitated. Thus, Western Blotting was employed to detect AP-1, Sp1 (internal positive controls) and BG4 in the immunoprecipitated samples.

For Western Blotting of both co-immunoprecapitation approaches, the eluted proteins were quantified by Thermo Scientific Pierce BCA Protein Assay kit-The INPUT, IP and Mock (negative control immunoprecipitated without BG4) were next loaded on a SDS-PAGE denaturing gel. The gel separated proteins were transferred on a PVDF membrane, blocked in TBS-tween 0.1% buffer supplemented with 5% BSA, incubated with primary antibodies (anti-AP-1 Thermo Scientific™ #MA5-15172, anti-Sp1 ChIPAb+™ Merck #17-601, anti-FLAG Sigma Aldrich #F3165, anti-NCL Santa Cruz Biotechnology #sc-803), washed in TBS-tween 0.1%, next incubated with secondary goat anti-rabbit and goat anti-mouse HRP antibody (Millipore #). Images were acquired on a Uvitec instrument by reading HRP bioluminescence.

### Data analysis

Raw FASTQ reads were trimmed to remove adaptor contamination and aligned to the primary assembly of the human reference genome version GRCh38 using Bowtie1 (http://bowtie-bio.sourceforge.net/index.shtml). Reads with more than 2 mismatches and multimapped reads were excluded from further analysis.

G4-ChIP peaks were identified and mapped using HOMER (http://homer.ucsd.edu/homer/index.html). Only peaks with at least 2 fold more normalized tags count in the target experiment with respect to the input (used as control) were considered, the analysis was performed with disabled local tag count and poisson p-value threshold of 0.0001. HOMER was next used to associate peaks with the nearby gene, determine the genomic annotation of the region occupied by the peak and merge peaks from replicates.

RNA-seq reads were aligned to the human reference genome with TopHat and filtered by using samtools ^33^ to remove alignments with quality lower than 20, not primary alignments and PCR duplicates. Gene expression levels were quantified as transcripts per million (TPM). Genes differentially expressed between 93T449 cells and HaCaT cells and (s-value < 0.1; fold change > 1.0) were identified using the Bioconductor package DESeq2 and ‘apeglm’ for LFC shrinkage ^25^.

ATAC-seq peaks were identified and mapped using HOMER and employing the ChIP-seq input sample as genomic reference. A fixed peak size was estimated by the software on the basis of autocorrelation analysis. The other parameters were set todefault. Bedtools were used to merge HOMER peak files into a single bed file ^29^. HOMER was next used to associate peaks with the nearby gene and determine the genomic annotation of the region occupied by the peak. For the intersection of ChIP-seq and ATAC-seq peaks the bedtools intersect function was employed.

All the further statistical analysis was performed using R ^34^.

## DATA AVAILABILITY

All genomic data produced in the present project (93T449 G4-ChIP-seq, ATAC-seq and RNA-seq) have been deposited in the NCBI GEO database under accession number GSE145543 (reviewer token: wzwvciymxvqrlal).

HaCaT cells datasets for G4-ChIP-seq, ATAC-seq and RNA-seq were instead downloaded from GEO at the following accession number GSE76688.

## Notes

### Competing Interest Statement

The authors have declared no competing interest.

## Reference

1. D’Alessio, A. C. et al. A systematic approach to identify candidate transcription factors that control cell identity. Stem Cell Reports 5, 763–775 (2015).

2. Natarajan, A., Yardimci, G. G., Sheffield, N. C., Crawford, G. E. & Ohler, U. Predicting cell-type- specific gene expression from regions of open chromatin. Genome Res. 22, 1711–1722 (2012).

3. Varizhuk, A., Isaakova, E. & Pozmogova, G. DNA G-Quadruplexes (G4s) Modulate Epigenetic (Re)Programming and Chromatin Remodeling. BioEssays 41, 1900091 (2019).

4. Hänsel-Hertsch, R. et al. G-quadruplex structures mark human regulatory chromatin. Nat. Genet. 48, 1267–1272 (2016).

5. Brooks, T. A. & Hurley, L. H. Targeting MYC Expression through G-Quadruplexes. Genes Cancer 1, 641–649 (2010).

6. Siddiqui-Jain, A., Grand, C. L., Bearss, D. J. & Hurley, L. H. Direct evidence for a G-quadruplex in a promoter region and its targeting with a small molecule to repress c-MYC transcription. Proc. Natl. Acad. Sci. 99, 11593–11598 (2002).

7. Cogoi, S. & Xodo, L. E. G-quadruplex formation within the promoter of the KRAS proto-oncogene and its effect on transcription. Nucleic Acids Res. 34, 2536–2549 (2006).

8. Recagni et al. The Oncogenic Signaling Pathways in BRAF-Mutant Melanoma Cells are Modulated by Naphthalene Diimide-Like G-Quadruplex Ligands. Cells 8, 1274 (2019).

9. Tosoni, E. et al. Nucleolin stabilizes G-quadruplex structures folded by the LTR promoter and silences HIV-1 viral transcription. Nucleic Acids Res. 43, 8884–8897 (2015).

10. Kim, N. The Interplay between G-quadruplex and Transcription. Curr. Med. Chem. 26, 2898–2917 (2017).

11. Raiber, E.-A., Kranaster, R., Lam, E., Nikan, M. & Balasubramanian, S. A non-canonical DNA structure is a binding motif for the transcription factor SP1 in vitro. Nucleic Acids Res. 40, 1499–1508 (2012).

12. Cogoi, S., Paramasivam, M., Membrino, A., Yokoyama, K. K. & Xodo, L. E. The KRAS promoter responds to Myc-associated zinc finger and poly(ADP-ribose) polymerase 1 proteins, which recognize a critical quadruplex-forming GA-element. J. Biol. Chem. 285, 22003–22016 (2010).

13. Uribe, D. J., Guo, K., Shin, Y. J. & Sun, D. Heterogeneous nuclear ribonucleoprotein K and nucleolin as transcriptional activators of the vascular endothelial growth factor promoter through interaction with secondary DNA structures. Biochemistry 50, 3796–3806 (2011).

14. Mao, S. Q. et al. DNA G-quadruplex structures mold the DNA methylome. Nat. Struct. Mol. Biol. 25, 951–957 (2018).

15. Hou, Y. et al. Integrative characterization of G-Quadruplexes in the three-dimensional chromatin structure. Epigenetics 14, 894–911 (2019).

16. Du, Z., Zhao, Y. & Li, N. Genome-wide analysis reveals regulatory role of G4 DNA in gene transcription. Genome Res. 18, 233–241 (2008).

17. Biffi, G., Tannahill, D., McCafferty, J. & Balasubramanian, S. Quantitative visualization of DNA G- quadruplex structures in human cells. Nat. Chem. 5, 182–186 (2013).

18. Kango-Singh, M. Vogel and Motulsky’s human genetics--problems and approaches. Hum. Genomics 5, 73 (2010).

19. Huppert, J. L. & Balasubramanian, S. Prevalence of quadruplexes in the human genome. Nucleic Acids Res. 33, 2908–2916 (2005).

20. Bedrat, A., Lacroix, L. & Mergny, J. L. Re-evaluation of G-quadruplex propensity with G4Hunter. Nucleic Acids Res. 44, 1746–1759 (2016).

21. Balasubramanian, S., Hurley, L. H. & Neidle, S. Targeting G-quadruplexes in gene promoters: A novel anticancer strategy? Nat. Rev. Drug Discov. 10, 261–275 (2011).

22. Neidle, S. Quadruplex Nucleic Acids as Novel Therapeutic Targets. J. Med. Chem. 59, 5987–6011 (2016).

23. Chambers, V. S. et al. High-throughput sequencing of DNA G-quadruplex structures in the human genome. Nat. Biotechnol. 33, 877–881 (2015).

24. Corces, M. R. et al. An improved ATAC-seq protocol reduces background and enables interrogation of frozen tissues. Nat. Methods 14, 959–962 (2017).

25. Zhu, A., Ibrahim, J. G. & Love, M. I. Heavy-tailed prior distributions for sequence count data: removing the noise and preserving large differences. Bioinformatics 35, 2084–2092 (2019).

26. Hänsel-Hertsch, R. et al. Landscape of G-quadruplex DNA structural regions in breast cancer. Nat. Genet. (2020) doi:10.1038/s41588-020-0672-8.

27. Jara-Espejo, M. & Peres Line, S. R. DNA G-quadruplex stability, position and chromatin accessibility are associated with CpG island methylation. FEBS J. (2019) doi:10.1111/febs.15065.

28. Puig Lombardi, E. & Londoño-Vallejo, A. A guide to computational methods for G-quadruplex prediction. Nucleic Acids Res. (2019) doi:10.1093/nar/gkz1097.

29. Quinlan, A. R. & Hall, I. M. BEDTools: A flexible suite of utilities for comparing genomic features. Bioinformatics 26, 841–842 (2010).

30. Marsico, G. et al. Whole genome experimental maps of DNA G-quadruplexes in multiple species. Nucleic Acids Res. 47, 3862–3874 (2019).

31. Buenrostro, J. D., Wu, B., Chang, H. Y. & Greenleaf, W. J. ATAC-seq: A method for assaying chromatin accessibility genome-wide. Curr. Protoc. Mol. Biol. 2015, 21.29.1–21.29.9 (2015).

32. Rouillard, A. D. et al. The harmonizome: a collection of processed datasets gathered to serve and mine knowledge about genes and proteins. Database (Oxford). 2016, (2016).

33. Li, H. et al. The Sequence Alignment/Map format and SAMtools. Bioinformatics 25, 2078–9 (2009).

34. R Core Team. R: A Language and Environment for Statistical Computing. (2013).

