## Supplementary Information for "Promoter G-quadruplexes and transcription factors cooperate to shape the cell type-specific transcriptome"

**Supplementary Table 1. Comparison of pG4s found by Quadparser and G4Hunter**

| TOOL | G4nr | + Strand | - Strand | Mean length (bp) |
|---|---|---|---|---|
| ***Quadparser 0-7*** | 7414 | 3597 | 3817 | 24 |
| ***Quadparser 0-12*** | 10053 | 4981 | 5072 | 36 |
| ***G4Hunter*** | 9700 | 4658 | 5042 | 21 |

Absolute number, strand localization and mean length of the pG4s found in the BG4 immunoprecipitated peaks according to the prediction tools Quadparser (loop length 0-7 and 0-12) and G4Hunter (window size 15, score threshold 1.25).

**Supplementary Table 2**. **List of primers used in qPCR after BG4-ChIP to test for G4 enrichment in the immunoprecipitated samples.**

| **Primer** | **Sequence** |
|---|---|
| *GAPDH* | FW: GCTACTAGCGGTTTTACGGGCG – RV: TGCGGCTGACTGTCGAACAGG |
| *EIF4A1* | FW: CCGGAGCGACTAGGAACTAAC – RV: GCCTTTCTTACCGGGAATCCT |
| *MDM2* | FW: GGATTTCGGACGGCTCTCG - RV: CGTTCACACTAGTGACCCGA |
| *CDK4* | FW: CCACCCTCACCATGTGACC - RV: CTTACACTCTTCGCCCTCCTC |
| *TMCC2* | FW: CCAGACACTTTGGGTGACCT – RV: AACACCTGCTCTGCCAACTT |
| *MAP3K13* | FW: GACATAGGAACGGGCAAAGA – RV: CCCATGCTGTATGTGGTCTG |
| *ESR1* | FW: GAAACAGCCCCAAATCTCAA – RV: TTGTAGCCAGCAAGCAAATG |
| *LRRN4* | FW: GAGGCTGGGATCTCAGTGTTCGG – RV: TACTCTCTGAACCAAGGGGCACT |

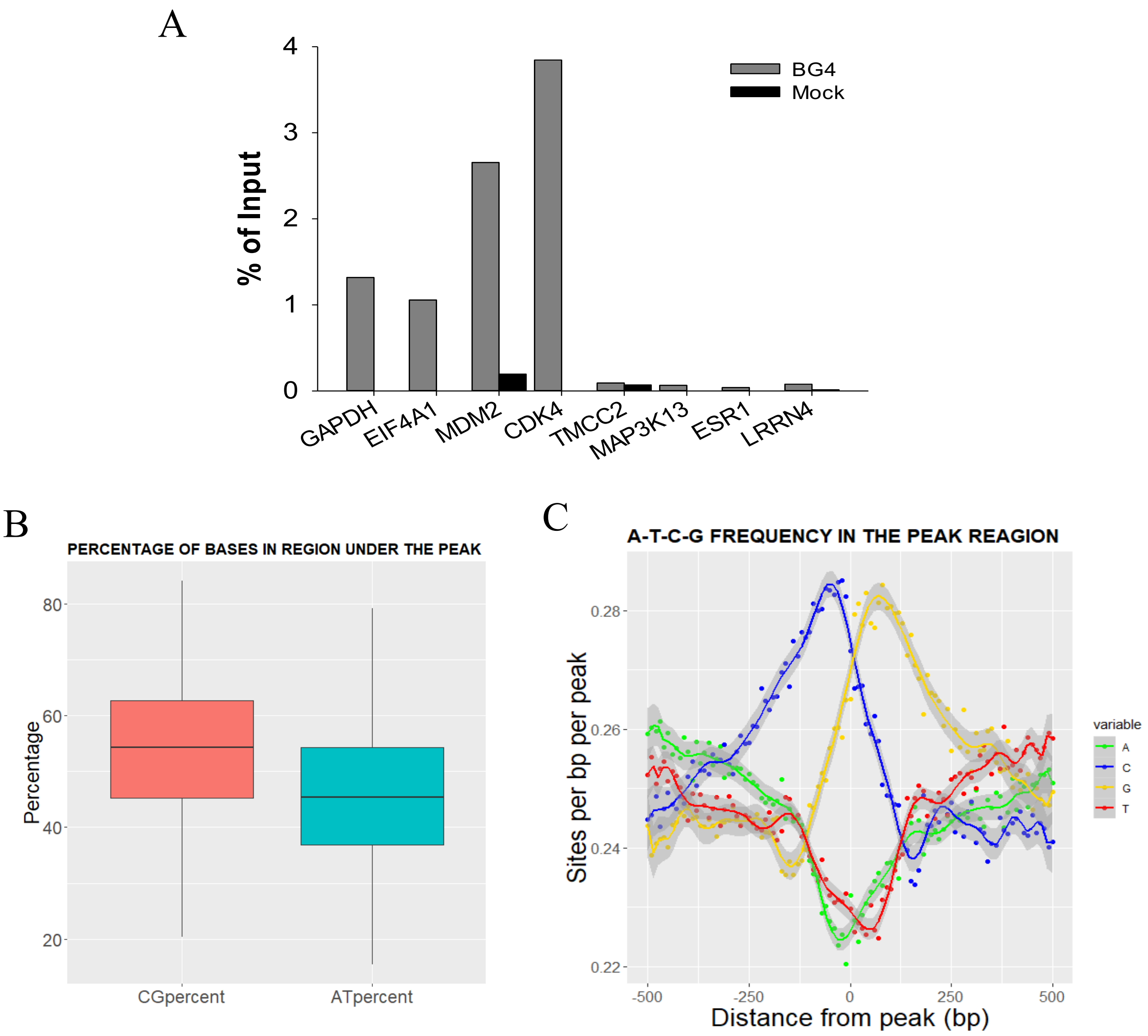


**Supplementary Figure 1.** ***Control parameters in BG4 ChIP-seq analysis*****. A)** Control of G4 enrichment in immunoprecipitated chromatin. ChIP-qPCR on G4-positive (GAPDH, EIF4A, MDM2, CDK4) and G4-negative (TMCC2, MAP3K13, ESR1, LRRN4) control regions. Reliable G4 enrichment was observed in *GAPDH*, *EIF4A1*, *MDM2* and *CDK4* gene promoters, where G4s have been previously detected[4]. These were chosen as *GAPDH* and *EIF4A* preserve a conserved expression along different cell lines[6,7], while *MDM2* and *CDK4* are highly expressed relevant oncogenes in WDLPS carcinogenesis[8,9]. In contrast, negligible G4 signal was obtained in *TMCC2*, *MAP3K13*, *ESR1* and *LRRN4* gene promoters, the sequences of which do not contain any G4s. Grey bars represent the BG4 immunoprecipitated regions, while black bars correspond to the sample immunoprecipitated in the absence of BG4 antibody. **B)** Percentage of CG and AT bases in BG4 immunoprecipitated peaks. **C)** A, T, C, G base frequency in BG4 immunoprecipitated peaks reported as function of the distance from the peak centre.

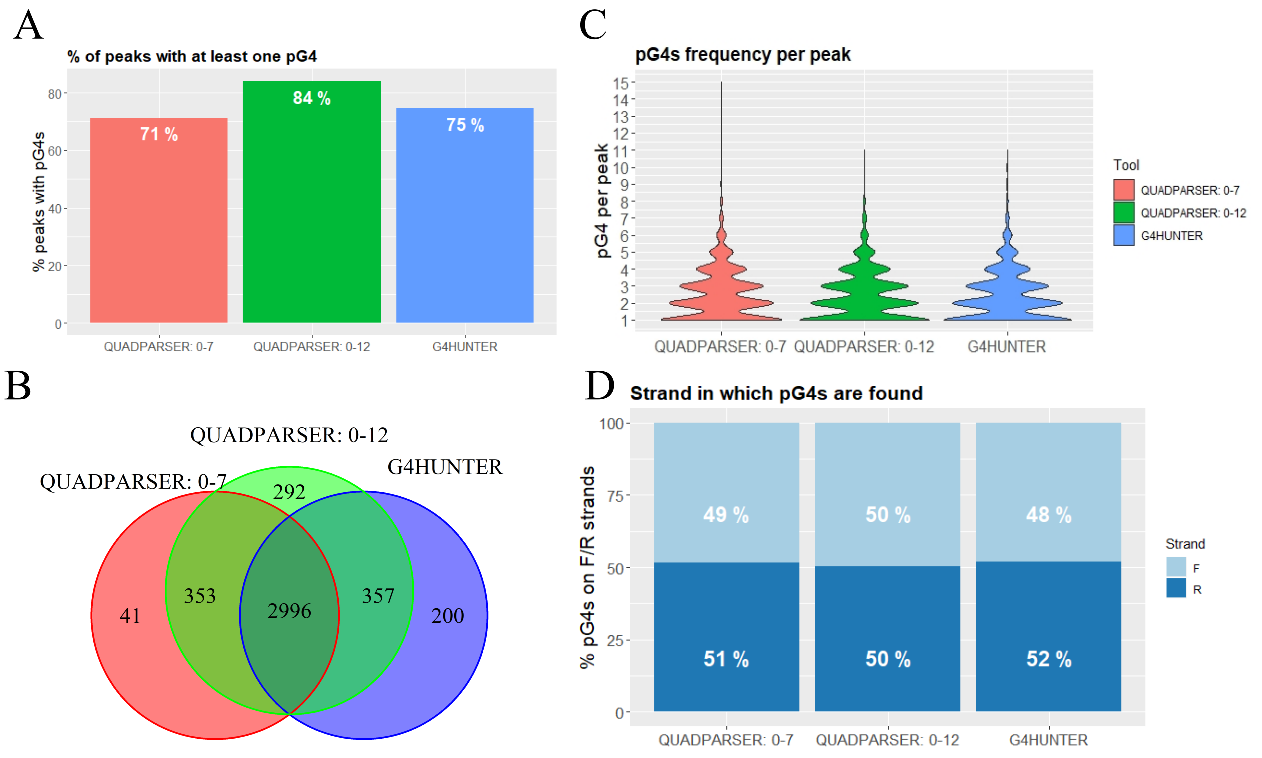


**Computational analysis of putative G4s (pG4s) in BG4 ChIP-seq peaks. A)** Percentage of peaks containing at least one pG4 according to Quadparser and G4Hunter computational prediction tools that identify canonical G4s based on a regular-expression-matching algorithm and canonical/non-canonical G4s based on a sliding window algorithm, respectively. Quadparser prediction was performed with two different settings: G4 loop-length in the range 0-7 and in the range 0-12. For G4Hunter prediction, the window size was set to 20 and the score threshold at 1.25. **B)** Venn diagram comparing the peaks in which at least one pG4 was found by Quadparser (loops length 0-7 or 0-12) and G4Hunter (window 20 and score threshold 1.25) **C)** Frequency plot of the amount of pG4s found in each ChIP-seq peak by Quadparser (loops length 0-7 or 0-12) and G4Hunter (window 20 and score threshold 1.25). **D)** Percentage of pG4s found on the forward and reverse strand according to Quadparser (loop length 0-7 or 0-12) and G4Hunter (window size 20, score threshold 1.25).

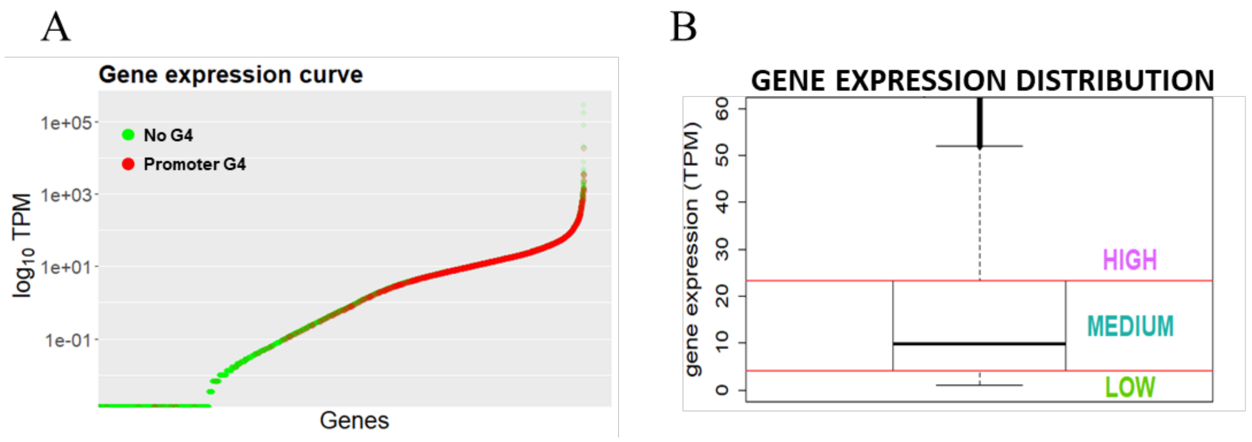


**Supplementary Figure 3.** ***G4 peaks distribution in the genome.*** **A)** A curve representing gene expression of all 93T449 genes is represented, where green dots are genes for which no promoter G4 was found and red dots are genes with G4 at their promoter. Gene expression level is reported in TPM and $\log_{10}$ scale on the y axis. **B)** Gene expression distribution of all the expressed genes (at least one transcript per gene) in 93T449 cells according to the RNA-seq data. Three expression categories were defined based on the expression distribution quartiles: the first quartile corresponded to low expression, the central two quartiles corresponded to medium expression and the upper quartile corresponded to high expression.

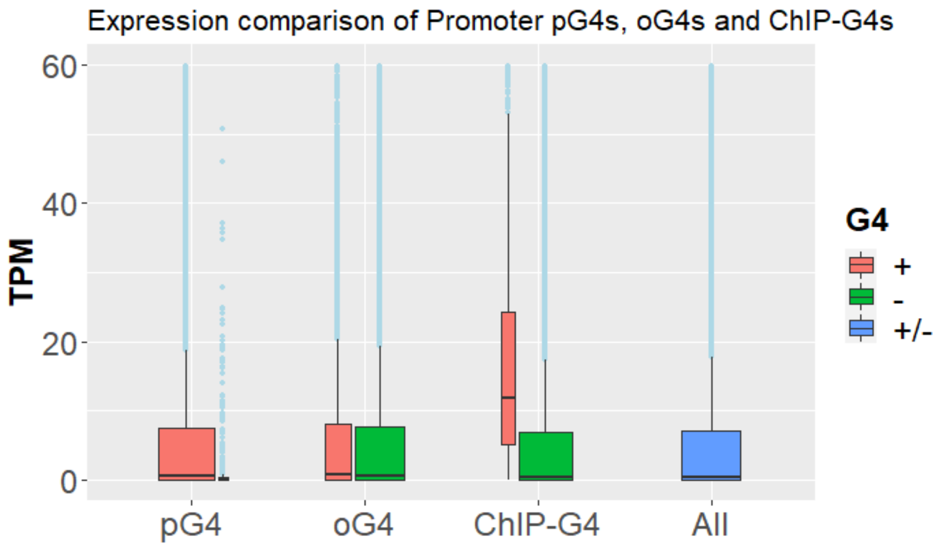


**Supplementary Figure 4.** ***Expression distribution of genes with (+) or without (-) pG4s, oG4s and ChIP-G4s compared to all genes (+/-).*** Both expressed and non-expressed (< 1 transcript per gene) were considered. Box widths are proportional to the numerosity of each category.

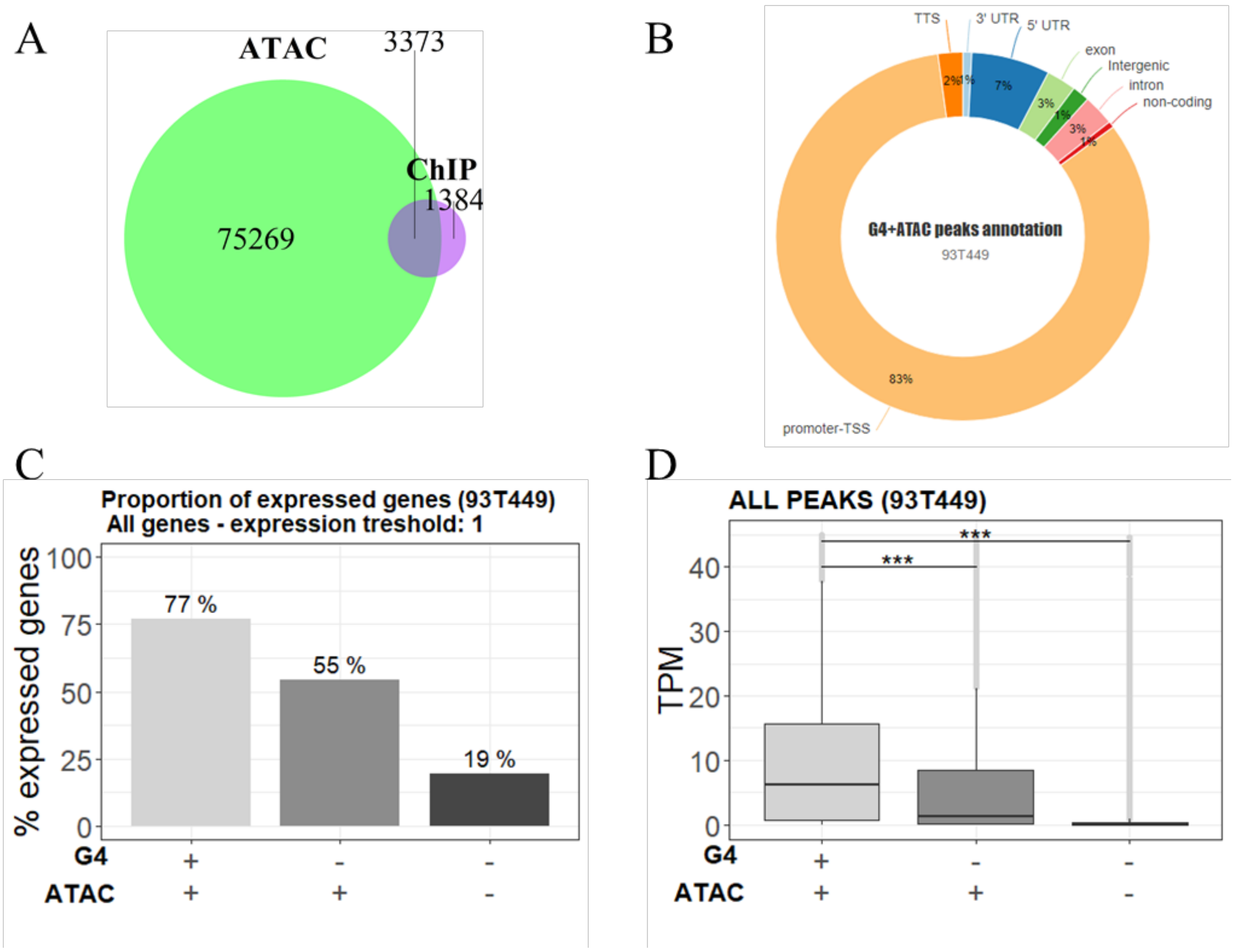


**Supplementary Figure 5.** ***Integration of G4 ChIP-seq, ATAC-seq and RNA-seq data*. A)** Venn diagram displaying the intersection between peak regions corresponding to immunoprecipitated G4s (violet) and open chromatin regions (light blue) mapped by ATAC-seq. **B)** Percentage distribution of G4 peaks that overlap with ATAC-seq peaks in functional genomic regions according to HOMER gene annotation. Percentages are normalized over the genomic abundance of each functional region. **C)** Percentage proportion of expressed genes grouped according to the presence of G4s and open chromatin signal in any functional region of the gene. Expression threshold was set to one transcript per gene. **D)** Expression distribution of all genes grouped according to the presence of G4s and open chromatin signal in any position. Expression threshold was set to 1 transcript per gene. T-test was applied to measure statistical significance (*** = pval 0.001).

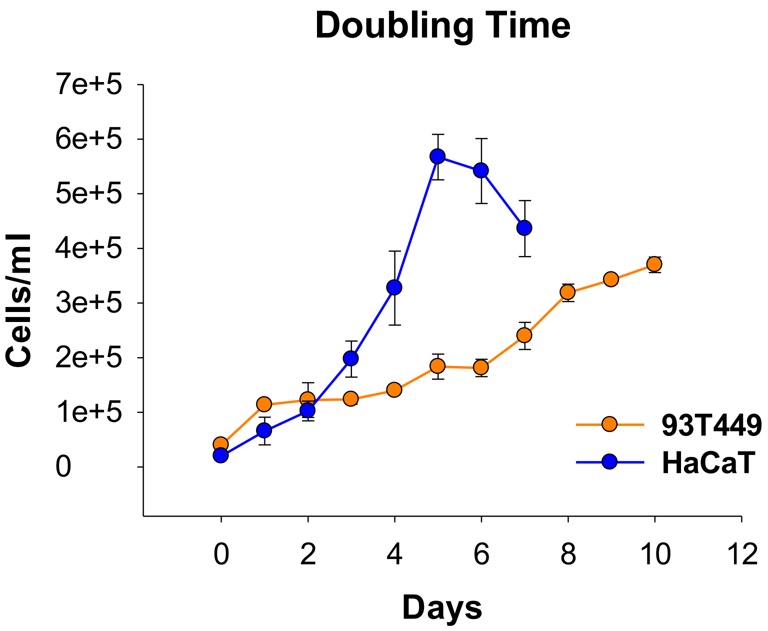


**Supplementary Figure 6.** ***Doubling time (DT) of 93T449 and HaCaT cell lines*****.** The proliferation speed of 93T449 and HaCaT cells was evaluated by monitoring the lie cells number over time. Cells number was evaluated until cells stopped growing or reach a plateau. The starting number of cells was determined based on growth curves to allow cells to proliferate for a sufficient time. Cells counts were reported in cells/ml and plotted in orange for 93T449 and blue for HaCaT. Error bars represent standard errors for three replicates.

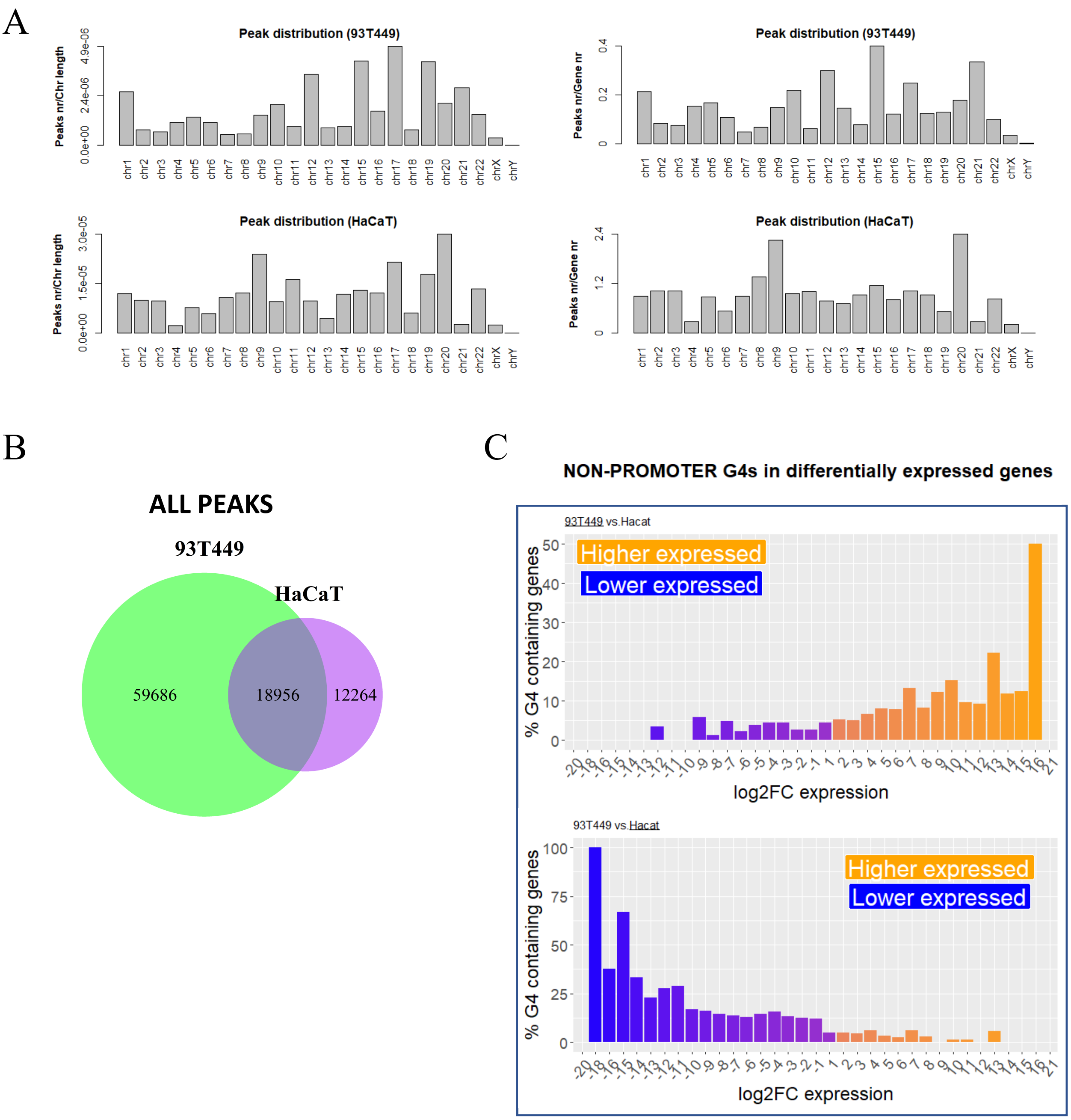


**Supplementary Figure 7. Comparison of genome-wide G4 distribution, open chromatin regions and gene expression between 93T449 and HaCaT cells. A)** Distribution of G4 peaks along chromosomes, normalized either by the chromosome length (left panels) and the number of genes encoded in each chromosome (right panels). **B)** Venn diagram showing the intersection of ATAC-seq peaks between 93T449 and HaCaT. **C)** Percentage of genes containing at least one G4 only outside of their promoter in 93T449 (upper panel) or HaCaT (lower panel) cells, in function of their differential expression in the two cell lines.

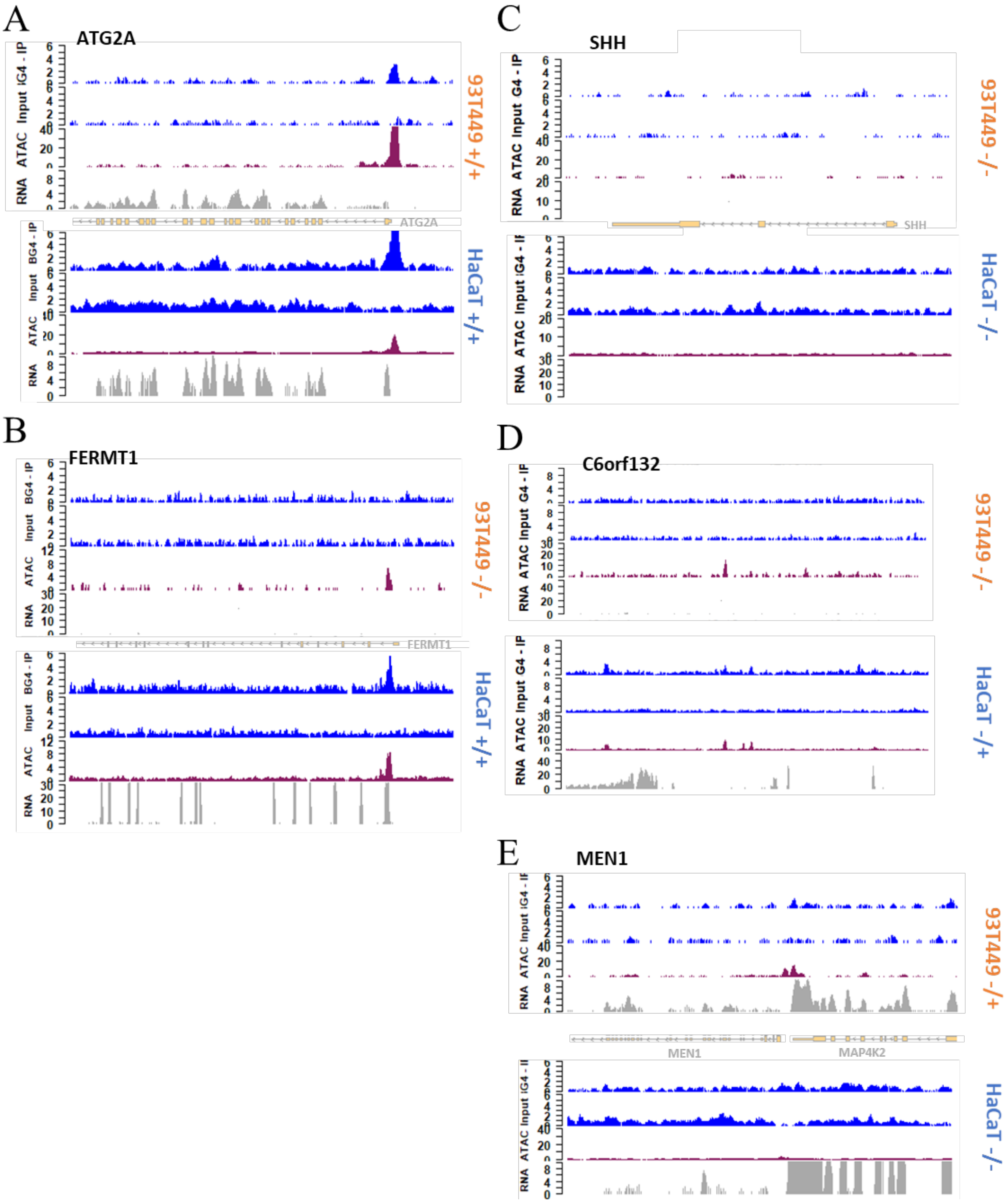


**Supplementary Figure 8. Genomic view of example regions showing how different combinations of open chromatin (ATAC) and G4 (BG4-IP) result in different transcriptional outputs (RNA) in 93T449 and HaCaT cells. A)** ATGA2 gene, with open chromatin and G4 at its promoter in both 93T449 and HaCaT cells; **B)** FERMT1 gene, with open chromatin and G4 at its promoter only in HaCaT cells; **C)** SHH gene, showing no open chromatin nor G4 in both 93T449 and HaCaT cells; **D)** C6orf132 gene, showing open chromatin but not G4 only in HaCaT cells and **E)** MEN1 gene, showing open chromatin but not G4 only in 93T449 cells.

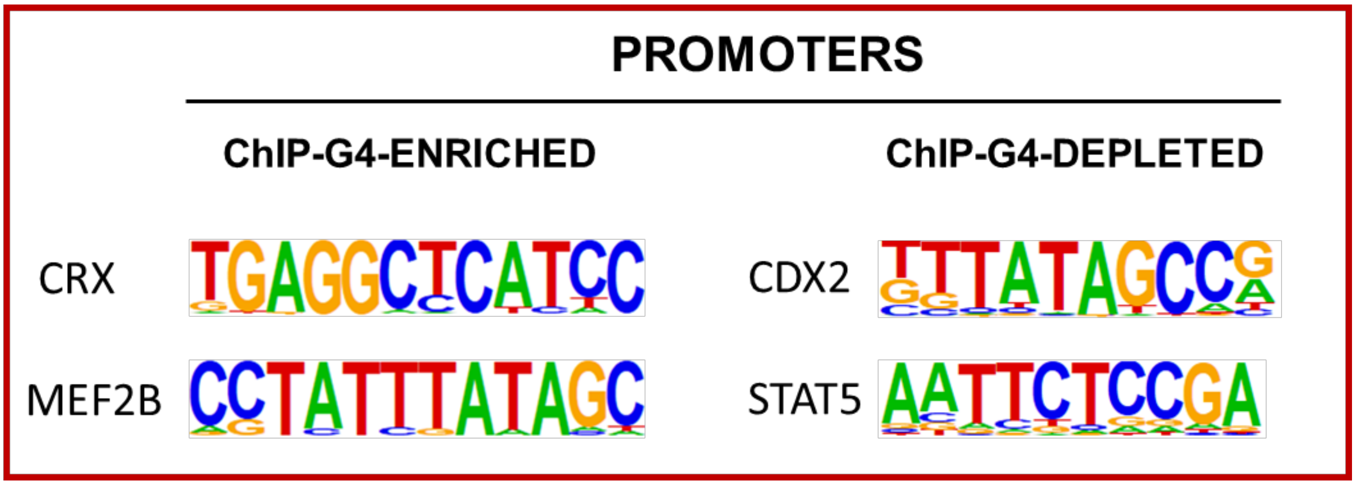


**Supplementary Figure 9. TFBS enrichment in ChIP-G4-enriched and depleted promoters.** Significantly represented motifs for TFs were calculated by HOMER software in the extended promoter region (-1000 to + 750 from TSS) of the ChIP-G4s enriched (left) or depleted (right) genes. CRX (percentage of target genes 2 %, percentage of background 0.18 %) and MEF2B (percentage of target genes 1.57 %, percentage of background 0.11 %) were identified in ChIP-G4s enriched promoters; while CDX2 (percentage of target genes 6.43 %, percentage of background 4.22 %) and STAT5 (percentage of target genes 12.80 %, percentage of background 9.77 %) were identified in promoters of ChIP-G4s depleted genes.
