## Supplementary File 8.1 for "Promoter G-quadruplexes and transcription factors cooperate to shape the cell type-specific transcriptome": SupplementaryFile8.1_GS404_HomerMotifs_knownResults.html

GS404\_IP\_BG4\_93T\_m1\_peaks\_MotifOutput/ - Homer Known Motif Enrichment Results


### Homer Known Motif Enrichment Results (GS404\_IP\_BG4\_93T\_m1\_peaks\_MotifOutput/)

Homer *de novo* Motif Results  
Gene Ontology Enrichment Results  
Known Motif Enrichment Results (txt file)  
Total Target Sequences = 2763, Total Background Sequences = 43729

|  |  |  |  |  |  |  |  |  |  |  |  |
| --- | --- | --- | --- | --- | --- | --- | --- | --- | --- | --- | --- |
| Rank | Motif | Name | P-value | log P-pvalue | q-value (Benjamini) | # Target Sequences with Motif | % of Targets Sequences with Motif | # Background Sequences with Motif | % of Background Sequences with Motif | Motif File | SVG |
| 1 | A C T G C T A G T C G A C G A T C A T G G C T A A T C G C G A T G T A C G C T A A G C T G T A C | Fra1(bZIP)/BT549-Fra1-ChIP-Seq(GSE46166)/Homer | 1e-428 | -9.869e+02 | 0.0000 | 701.0 | 25.37% | 1259.4 | 2.88% | motif file (matrix) | svg |
| 2 | C A T G C T A G T C G A A C G T A C T G C G T A T A G C C G A T T G A C C G T A A G C T G A T C | Fra2(bZIP)/Striatum-Fra2-ChIP-Seq(GSE43429)/Homer | 1e-418 | -9.638e+02 | 0.0000 | 673.0 | 24.36% | 1171.2 | 2.68% | motif file (matrix) | svg |
| 3 | C T A G T C G A A C G T A C T G C G T A A T G C A C G T G T A C C G T A A G C T G A T C G T A C | Atf3(bZIP)/GBM-ATF3-ChIP-Seq(GSE33912)/Homer | 1e-396 | -9.122e+02 | 0.0000 | 740.0 | 26.78% | 1625.8 | 3.71% | motif file (matrix) | svg |
| 4 | C A G T T G C A A C G T A C T G C G T A A T C G C G A T T G A C C G T A A C G T | BATF(bZIP)/Th17-BATF-ChIP-Seq(GSE39756)/Homer | 1e-394 | -9.095e+02 | 0.0000 | 721.0 | 26.09% | 1528.9 | 3.49% | motif file (matrix) | svg |
| 5 | C T A G T C G A G C A T C A T G G C T A T A G C C G A T G T A C C T G A A G C T | JunB(bZIP)/DendriticCells-Junb-ChIP-Seq(GSE36099)/Homer | 1e-393 | -9.060e+02 | 0.0000 | 682.0 | 24.68% | 1332.2 | 3.04% | motif file (matrix) | svg |
| 6 | C T A G T C G A C G A T A C T G C G T A T A C G A G C T T G A C G C T A A C G T G A T C T A G C | Fosl2(bZIP)/3T3L1-Fosl2-ChIP-Seq(GSE56872)/Homer | 1e-389 | -8.972e+02 | 0.0000 | 577.0 | 20.88% | 864.0 | 1.97% | motif file (matrix) | svg |
| 7 | T C G A A C G T C A T G G C T A T A G C C G A T G T A C G C T A A C G T A T G C | AP-1(bZIP)/ThioMac-PU.1-ChIP-Seq(GSE21512)/Homer | 1e-377 | -8.703e+02 | 0.0000 | 765.0 | 27.69% | 1878.7 | 4.29% | motif file (matrix) | svg |
| 8 | C T A G T C G A A C G T A C T G C G T A T A G C C G A T G T A C C G T A A G C T G A T C G T A C | Jun-AP1(bZIP)/K562-cJun-ChIP-Seq(GSE31477)/Homer | 1e-334 | -7.713e+02 | 0.0000 | 468.0 | 16.94% | 621.8 | 1.42% | motif file (matrix) | svg |
| 9 | T C G A C T A G A G T C A G T C C G T A C G T A A C G T T A G C T C A G T A C G | NFY(CCAAT)/Promoter/Homer | 1e-262 | -6.053e+02 | 0.0000 | 819.0 | 29.64% | 3245.9 | 7.41% | motif file (matrix) | svg |
| 10 | C G A T T A C G T G A C G A C T C A T G C G T A T A C G A C G T G T A C C T G A | Bach2(bZIP)/OCILy7-Bach2-ChIP-Seq(GSE44420)/Homer | 1e-207 | -4.776e+02 | 0.0000 | 334.0 | 12.09% | 550.9 | 1.26% | motif file (matrix) | svg |
| 11 | T A C G C T A G A T G C G A T C G T A C A G T C C T A G A G T C A G T C A G T C G T A C A G T C | Sp1(Zf)/Promoter/Homer | 1e-156 | -3.596e+02 | 0.0000 | 684.0 | 24.76% | 3494.9 | 7.98% | motif file (matrix) | svg |
| 12 | C T G A T C A G C A G T C T A G A C T G C T A G G A T C A T C G A C T G C T G A T C A G G A T C | Sp5(Zf)/mES-Sp5.Flag-ChIP-Seq(GSE72989)/Homer | 1e-136 | -3.136e+02 | 0.0000 | 1145.0 | 41.44% | 8991.6 | 20.54% | motif file (matrix) | svg |
| 13 | G A C T T C A G C T A G A G T C A G T C G T A C A G T C C T G A A G T C A G T C A G T C G A C T A G T C A C T G A T G C | KLF3(Zf)/MEF-Klf3-ChIP-Seq(GSE44748)/Homer | 1e-123 | -2.843e+02 | 0.0000 | 712.0 | 25.77% | 4362.0 | 9.96% | motif file (matrix) | svg |
| 14 | C G T A C T A G A C T G A C T G G A C T C T A G C A G T C T A G C A T G G A T C | KLF5(Zf)/LoVo-KLF5-ChIP-Seq(GSE49402)/Homer | 1e-93 | -2.145e+02 | 0.0000 | 1174.0 | 42.49% | 10775.4 | 24.61% | motif file (matrix) | svg |
| 15 | T A C G T C G A C A G T A C T G G C T A A T G C C G A T G T A C C G T A A C T G T A G C C G T A | NF-E2(bZIP)/K562-NFE2-ChIP-Seq(GSE31477)/Homer | 1e-88 | -2.035e+02 | 0.0000 | 126.0 | 4.56% | 166.9 | 0.38% | motif file (matrix) | svg |
| 16 | G T A C C A G T A C T G A C T G A C T G G A T C A C T G A C G T A C T G A C T G A G T C G A T C | KLF6(Zf)/PDAC-KLF6-ChIP-Seq(GSE64557)/Homer | 1e-86 | -1.981e+02 | 0.0000 | 985.0 | 35.65% | 8562.2 | 19.56% | motif file (matrix) | svg |
| 17 | C T A G T C A G C A G T T C A G A C T G A C T G G A T C C T A G A C T G C T A G T C A G A T G C | KLF14(Zf)/HEK293-KLF14.GFP-ChIP-Seq(GSE58341)/Homer | 1e-73 | -1.684e+02 | 0.0000 | 1372.0 | 49.66% | 14408.3 | 32.91% | motif file (matrix) | svg |
| 18 | A T G C A G C T T C A G T G A C T C A G A T G C T G C A A C G T A T C G G A T C A C T G A G T C | NRF1(NRF)/MCF7-NRF1-ChIP-Seq(Unpublished)/Homer | 1e-72 | -1.677e+02 | 0.0000 | 331.0 | 11.98% | 1661.8 | 3.80% | motif file (matrix) | svg |
| 19 | T C A G C G T A A G T C A G C T C G T A A G T C C T G A C G T A A G T C G C A T A G T C A G T C A G T C C T G A A C T G T G C A T C G A C A T G A T C G G A T C | Ronin(THAP)/ES-Thap11-ChIP-Seq(GSE51522)/Homer | 1e-69 | -1.596e+02 | 0.0000 | 150.0 | 5.43% | 367.5 | 0.84% | motif file (matrix) | svg |
| 20 | G T C A G C A T A C T G G T A C G A C T A C T G G C T A A T C G C A G T G T A C C G T A A G C T | Nrf2(bZIP)/Lymphoblast-Nrf2-ChIP-Seq(GSE37589)/Homer | 1e-69 | -1.591e+02 | 0.0000 | 102.0 | 3.69% | 143.5 | 0.33% | motif file (matrix) | svg |
| 21 | T C A G T G A C G T A C T G C A G T A C C T A G G T A C A T G C A G T C G T C A A G T C G A C T | Klf9(Zf)/GBM-Klf9-ChIP-Seq(GSE62211)/Homer | 1e-68 | -1.589e+02 | 0.0000 | 482.0 | 17.44% | 3197.6 | 7.30% | motif file (matrix) | svg |
| 22 | C G T A C G T A C G T A G C A T G C A T T A C G G T A C G A C T A C T G C G T A A T C G A C G T G T A C C G T A A G C T | Bach1(bZIP)/K562-Bach1-ChIP-Seq(GSE31477)/Homer | 1e-68 | -1.581e+02 | 0.0000 | 95.0 | 3.44% | 119.7 | 0.27% | motif file (matrix) | svg |
| 23 | A T C G A G C T A C T G A G T C A C T G A G T C C G T A A C G T A C T G A G T C A C T G A G T C | NRF(NRF)/Promoter/Homer | 1e-64 | -1.487e+02 | 0.0000 | 328.0 | 11.87% | 1776.5 | 4.06% | motif file (matrix) | svg |
| 24 | T C A G T A G C G A C T C A T G C T G A A T C G G C A T G T A C C G T A A C T G T A G C T G C A | MafK(bZIP)/C2C12-MafK-ChIP-Seq(GSE36030)/Homer | 1e-64 | -1.474e+02 | 0.0000 | 212.0 | 7.67% | 818.0 | 1.87% | motif file (matrix) | svg |
| 25 | A G T C C T G A A G T C C G A T C A G T G A T C A T G C A C T G A T C G G A C T | Fli1(ETS)/CD8-FLI-ChIP-Seq(GSE20898)/Homer | 1e-63 | -1.452e+02 | 0.0000 | 735.0 | 26.60% | 6264.8 | 14.31% | motif file (matrix) | svg |
| 26 | T C G A C G T A A G T C A G C T C G T A A G T C T C G A G C T A G A C T C G A T A G T C A G T C A G T C C T G A T C A G T G C A T C G A C A G T A T C G A G T C | GFY-Staf(?,Zf)/Promoter/Homer | 1e-62 | -1.431e+02 | 0.0000 | 159.0 | 5.75% | 473.7 | 1.08% | motif file (matrix) | svg |
| 27 | G A T C C T G A A G T C C G A T C G A T G A T C A G T C A C T G A T C G A G C T | Elk4(ETS)/Hela-Elk4-ChIP-Seq(GSE31477)/Homer | 1e-59 | -1.376e+02 | 0.0000 | 539.0 | 19.51% | 4066.2 | 9.29% | motif file (matrix) | svg |
| 28 | G A T C T C G A A G T C C G A T C G A T A G T C A T G C A C T G A T C G G A C T | Elk1(ETS)/Hela-Elk1-ChIP-Seq(GSE31477)/Homer | 1e-58 | -1.354e+02 | 0.0000 | 537.0 | 19.44% | 4072.9 | 9.30% | motif file (matrix) | svg |
| 29 | T G C A A G C T C T G A A T C G G A C T C T A G G T A C G A T C G T C A A G T C G T A C G A C T C T A G A T C G G C A T C A T G C A T G G A T C G T A C C T G A | CTCF(Zf)/CD4+-CTCF-ChIP-Seq(Barski\_et\_al.)/Homer | 1e-57 | -1.328e+02 | 0.0000 | 169.0 | 6.12% | 579.7 | 1.32% | motif file (matrix) | svg |
| 30 | C T G A T G C A T A G C T G A C T A C G T C A G C T G A G C T A T C A G G A C T | ELF1(ETS)/Jurkat-ELF1-ChIP-Seq(SRA014231)/Homer | 1e-55 | -1.275e+02 | 0.0000 | 497.0 | 17.99% | 3726.3 | 8.51% | motif file (matrix) | svg |
| 31 | T G C A T C G A T A G C G T A C T C A G C T A G G T C A G C T A T C A G G A C T | ETS(ETS)/Promoter/Homer | 1e-53 | -1.229e+02 | 0.0000 | 360.0 | 13.03% | 2317.0 | 5.29% | motif file (matrix) | svg |
| 32 | T G C A T A G C G A C T T G C A T G A C T G C A C G T A A G C T A G C T A G T C A G T C G T A C | GFY(?)/Promoter/Homer | 1e-51 | -1.187e+02 | 0.0000 | 142.0 | 5.14% | 457.6 | 1.05% | motif file (matrix) | svg |
| 33 | T A G C G C T A T C G A C T G A A G T C A G T C C T G A A G T C C G T A C T A G | RUNX(Runt)/HPC7-Runx1-ChIP-Seq(GSE22178)/Homer | 1e-47 | -1.104e+02 | 0.0000 | 343.0 | 12.41% | 2274.6 | 5.20% | motif file (matrix) | svg |
| 34 | T C G A C T G A T A G C T G A C T C A G T C A G C G T A C G T A T C A G A G C T | ETV1(ETS)/GIST48-ETV1-ChIP-Seq(GSE22441)/Homer | 1e-44 | -1.021e+02 | 0.0000 | 742.0 | 26.85% | 7120.0 | 16.26% | motif file (matrix) | svg |
| 35 | G C T A C T G A T C G A A G T C A G T C C T G A A G T C G T C A C T G A T G C A | RUNX1(Runt)/Jurkat-RUNX1-ChIP-Seq(GSE29180)/Homer | 1e-43 | -1.008e+02 | 0.0000 | 412.0 | 14.91% | 3131.4 | 7.15% | motif file (matrix) | svg |
| 36 | T C G A T C G A T A G C G T A C T C A G T A C G C G T A C G T A T C A G A G C T | GABPA(ETS)/Jurkat-GABPa-ChIP-Seq(GSE17954)/Homer | 1e-43 | -1.002e+02 | 0.0000 | 561.0 | 20.30% | 4880.3 | 11.15% | motif file (matrix) | svg |
| 37 | A T G C A T C G T A C G A G C T A T C G C T G A A G T C C T A G A G C T A T G C C T G A A T G C | CRE(bZIP)/Promoter/Homer | 1e-43 | -1.002e+02 | 0.0000 | 239.0 | 8.65% | 1352.1 | 3.09% | motif file (matrix) | svg |
| 38 | T C G A T A G C T G C A A C T G A C T G C G T A C G T A C T A G G A C T T A C G | ETS1(ETS)/Jurkat-ETS1-ChIP-Seq(GSE17954)/Homer | 1e-40 | -9.243e+01 | 0.0000 | 577.0 | 20.88% | 5214.8 | 11.91% | motif file (matrix) | svg |
| 39 | C T A G G T A C A G T C T G C A A G T C C T G A A G T C A G T C A G T C G C T A | Klf4(Zf)/mES-Klf4-ChIP-Seq(GSE11431)/Homer | 1e-39 | -9.158e+01 | 0.0000 | 372.0 | 13.46% | 2811.0 | 6.42% | motif file (matrix) | svg |
| 40 | T G A C G C T A T C G A T G C A A G T C A G T C C G T A A G T C C G T A C T G A G C T A G T A C | RUNX2(Runt)/PCa-RUNX2-ChIP-Seq(GSE33889)/Homer | 1e-38 | -8.974e+01 | 0.0000 | 325.0 | 11.76% | 2324.0 | 5.31% | motif file (matrix) | svg |
| 41 | T C G A T A G C G T C A A C T G A C T G C G T A C G T A C T A G A G C T T C A G | ERG(ETS)/VCaP-ERG-ChIP-Seq(GSE14097)/Homer | 1e-38 | -8.915e+01 | 0.0000 | 727.0 | 26.31% | 7198.8 | 16.44% | motif file (matrix) | svg |
| 42 | T A G C A G T C T G A C A G T C C T A G A T C G A G T C C A T G T G A C A G T C G T A C A G T C A G T C G C A T C T A G A T C G G C A T A C T G A T C G G A T C | BORIS(Zf)/K562-CTCFL-ChIP-Seq(GSE32465)/Homer | 1e-38 | -8.913e+01 | 0.0000 | 234.0 | 8.47% | 1403.0 | 3.20% | motif file (matrix) | svg |
| 43 | A C T G G A T C G A C T A C T G A C G T C A T G A C T G A C G T A G C T C G A T | RUNX-AML(Runt)/CD4+-PolII-ChIP-Seq(Barski\_et\_al.)/Homer | 1e-38 | -8.913e+01 | 0.0000 | 308.0 | 11.15% | 2150.3 | 4.91% | motif file (matrix) | svg |
| 44 | G A C T C T A G G A T C C A G T A C T G C T G A A T G C G C A T A T G C C T G A | MafA(bZIP)/Islet-MafA-ChIP-Seq(GSE30298)/Homer | 1e-36 | -8.360e+01 | 0.0000 | 380.0 | 13.75% | 3013.6 | 6.88% | motif file (matrix) | svg |
| 45 | A G T C A T C G C T A G A G C T G A C T C T A G A G T C A G T C G C T A C A G T T C A G T C A G G A T C C T G A T C G A G A T C | RFX(HTH)/K562-RFX3-ChIP-Seq(SRA012198)/Homer | 1e-35 | -8.275e+01 | 0.0000 | 121.0 | 4.38% | 473.9 | 1.08% | motif file (matrix) | svg |
| 46 | C T A G A G C T G A C T C A T G A G T C A G T C G T C A C A G T C T A G T C A G G T A C C T G A T C G A G A T C T G A C | Rfx2(HTH)/LoVo-RFX2-ChIP-Seq(GSE49402)/Homer | 1e-33 | -7.647e+01 | 0.0000 | 120.0 | 4.34% | 499.1 | 1.14% | motif file (matrix) | svg |
| 47 | A T G C A G T C C T G A A G T C C G A T A C G T A G T C A G T C A C G T A T C G G A C T A C G T | Etv2(ETS)/ES-ER71-ChIP-Seq(GSE59402)/Homer(0.967) | 1e-32 | -7.375e+01 | 0.0000 | 461.0 | 16.68% | 4133.7 | 9.44% | motif file (matrix) | svg |
| 48 | C T A G A C T G C T A G T C A G T C A G T A C G C T A G A C T G | Maz(Zf)/HepG2-Maz-ChIP-Seq(GSE31477)/Homer | 1e-30 | -7.129e+01 | 0.0000 | 936.0 | 33.88% | 10511.7 | 24.01% | motif file (matrix) | svg |
| 49 | A T G C T C A G T C G A G C A T A C T G C G T A A G T C T C A G G A C T T G A C C G T A A G C T | Atf2(bZIP)/3T3L1-Atf2-ChIP-Seq(GSE56872)/Homer | 1e-27 | -6.359e+01 | 0.0000 | 205.0 | 7.42% | 1376.3 | 3.14% | motif file (matrix) | svg |
| 50 | T C A G A G C T A T G C C G T A A G C T T C A G C A G T A C T G C T G A A G T C | MITF(bHLH)/MastCells-MITF-ChIP-Seq(GSE48085)/Homer | 1e-26 | -6.176e+01 | 0.0000 | 412.0 | 14.91% | 3762.7 | 8.60% | motif file (matrix) | svg |
| 51 | A T G C A T G C A T C G T A C G A G C T A G T C G C T A A G T C T C A G G A C T A C T G T C G A | E-box(bHLH)/Promoter/Homer | 1e-23 | -5.517e+01 | 0.0000 | 107.0 | 3.87% | 527.9 | 1.21% | motif file (matrix) | svg |
| 52 | A C T G T C A G A G C T G A C T C A T G A G T C A G T C G C T A C G A T C T A G T C A G G T A C C T G A T C G A | Rfx1(HTH)/NPC-H3K4me1-ChIP-Seq(GSE16256)/Homer | 1e-23 | -5.462e+01 | 0.0000 | 143.0 | 5.18% | 856.4 | 1.96% | motif file (matrix) | svg |
| 53 | T C A G A G C T A T G C C G T A A G T C T C A G A C G T A T C G T C G A A G T C G A T C T G A C | TFE3(bHLH)/MEF-TFE3-ChIP-Seq(GSE75757)/Homer | 1e-23 | -5.344e+01 | 0.0000 | 89.0 | 3.22% | 391.0 | 0.89% | motif file (matrix) | svg |
| 54 | C G T A G A C T C G A T A T C G G T A C G C A T C A T G C G T A T A C G G C A T G T A C C G T A C A T G A T G C G C T A C T A G G C A T G C A T G C A T G A C T | MafB(bZIP)/BMM-Mafb-ChIP-Seq(GSE75722)/Homer | 1e-22 | -5.260e+01 | 0.0000 | 181.0 | 6.55% | 1256.5 | 2.87% | motif file (matrix) | svg |
| 55 | T C A G C A T G C A T G A C T G A C T G A G C T A C T G A C G T A C T G C A G T A T G C A G T C | KLF10(Zf)/HEK293-KLF10.GFP-ChIP-Seq(GSE58341)/Homer | 1e-22 | -5.245e+01 | 0.0000 | 368.0 | 13.32% | 3407.1 | 7.78% | motif file (matrix) | svg |
| 56 | A T G C A G T C G T A C A G C T T C G A C T A G G A T C C T G A G T C A A G T C G C T A T C A G | Rfx5(HTH)/GM12878-Rfx5-ChIP-Seq(GSE31477)/Homer | 1e-22 | -5.120e+01 | 0.0000 | 179.0 | 6.48% | 1252.1 | 2.86% | motif file (matrix) | svg |
| 57 | T C G A G C A T A C T G C T G A A G T C T C A G G A C T G T A C C G T A A G C T A G T C G A T C | c-Jun-CRE(bZIP)/K562-cJun-ChIP-Seq(GSE31477)/Homer | 1e-22 | -5.104e+01 | 0.0000 | 179.0 | 6.48% | 1254.3 | 2.87% | motif file (matrix) | svg |
| 58 | T C G A A C G T A C T G C T G A A G T C T C A G A G C T G T A C C G T A A G C T G A T C T C G A | JunD(bZIP)/K562-JunD-ChIP-Seq/Homer | 1e-21 | -4.842e+01 | 0.0000 | 85.0 | 3.08% | 390.6 | 0.89% | motif file (matrix) | svg |
| 59 | C G A T C T A G A C G T G T C A C G T A C G T A A G T C C G T A | Foxo3(Forkhead)/U2OS-Foxo3-ChIP-Seq(E-MTAB-2701)/Homer | 1e-21 | -4.840e+01 | 0.0000 | 176.0 | 6.37% | 1255.6 | 2.87% | motif file (matrix) | svg |
| 60 | T G C A C T G A A G T C G T C A A C T G A C T G C G T A C G T A C T G A A G C T | EWS:FLI1-fusion(ETS)/SK\_N\_MC-EWS:FLI1-ChIP-Seq(SRA014231)/Homer | 1e-20 | -4.800e+01 | 0.0000 | 307.0 | 11.11% | 2752.6 | 6.29% | motif file (matrix) | svg |
| 61 | C A T G C T A G A G C T G A C T C A T G A G T C G A T C G C T A C G A T C T A G T C A G G T A C C T G A T C G A | X-box(HTH)/NPC-H3K4me1-ChIP-Seq(GSE16256)/Homer | 1e-20 | -4.698e+01 | 0.0000 | 83.0 | 3.00% | 383.4 | 0.88% | motif file (matrix) | svg |
| 62 | T G C A G C A T A G C T G C A T A G T C A G T C A G T C C T G A A C T G T C G A T C G A C A G T A T C G A G T C G A T C | ZNF143\|STAF(Zf)/CUTLL-ZNF143-ChIP-Seq(GSE29600)/Homer | 1e-20 | -4.651e+01 | 0.0000 | 218.0 | 7.89% | 1739.9 | 3.97% | motif file (matrix) | svg |
| 63 | A G T C G A C T C A G T A C T G C T A G T G A C G C T A A T G C G C A T A T C G C G A T A C T G G A T C G T A C G T C A C T G A | NF1(CTF)/LNCAP-NF1-ChIP-Seq(Unpublished)/Homer | 1e-19 | -4.468e+01 | 0.0000 | 239.0 | 8.65% | 2008.8 | 4.59% | motif file (matrix) | svg |
| 64 | C T A G T A C G G A T C G T A C G C T A A G C T A G C T G T C A T C G A T A G C | Nanog(Homeobox)/mES-Nanog-ChIP-Seq(GSE11724)/Homer | 1e-19 | -4.398e+01 | 0.0000 | 1071.0 | 38.76% | 13417.8 | 30.65% | motif file (matrix) | svg |
| 65 | T A C G T C A G A G C T A T G C C G T A A G T C T C A G A C G T A C T G T C G A | USF1(bHLH)/GM12878-Usf1-ChIP-Seq(GSE32465)/Homer | 1e-17 | -4.037e+01 | 0.0000 | 259.0 | 9.37% | 2322.9 | 5.31% | motif file (matrix) | svg |
| 66 | T A G C C T A G T C G A G A C T A C T G C T G A A G T C T C A G G C A T T G A C C T G A A G C T | Atf7(bZIP)/3T3L1-Atf7-ChIP-Seq(GSE56872)/Homer | 1e-16 | -3.853e+01 | 0.0000 | 257.0 | 9.30% | 2334.5 | 5.33% | motif file (matrix) | svg |
| 67 | A G C T G C A T A C T G A C G T A G T C A C G T C T A G T A C G | Smad3(MAD)/NPC-Smad3-ChIP-Seq(GSE36673)/Homer | 1e-16 | -3.827e+01 | 0.0000 | 931.0 | 33.70% | 11581.5 | 26.46% | motif file (matrix) | svg |
| 68 | T A C G T C G A G A C T A C T G C T G A A G T C T C A G G A C T T G A C C T G A | Atf1(bZIP)/K562-ATF1-ChIP-Seq(GSE31477)/Homer | 1e-15 | -3.546e+01 | 0.0000 | 325.0 | 11.76% | 3248.9 | 7.42% | motif file (matrix) | svg |
| 69 | C G T A T A G C T A G C T G C A A C T G C T A G C G T A C G T A T C A G G A C T | EHF(ETS)/LoVo-EHF-ChIP-Seq(GSE49402)/Homer | 1e-15 | -3.517e+01 | 0.0000 | 481.0 | 17.41% | 5313.5 | 12.14% | motif file (matrix) | svg |
| 70 | A G C T A G T C A G T C A C G T C T A G A C G T A C G T A C G T C G T A A G T C G A T C C G T A | FOXP1(Forkhead)/H9-FOXP1-ChIP-Seq(GSE31006)/Homer | 1e-14 | -3.405e+01 | 0.0000 | 110.0 | 3.98% | 744.3 | 1.70% | motif file (matrix) | svg |
| 71 | C T A G G C A T G A T C C G T A A G T C T C A G G A C T C T A G | CLOCK(bHLH)/Liver-Clock-ChIP-Seq(GSE39860)/Homer | 1e-14 | -3.393e+01 | 0.0000 | 270.0 | 9.77% | 2591.4 | 5.92% | motif file (matrix) | svg |
| 72 | G C A T G C A T C T G A A C G T C T G A A C G T C G T A C G T A C G T A A G T C G T C A G T C A | Foxf1(Forkhead)/Lung-Foxf1-ChIP-Seq(GSE77951)/Homer | 1e-14 | -3.331e+01 | 0.0000 | 176.0 | 6.37% | 1474.2 | 3.37% | motif file (matrix) | svg |
| 73 | C A G T T C A G A G C T G A C T A C G T A G T C G A T C G A C T C T G A A C T G G A T C C G T A C T G A A G T C G T A C | Rfx6(HTH)/Min6b1-Rfx6.HA-ChIP-Seq(GSE62844)/Homer | 1e-14 | -3.303e+01 | 0.0000 | 447.0 | 16.18% | 4921.7 | 11.24% | motif file (matrix) | svg |
| 74 | C A T G G T A C C G T A A G T C C T A G A C G T A C T G G T A C A G T C A G C T | bHLHE40(bHLH)/HepG2-BHLHE40-ChIP-Seq(GSE31477)/Homer | 1e-14 | -3.266e+01 | 0.0000 | 199.0 | 7.20% | 1754.7 | 4.01% | motif file (matrix) | svg |
| 75 | T C A G A C G T A G T C T C G A A G T C T C A G G C A T C T A G C T A G A G C T | Usf2(bHLH)/C2C12-Usf2-ChIP-Seq(GSE36030)/Homer | 1e-14 | -3.262e+01 | 0.0000 | 198.0 | 7.17% | 1743.7 | 3.98% | motif file (matrix) | svg |
| 76 | A G T C G A C T C A G T G T A C A G T C A T C G T C A G A C T G G T C A C G T A | Stat3(Stat)/mES-Stat3-ChIP-Seq(GSE11431)/Homer | 1e-13 | -3.220e+01 | 0.0000 | 212.0 | 7.67% | 1918.7 | 4.38% | motif file (matrix) | svg |
| 77 | A G C T T G A C C G A T C G A T C T A G A C G T C A G T C A G T G C T A A G T C | FOXK1(Forkhead)/HEK293-FOXK1-ChIP-Seq(GSE51673)/Homer | 1e-13 | -3.113e+01 | 0.0000 | 223.0 | 8.07% | 2073.2 | 4.74% | motif file (matrix) | svg |
| 78 | A T G C T C G A A G T C A G C T A C G T G T A C A G T C G C T A C T A G C A T G G T C A C T G A T C A G A G T C | Stat3+il21(Stat)/CD4-Stat3-ChIP-Seq(GSE19198)/Homer | 1e-13 | -3.072e+01 | 0.0000 | 245.0 | 8.87% | 2354.0 | 5.38% | motif file (matrix) | svg |
| 79 | C T A G C T A G A G T C T C A G A C T G A C G T A C G T C T G A | MYB(HTH)/ERMYB-Myb-ChIPSeq(GSE22095)/Homer | 1e-13 | -3.050e+01 | 0.0000 | 573.0 | 20.74% | 6745.2 | 15.41% | motif file (matrix) | svg |
| 80 | A T G C G A C T A C G T C T A G A C G T A C G T A C G T C T G A G A T C G C T A A G C T C G T A | Foxa2(Forkhead)/Liver-Foxa2-ChIP-Seq(GSE25694)/Homer | 1e-13 | -3.013e+01 | 0.0000 | 176.0 | 6.37% | 1530.6 | 3.50% | motif file (matrix) | svg |
| 81 | G C T A T C G A C G T A C T A G A G C T G T C A G T C A C G T A A G T C C G T A | FOXA1(Forkhead)/LNCAP-FOXA1-ChIP-Seq(GSE27824)/Homer | 1e-12 | -2.874e+01 | 0.0000 | 228.0 | 8.25% | 2187.5 | 5.00% | motif file (matrix) | svg |
| 82 | G C T A A G T C T A C G T G C A A T C G T C A G G C T A T C G A T C A G A G C T | ELF5(ETS)/T47D-ELF5-ChIP-Seq(GSE30407)/Homer | 1e-12 | -2.846e+01 | 0.0000 | 292.0 | 10.57% | 3005.5 | 6.87% | motif file (matrix) | svg |
| 83 | C T G A A T G C C G T A A C G T A G T C A G T C A C G T A C T G A T C G G C A T | SPDEF(ETS)/VCaP-SPDEF-ChIP-Seq(SRA014231)/Homer | 1e-12 | -2.828e+01 | 0.0000 | 382.0 | 13.83% | 4197.4 | 9.59% | motif file (matrix) | svg |
| 84 | T C G A T A G C G T C A A C T G C T A G C G T A C G A T A C T G A C G T A C T G A C T G A C G T | ETS:RUNX(ETS,Runt)/Jurkat-RUNX1-ChIP-Seq(GSE17954)/Homer | 1e-11 | -2.728e+01 | 0.0000 | 81.0 | 2.93% | 528.4 | 1.21% | motif file (matrix) | svg |
| 85 | A C G T C T A G A G C T A C G T A C G T C T G A A G T C G A C T A G C T C G T A | FOXM1(Forkhead)/MCF7-FOXM1-ChIP-Seq(GSE72977)/Homer | 1e-11 | -2.649e+01 | 0.0000 | 201.0 | 7.27% | 1904.8 | 4.35% | motif file (matrix) | svg |
| 86 | T C A G A G C T A C G T A C G T G T A C G A T C C G T A C T A G C A T G G T C A C G T A T C G A | STAT4(Stat)/CD4-Stat4-ChIP-Seq(GSE22104)/Homer | 1e-11 | -2.638e+01 | 0.0000 | 253.0 | 9.16% | 2559.6 | 5.85% | motif file (matrix) | svg |
| 87 | C G T A T A C G T C G A A C T G A C T G C G T A C G T A T A C G A G C T T A C G | PU.1(ETS)/ThioMac-PU.1-ChIP-Seq(GSE21512)/Homer | 1e-11 | -2.628e+01 | 0.0000 | 195.0 | 7.06% | 1835.3 | 4.19% | motif file (matrix) | svg |
| 88 | G C T A T C G A C G T A C T A G A G C T G T C A G T C A C G T A A G T C C G T A | FOXA1(Forkhead)/MCF7-FOXA1-ChIP-Seq(GSE26831)/Homer | 1e-11 | -2.565e+01 | 0.0000 | 188.0 | 6.80% | 1763.4 | 4.03% | motif file (matrix) | svg |
| 89 | C T A G C A G T T G A C C G T A G A T C T C A G G A C T C A T G | BMAL1(bHLH)/Liver-Bmal1-ChIP-Seq(GSE39860)/Homer | 1e-10 | -2.486e+01 | 0.0000 | 569.0 | 20.59% | 6917.2 | 15.80% | motif file (matrix) | svg |
| 90 | G A C T C T A G C T A G A G T C T G C A A C T G A C G T A C G T C T A G T C A G | AMYB(HTH)/Testes-AMYB-ChIP-Seq(GSE44588)/Homer | 1e-10 | -2.460e+01 | 0.0000 | 467.0 | 16.90% | 5491.1 | 12.54% | motif file (matrix) | svg |
| 91 | T A G C T A G C G C A T C A T G A C T G G C T A C G T A A C G T A C T G G A T C | TEAD4(TEA)/Tropoblast-Tead4-ChIP-Seq(GSE37350)/Homer | 1e-10 | -2.430e+01 | 0.0000 | 263.0 | 9.52% | 2744.7 | 6.27% | motif file (matrix) | svg |
| 92 | C G T A G C T A C G A T C T A G A C G T G T C A C G T A C G T A A G T C C G T A T G C A T A C G | FoxL2(Forkhead)/Ovary-FoxL2-ChIP-Seq(GSE60858)/Homer | 1e-10 | -2.402e+01 | 0.0000 | 157.0 | 5.68% | 1422.3 | 3.25% | motif file (matrix) | svg |
| 93 | T A G C T A G C G A C T C T A G A G C T A G T C G T C A T G C A A C G T A T G C G C T A T G C A | Pbx3(Homeobox)/GM12878-PBX3-ChIP-Seq(GSE32465)/Homer | 1e-10 | -2.388e+01 | 0.0000 | 104.0 | 3.76% | 813.6 | 1.86% | motif file (matrix) | svg |
| 94 | T C G A T A G C T G A C C T G A A G T C A C T G G A C T C A T G | c-Myc(bHLH)/LNCAP-cMyc-ChIP-Seq(Unpublished)/Homer | 1e-10 | -2.371e+01 | 0.0000 | 291.0 | 10.53% | 3128.6 | 7.15% | motif file (matrix) | svg |
| 95 | A G T C A C G T A C T G A G C T A C G T A C G T G T C A A G T C | Foxo1(Forkhead)/RAW-Foxo1-ChIP-Seq(Fan\_et\_al.)/Homer | 1e-10 | -2.366e+01 | 0.0000 | 482.0 | 17.44% | 5738.0 | 13.11% | motif file (matrix) | svg |
| 96 | T C G A T G A C A G T C C G T A A G T C C T A G A C G T A C T G A C T G A G C T A G T C G C A T | Max(bHLH)/K562-Max-ChIP-Seq(GSE31477)/Homer | 1e-10 | -2.331e+01 | 0.0000 | 273.0 | 9.88% | 2903.3 | 6.63% | motif file (matrix) | svg |
| 97 | A G C T C A T G G C A T G A T C T G C A C T A G G A T C A C G T | Tgif2(Homeobox)/mES-Tgif2-ChIP-Seq(GSE55404)/Homer | 1e-9 | -2.223e+01 | 0.0000 | 837.0 | 30.29% | 10956.2 | 25.03% | motif file (matrix) | svg |
| 98 | T C G A A G C T A C G T A C G T A G T C A G T C A C G T A T C G G A C T A T C G | EWS:ERG-fusion(ETS)/CADO\_ES1-EWS:ERG-ChIP-Seq(SRA014231)/Homer | 1e-9 | -2.211e+01 | 0.0000 | 209.0 | 7.56% | 2110.0 | 4.82% | motif file (matrix) | svg |
| 99 | T A C G A T G C G A C T A C T G A G C T A G T C G T C A T G C A A C G T A G T C G C T A T G C A | Pknox1(Homeobox)/ES-Prep1-ChIP-Seq(GSE63282)/Homer | 1e-9 | -2.164e+01 | 0.0000 | 105.0 | 3.80% | 859.7 | 1.96% | motif file (matrix) | svg |
| 100 | C G T A T G A C T A G C T G C A A C T G A C T G C G T A C G T A T C A G G A C T | ELF3(ETS)/PDAC-ELF3-ChIP-Seq(GSE64557)/Homer | 1e-9 | -2.152e+01 | 0.0000 | 251.0 | 9.08% | 2668.4 | 6.10% | motif file (matrix) | svg |
| 101 | G A T C G T A C C G A T A C T G A C T G C G T A C G T A A C G T A C T G G A T C | TEAD(TEA)/Fibroblast-PU.1-ChIP-Seq(Unpublished)/Homer | 1e-9 | -2.148e+01 | 0.0000 | 175.0 | 6.33% | 1697.9 | 3.88% | motif file (matrix) | svg |
| 102 | C T A G C A T G T G C A A C G T A G T C C G T A C A T G T C A G A C G T A C G T G C T A A G T C | Six1(Homeobox)/Myoblast-Six1-ChIP-Chip(GSE20150)/Homer | 1e-9 | -2.098e+01 | 0.0000 | 79.0 | 2.86% | 583.4 | 1.33% | motif file (matrix) | svg |
| 103 | T A C G T A C G G T A C A T C G T A C G T A C G G T C A C T G A C G T A G A C T | E2F4(E2F)/K562-E2F4-ChIP-Seq(GSE31477)/Homer | 1e-8 | -2.028e+01 | 0.0000 | 391.0 | 14.15% | 4605.4 | 10.52% | motif file (matrix) | svg |
| 104 | A G T C C G A T A C T G A T C G T G A C G C T A C A T G A T C G T G A C C G A T A C T G T A G C G T A C G T C A | Tlx?(NR)/NPC-H3K4me1-ChIP-Seq(GSE16256)/Homer | 1e-8 | -1.976e+01 | 0.0000 | 182.0 | 6.59% | 1827.9 | 4.18% | motif file (matrix) | svg |
| 105 | A G T C T A G C G A C T A C G T C T A G A C G T A C G T A C G T C T G A A G T C G C T A G A C T C G T A C T A G A C T G | Foxa3(Forkhead)/Liver-Foxa3-ChIP-Seq(GSE77670)/Homer | 1e-8 | -1.922e+01 | 0.0000 | 72.0 | 2.61% | 532.9 | 1.22% | motif file (matrix) | svg |
| 106 | C T A G T C G A C G A T C T A G G C A T C A G T C T A G G A T C C G T A G T C A | CEBP:AP1(bZIP)/ThioMac-CEBPb-ChIP-Seq(GSE21512)/Homer | 1e-8 | -1.871e+01 | 0.0000 | 177.0 | 6.41% | 1791.2 | 4.09% | motif file (matrix) | svg |
| 107 | A C T G A C G T C A T G A T C G A T C G T G A C A C T G A T C G A T C G T G C A C T G A C G T A | E2F3(E2F)/MEF-E2F3-ChIP-Seq(GSE71376)/Homer | 1e-8 | -1.852e+01 | 0.0000 | 438.0 | 15.85% | 5339.2 | 12.20% | motif file (matrix) | svg |
| 108 | C G A T A C G T A C G T A C G T C G T A A G C T C A G T C T A G A T C G A C T G | HOXB13(Homeobox)/ProstateTumor-HOXB13-ChIP-Seq(GSE56288)/Homer | 1e-8 | -1.842e+01 | 0.0000 | 175.0 | 6.33% | 1773.4 | 4.05% | motif file (matrix) | svg |
| 109 | C G A T T G C A G T A C C G T A A G T C C T A G G A C T C A T G | NPAS(bHLH)/Liver-NPAS-ChIP-Seq(GSE39860)/Homer | 1e-7 | -1.816e+01 | 0.0000 | 500.0 | 18.10% | 6239.0 | 14.25% | motif file (matrix) | svg |
| 110 | A G T C G C A T C G T A C G T A G T A C A C G T A C T G G A T C G A T C T C G A | BMYB(HTH)/Hela-BMYB-ChIP-Seq(GSE27030)/Homer | 1e-7 | -1.812e+01 | 0.0000 | 413.0 | 14.95% | 5004.4 | 11.43% | motif file (matrix) | svg |
| 111 | T C G A G A C T A G C T T G A C A G C T G T A C T C A G G A T C A T C G T G C A A C T G C T G A | GFX(?)/Promoter/Homer | 1e-7 | -1.809e+01 | 0.0000 | 34.0 | 1.23% | 172.3 | 0.39% | motif file (matrix) | svg |
| 112 | T G C A C T G A A T G C G T C A A C T G A C T G C G T A C G T A C T A G A G C T | Ets1-distal(ETS)/CD4+-PolII-ChIP-Seq(Barski\_et\_al.)/Homer | 1e-7 | -1.790e+01 | 0.0000 | 118.0 | 4.27% | 1079.3 | 2.47% | motif file (matrix) | svg |
| 113 | A G C T A C G T A C T G A T G C A G T C C G T A C T G A T A C G | NF1-halfsite(CTF)/LNCaP-NF1-ChIP-Seq(Unpublished)/Homer | 1e-7 | -1.726e+01 | 0.0000 | 610.0 | 22.08% | 7881.2 | 18.00% | motif file (matrix) | svg |
| 114 | A T G C C T G A A T C G T A C G A G T C C G A T T C A G C G A T C T A G A G C T G T C A G T C A C G T A A G T C C G T A T A C G C T G A | Fox:Ebox(Forkhead,bHLH)/Panc1-Foxa2-ChIP-Seq(GSE47459)/Homer | 1e-7 | -1.725e+01 | 0.0000 | 206.0 | 7.46% | 2207.4 | 5.04% | motif file (matrix) | svg |
| 115 | G A C T G C A T C T A G C G A T G A T C T C G A C A T G G A T C | Tgif1(Homeobox)/mES-Tgif1-ChIP-Seq(GSE55404)/Homer | 1e-7 | -1.721e+01 | 0.0000 | 727.0 | 26.31% | 9610.5 | 21.95% | motif file (matrix) | svg |
| 116 | T A C G T C G A T A G C A G T C C G T A A G T C C T A G G C A T A C T G A T C G | n-Myc(bHLH)/mES-nMyc-ChIP-Seq(GSE11431)/Homer | 1e-7 | -1.661e+01 | 0.0000 | 304.0 | 11.00% | 3550.0 | 8.11% | motif file (matrix) | svg |
| 117 | T A G C C G T A C T G A T A C G C G T A A C G T A C T G A C T G A G T C T A C G C T A G G T A C | YY1(Zf)/Promoter/Homer | 1e-7 | -1.649e+01 | 0.0000 | 71.0 | 2.57% | 560.7 | 1.28% | motif file (matrix) | svg |
| 118 | G A C T G T A C T G C A A C G T G A T C G C T A T C G A A C G T A G T C C G T A | Pdx1(Homeobox)/Islet-Pdx1-ChIP-Seq(SRA008281)/Homer | 1e-6 | -1.584e+01 | 0.0000 | 169.0 | 6.12% | 1767.4 | 4.04% | motif file (matrix) | svg |
| 119 | G A C T C A G T A G C T C G A T A G T C G A T C A G T C C G T A A T G C T C A G | Rbpj1(?)/Panc1-Rbpj1-ChIP-Seq(GSE47459)/Homer | 1e-6 | -1.564e+01 | 0.0000 | 441.0 | 15.96% | 5519.1 | 12.61% | motif file (matrix) | svg |
| 120 | T A C G T C A G T G C A A G C T T G A C A G C T A G T C A C T G G A T C A C T G T C G A A C T G C T G A C T G A A T G C | ZBTB33(Zf)/GM12878-ZBTB33-ChIP-Seq(GSE32465)/Homer | 1e-6 | -1.554e+01 | 0.0000 | 59.0 | 2.14% | 443.5 | 1.01% | motif file (matrix) | svg |
| 121 | A T G C C T G A G A C T A C G T A C G T G T A C G A T C C G A T C T A G C A T G C G T A C G T A C T G A G A C T | STAT1(Stat)/HelaS3-STAT1-ChIP-Seq(GSE12782)/Homer | 1e-6 | -1.539e+01 | 0.0000 | 81.0 | 2.93% | 691.8 | 1.58% | motif file (matrix) | svg |
| 122 | T A C G T A G C G C T A C G A T C T A G A C G T C A G T C A G T G C T A A G T C G T C A G C A T | FOXK2(Forkhead)/U2OS-FOXK2-ChIP-Seq(E-MTAB-2204)/Homer | 1e-6 | -1.415e+01 | 0.0000 | 143.0 | 5.18% | 1481.0 | 3.38% | motif file (matrix) | svg |
| 123 | T A C G C T G A T C G A C G A T C T A G C T A G T C G A C T G A T C G A T C G A C G T A T C G A G C A T C A T G C G T A T A C G G C A T T G A C C G T A A G C T | NFAT:AP1(RHD,bZIP)/Jurkat-NFATC1-ChIP-Seq(Jolma\_et\_al.)/Homer | 1e-5 | -1.367e+01 | 0.0000 | 50.0 | 1.81% | 372.9 | 0.85% | motif file (matrix) | svg |
| 124 | C T A G C G T A G T C A C G T A A G T C G A T C A G C T C T A G C G T A A C G T G T C A G A T C | Six2(Homeobox)/NephronProgenitor-Six2-ChIP-Seq(GSE39837)/Homer | 1e-5 | -1.357e+01 | 0.0000 | 214.0 | 7.75% | 2437.8 | 5.57% | motif file (matrix) | svg |
| 125 | C T G A A C G T A C G T A C G T A G T C G A C T C G A T C T G A A C T G C G T A C G T A T C G A | STAT5(Stat)/mCD4+-Stat5-ChIP-Seq(GSE12346)/Homer | 1e-5 | -1.344e+01 | 0.0000 | 87.0 | 3.15% | 799.8 | 1.83% | motif file (matrix) | svg |
| 126 | A T G C G A T C C G A T C T A G A C T G G C T A C G T A A G C T A C T G A G C T | TEAD2(TEA)/Py2T-Tead2-ChIP-Seq(GSE55709)/Homer | 1e-5 | -1.314e+01 | 0.0000 | 151.0 | 5.47% | 1613.2 | 3.69% | motif file (matrix) | svg |
| 127 | G T C A T G C A G C T A A G T C C G T A A C T G T G A C G C A T T C A G C A G T | Ap4(bHLH)/AML-Tfap4-ChIP-Seq(GSE45738)/Homer | 1e-5 | -1.313e+01 | 0.0000 | 458.0 | 16.58% | 5899.9 | 13.48% | motif file (matrix) | svg |
| 128 | T C A G T C A G G C T A C G T A T A C G G A C T T C A G T C G A C T G A C G T A T A C G G A C T | IRF8(IRF)/BMDM-IRF8-ChIP-Seq(GSE77884)/Homer | 1e-5 | -1.311e+01 | 0.0000 | 101.0 | 3.66% | 977.4 | 2.23% | motif file (matrix) | svg |
| 129 | C G T A C T G A C G T A C T A G T C G A C T A G A C T G C G T A C G T A T A C G A G C T A T C G | SpiB(ETS)/OCILY3-SPIB-ChIP-Seq(GSE56857)/Homer | 1e-5 | -1.293e+01 | 0.0000 | 89.0 | 3.22% | 834.9 | 1.91% | motif file (matrix) | svg |
| 130 | C T A G T A C G A G T C G T A C T C G A A C G T G T C A G C T A G C T A C G A T A G T C G C T A | HOXA9(Homeobox)/HSC-Hoxa9-ChIP-Seq(GSE33509)/Homer | 1e-5 | -1.249e+01 | 0.0000 | 118.0 | 4.27% | 1206.3 | 2.76% | motif file (matrix) | svg |
| 131 | T G C A A G C T C A T G C G T A A G C T A C T G G A T C G T C A C G T A A G C T | Atf4(bZIP)/MEF-Atf4-ChIP-Seq(GSE35681)/Homer | 1e-5 | -1.223e+01 | 0.0000 | 77.0 | 2.79% | 704.9 | 1.61% | motif file (matrix) | svg |
| 132 | T C A G G A C T C A G T C T G A A G C T C T A G G A C T T G C A C T G A A G T C | HLF(bZIP)/HSC-HLF.Flag-ChIP-Seq(GSE69817)/Homer | 1e-5 | -1.178e+01 | 0.0000 | 137.0 | 4.96% | 1472.9 | 3.36% | motif file (matrix) | svg |
| 133 | T G A C C G A T A C T G A C T G A C T G G A C T A C T G A C G T A C T G A C T G G A T C G A T C | EKLF(Zf)/Erythrocyte-Klf1-ChIP-Seq(GSE20478)/Homer | 1e-4 | -1.151e+01 | 0.0000 | 105.0 | 3.80% | 1067.9 | 2.44% | motif file (matrix) | svg |
| 134 | T A C G T A C G G T A C A T C G A C T G T A C G T C G A C T G A T C G A A T C G | E2F6(E2F)/Hela-E2F6-ChIP-Seq(GSE31477)/Homer | 1e-4 | -1.088e+01 | 0.0001 | 362.0 | 13.10% | 4639.1 | 10.60% | motif file (matrix) | svg |
| 135 | C T G A A C T G A C T G A G T C A G T C A G C T C T A G T A C G | ZFX(Zf)/mES-Zfx-ChIP-Seq(GSE11431)/Homer | 1e-4 | -1.079e+01 | 0.0001 | 596.0 | 21.57% | 8084.0 | 18.47% | motif file (matrix) | svg |
| 136 | A G T C C G A T C T G A C G T A A C G T C A G T T C A G T G A C | Isl1(Homeobox)/Neuron-Isl1-ChIP-Seq(GSE31456)/Homer | 1e-4 | -1.072e+01 | 0.0001 | 347.0 | 12.56% | 4432.4 | 10.13% | motif file (matrix) | svg |
| 137 | T G C A A G C T A C G T C T A G G A T C C T A G G A T C G T C A C T G A A G T C | CEBP(bZIP)/ThioMac-CEBPb-ChIP-Seq(GSE21512)/Homer | 1e-4 | -9.511e+00 | 0.0002 | 165.0 | 5.97% | 1927.6 | 4.40% | motif file (matrix) | svg |
| 138 | C G T A C G T A A G C T G A C T T G C A G T C A A C G T A G C T C T G A T C A G | Lhx3(Homeobox)/Neuron-Lhx3-ChIP-Seq(GSE31456)/Homer | 1e-4 | -9.451e+00 | 0.0002 | 257.0 | 9.30% | 3215.0 | 7.34% | motif file (matrix) | svg |
| 139 | G A C T C G A T T C A G G A T C G A C T A G C T A G C T A G T C G A T C C G T A C T A G C T A G T C G A T C G A C T G A | Bcl6(Zf)/Liver-Bcl6-ChIP-Seq(GSE31578)/Homer | 1e-3 | -9.045e+00 | 0.0003 | 320.0 | 11.58% | 4140.3 | 9.46% | motif file (matrix) | svg |
| 140 | A C T G C A T G G C T A T C G A G C T A A G C T A G C T G T A C A G T C T G A C | NFkB-p65-Rel(RHD)/ThioMac-LPS-Expression(GSE23622)/Homer | 1e-3 | -8.946e+00 | 0.0003 | 26.0 | 0.94% | 180.8 | 0.41% | motif file (matrix) | svg |
| 141 | T A G C C G A T A C G T A G C T A G C T A G T C A T G C A G T C A C T G A T G C A T G C G C T A | E2F7(E2F)/Hela-E2F7-ChIP-Seq(GSE32673)/Homer | 1e-3 | -8.868e+00 | 0.0004 | 105.0 | 3.80% | 1144.9 | 2.62% | motif file (matrix) | svg |
| 142 | A G C T A G C T T A G C A T C G A G T C A C T G A T G C A T C G T C G A C T G A T C G A C T G A | E2F(E2F)/Hela-CellCycle-Expression/Homer | 1e-3 | -8.406e+00 | 0.0006 | 49.0 | 1.77% | 447.8 | 1.02% | motif file (matrix) | svg |
| 143 | T A G C C G A T T A C G A C T G A G T C A C T G A T C G A T C G C G T A C T G A | E2F1(E2F)/Hela-E2F1-ChIP-Seq(GSE22478)/Homer | 1e-3 | -8.171e+00 | 0.0007 | 197.0 | 7.13% | 2431.5 | 5.55% | motif file (matrix) | svg |
| 144 | C T A G C T A G C G T A C G T A T A C G C G A T C T A G C T G A C T G A C G T A T A C G G A C T | PU.1:IRF8(ETS:IRF)/pDC-Irf8-ChIP-Seq(GSE66899)/Homer | 1e-3 | -8.021e+00 | 0.0008 | 55.0 | 1.99% | 528.2 | 1.21% | motif file (matrix) | svg |
| 145 | C T G A C A G T C T G A A G T C C T A G G A C T A T C G G T A C | HIF-1b(HLH)/T47D-HIF1b-ChIP-Seq(GSE59937)/Homer | 1e-3 | -7.955e+00 | 0.0009 | 453.0 | 16.40% | 6168.1 | 14.09% | motif file (matrix) | svg |
| 146 | A C G T T C G A T C G A A G T C G T C A T A C G A T G C A C G T A C T G A G C T | Myf5(bHLH)/GM-Myf5-ChIP-Seq(GSE24852)/Homer | 1e-3 | -7.576e+00 | 0.0013 | 255.0 | 9.23% | 3290.4 | 7.52% | motif file (matrix) | svg |
| 147 | C G A T C T A G T C G A A G C T C G A T C T G A C G T A A G C T A C T G C T A G A T G C G A T C | Hoxb4(Homeobox)/ES-Hoxb4-ChIP-Seq(GSE34014)/Homer | 1e-3 | -7.555e+00 | 0.0013 | 39.0 | 1.41% | 345.1 | 0.79% | motif file (matrix) | svg |
| 148 | C G A T T G C A T G C A G A T C C G T A A C T G T G A C G A C T C A T G A C T G | Tcf21(bHLH)/ArterySmoothMuscle-Tcf21-ChIP-Seq(GSE61369)/Homer | 1e-3 | -7.482e+00 | 0.0014 | 360.0 | 13.03% | 4827.2 | 11.03% | motif file (matrix) | svg |
| 149 | T A C G A T C G G A T C G T C A C T G A G C A T C G A T G C T A T C G A G C T A | Unknown(Homeobox)/Limb-p300-ChIP-Seq/Homer | 1e-3 | -7.463e+00 | 0.0014 | 94.0 | 3.40% | 1045.7 | 2.39% | motif file (matrix) | svg |
| 150 | G C A T A T C G C A T G G T A C G C T A A G T C T C A G T G A C G T C A T G C A | Arnt:Ahr(bHLH)/MCF7-Arnt-ChIP-Seq(Lo\_et\_al.)/Homer | 1e-3 | -7.411e+00 | 0.0015 | 288.0 | 10.42% | 3778.3 | 8.63% | motif file (matrix) | svg |
| 151 | A G T C G T A C A G C T C T A G A G T C C G A T A C T G C G T A A C T G G T C A | Zic(Zf)/Cerebellum-ZIC1.2-ChIP-Seq(GSE60731)/Homer | 1e-3 | -7.410e+00 | 0.0015 | 331.0 | 11.98% | 4406.1 | 10.07% | motif file (matrix) | svg |
| 152 | C T A G A T G C A T G C C G A T A C T G G A C T A T G C G C T A T G A C A G C T T A G C G C T A | PBX1(Homeobox)/MCF7-PBX1-ChIP-Seq(GSE28007)/Homer | 1e-3 | -7.294e+00 | 0.0016 | 35.0 | 1.27% | 303.6 | 0.69% | motif file (matrix) | svg |
| 153 | T A C G T C A G G A T C G T A C T C G A G A C T G C T A G C T A G C T A C G T A G A T C G T C A | CDX4(Homeobox)/ZebrafishEmbryos-Cdx4.Myc-ChIP-Seq(GSE48254)/Homer | 1e-3 | -7.121e+00 | 0.0019 | 119.0 | 4.31% | 1395.4 | 3.19% | motif file (matrix) | svg |
| 154 | A T G C A G T C G C A T A G C T A C G T T C A G C G A T A G C T G A T C A T C G | Sox10(HMG)/SciaticNerve-Sox3-ChIP-Seq(GSE35132)/Homer | 1e-3 | -7.033e+00 | 0.0021 | 330.0 | 11.94% | 4418.2 | 10.09% | motif file (matrix) | svg |
| 155 | C T A G T C G A C T G A C G T A T A C G G A C T T C A G T C G A G T C A T G C A T A C G A G C T | IRF2(IRF)/Erythroblas-IRF2-ChIP-Seq(GSE36985)/Homer | 1e-3 | -6.996e+00 | 0.0022 | 31.0 | 1.12% | 262.2 | 0.60% | motif file (matrix) | svg |
| 156 | C A G T T C A G G A T C A C T G A C G T C T A G A C T G A C T G G A C T C T A G | Egr1(Zf)/K562-Egr1-ChIP-Seq(GSE32465)/Homer | 1e-2 | -6.899e+00 | 0.0024 | 376.0 | 13.61% | 5107.9 | 11.67% | motif file (matrix) | svg |
| 157 | A C G T T A C G G A T C A C T G A C G T C T A G A C T G A C T G G A T C C T A G C A T G C T A G | Egr2(Zf)/Thymocytes-Egr2-ChIP-Seq(GSE34254)/Homer | 1e-2 | -6.805e+00 | 0.0026 | 128.0 | 4.63% | 1532.7 | 3.50% | motif file (matrix) | svg |
| 158 | G C T A A C G T C T A G G T C A C G T A A C G T C G T A C G A T C A G T A G T C C G T A A C G T C T A G C T G A A T C G | OCT:OCT(POU,Homeobox)/NPC-Brn1-ChIP-Seq(GSE35496)/Homer | 1e-2 | -6.701e+00 | 0.0028 | 7.0 | 0.25% | 25.8 | 0.06% | motif file (matrix) | svg |
| 159 | C G T A T A G C A G T C C T A G C A G T C T A G C T G A G T A C G C A T T C G A C G T A G C A T A G C T C T A G T C G A | PAX3:FKHR-fusion(Paired,Homeobox)/Rh4-PAX3:FKHR-ChIP-Seq(GSE19063)/Homer | 1e-2 | -6.695e+00 | 0.0028 | 40.0 | 1.45% | 374.3 | 0.86% | motif file (matrix) | svg |
| 160 | T G A C G C T A T G A C C G T A T C A G G A T C C G T A C A T G C A T G C T A G C T A G C T A G | Unknown-ESC-element(?)/mES-Nanog-ChIP-Seq(GSE11724)/Homer | 1e-2 | -6.645e+00 | 0.0030 | 230.0 | 8.32% | 2986.3 | 6.82% | motif file (matrix) | svg |
| 161 | T G C A C T G A A T G C G T C A A C G T A T G C A C G T A C T G A C T G T G C A | ZBTB18(Zf)/HEK293-ZBTB18.GFP-ChIP-Seq(GSE58341)/Homer | 1e-2 | -6.517e+00 | 0.0033 | 161.0 | 5.83% | 2008.0 | 4.59% | motif file (matrix) | svg |
| 162 | T C A G T A C G T A G C A C G T A C T G C G A T A G T C C G T A T A C G A G T C | Meis1(Homeobox)/MastCells-Meis1-ChIP-Seq(GSE48085)/Homer | 1e-2 | -6.363e+00 | 0.0039 | 474.0 | 17.16% | 6617.0 | 15.12% | motif file (matrix) | svg |
| 163 | C G A T G A C T C G A T T C A G G A C T A C G T C A G T C T G A G A C T G A C T A G C T C G A T A C T G A T C G G T A C G C T A | NF1:FOXA1(CTF,Forkhead)/LNCAP-FOXA1-ChIP-Seq(GSE27824)/Homer | 1e-2 | -5.852e+00 | 0.0064 | 12.0 | 0.43% | 73.7 | 0.17% | motif file (matrix) | svg |
| 164 | T G A C C T A G T C A G G T C A C G T A T C A G C G A T T C A G T C G A T G C A C T G A T A G C | PU.1-IRF(ETS:IRF)/Bcell-PU.1-ChIP-Seq(GSE21512)/Homer | 1e-2 | -5.843e+00 | 0.0064 | 341.0 | 12.34% | 4672.7 | 10.67% | motif file (matrix) | svg |
| 165 | C T A G T A C G G A T C G T A C C T G A A G C T T G C A G C T A C G T A G C A T G A T C G C T A | Hoxc9(Homeobox)/Ainv15-Hoxc9-ChIP-Seq(GSE21812)/Homer | 1e-2 | -5.802e+00 | 0.0067 | 77.0 | 2.79% | 876.5 | 2.00% | motif file (matrix) | svg |
| 166 | G A C T C G A T C T G A G T C A G A C T C G A T T C G A C G T A G C T A G C T A T G A C G T A C C G T A A C T G T G C A C G A T A C T G A C G T | Pitx1:Ebox(Homeobox,bHLH)/Hindlimb-Pitx1-ChIP-Seq(GSE41591)/Homer | 1e-2 | -5.457e+00 | 0.0094 | 26.0 | 0.94% | 231.7 | 0.53% | motif file (matrix) | svg |
| 167 | C G T A A C G T A C T G G T A C C G T A A C G T C G T A C G T A A C G T A C T G A G T C C G T A A C G T C T G A G C A T | OCT:OCT-short(POU,Homeobox)/NPC-OCT6-ChIP-Seq(GSE43916)/Homer | 1e-2 | -5.431e+00 | 0.0095 | 84.0 | 3.04% | 985.3 | 2.25% | motif file (matrix) | svg |
| 168 | A C T G G A C T A G T C C T G A G A T C T C A G A T G C G A C T A G T C A T G C T A G C A G C T A T C G T G C A | PAX5(Paired,Homeobox),condensed/GM12878-PAX5-ChIP-Seq(GSE32465)/Homer | 1e-2 | -5.263e+00 | 0.0112 | 52.0 | 1.88% | 562.1 | 1.28% | motif file (matrix) | svg |
| 169 | T C G A G C A T A C G T C T A G G T A C T C G A G C A T T G A C T C G A A C G T | Chop(bZIP)/MEF-Chop-ChIP-Seq(GSE35681)/Homer | 1e-2 | -5.192e+00 | 0.0120 | 50.0 | 1.81% | 538.6 | 1.23% | motif file (matrix) | svg |
| 170 | A G T C C T G A A T C G A G C T A G C T G A C T A G T C G C T A A C G T C G A T G C A T C G A T A T C G C G T A T A G C G C A T A T G C C G T A | bZIP:IRF(bZIP,IRF)/Th17-BatF-ChIP-Seq(GSE39756)/Homer | 1e-2 | -5.085e+00 | 0.0133 | 70.0 | 2.53% | 808.7 | 1.85% | motif file (matrix) | svg |
| 171 | T C A G A C G T T C G A T A G C A G T C C G T A A C T G G T A C A C G T A C T G A T C G A G T C | Atoh1(bHLH)/Cerebellum-Atoh1-ChIP-Seq(GSE22111)/Homer | 1e-2 | -4.824e+00 | 0.0171 | 358.0 | 12.96% | 5016.6 | 11.46% | motif file (matrix) | svg |
| 172 | G A C T C T A G C T A G C T A G A C T G T C G A C T G A C T A G C T A G C T A G G T A C G T C A | ZNF467(Zf)/HEK293-ZNF467.GFP-ChIP-Seq(GSE58341)/Homer | 1e-2 | -4.651e+00 | 0.0202 | 365.0 | 13.21% | 5138.4 | 11.74% | motif file (matrix) | svg |
