## Supplementary File 8.1.1 for "Promoter G-quadruplexes and transcription factors cooperate to shape the cell type-specific transcriptome": SupplementaryFile8.1.1_GS404_homerResults_denovoMotifs.html

GS404\_IP\_BG4\_93T\_m1\_F2L0\_peaks\_MotifOutput// - Homer de novo Motif Results


### Homer *de novo* Motif Results (GS404\_IP\_BG4\_93T\_m1\_F2L0\_peaks\_MotifOutput//)

Known Motif Enrichment Results  
Gene Ontology Enrichment Results  
If Homer is having trouble matching a motif to a known motif, try copy/pasting the matrix file into
STAMP  
More information on motif finding results: HOMER
| Description of Results
| Tips
  
Total target sequences = 3867  
Total background sequences = 43555  
\* - possible false positive  

|  |  |  |  |  |  |  |  |  |
| --- | --- | --- | --- | --- | --- | --- | --- | --- |
| Rank | Motif | P-value | log P-pvalue | % of Targets | % of Background | STD(Bg STD) | Best Match/Details | Motif File |
| 1 | A C T G T A G C C A G T A C T G G T C A T A C G C G A T T A G C C G T A A G C T A G T C T C A G | 1e-490 | -1.130e+03 | 24.08% | 3.36% | 48.6bp (61.5bp) | Fosl2(bZIP)/3T3L1-Fosl2-ChIP-Seq(GSE56872)/Homer(0.981) More Information \| Similar Motifs Found | motif file (matrix) |
| 2 | T C G A C T A G A G T C A T G C C G T A C T G A C G A T T A G C T C G A T C A G T G C A C T A G | 1e-247 | -5.695e+02 | 17.51% | 3.65% | 50.0bp (66.3bp) | NFY(CCAAT)/Promoter/Homer(0.972) More Information \| Similar Motifs Found | motif file (matrix) |
| 3 | C T A G A T G C A G T C G T A C A G T C C T A G A G T C A G T C A G T C G A T C A G C T A G T C | 1e-172 | -3.971e+02 | 24.98% | 9.45% | 53.1bp (66.1bp) | Sp1(Zf)/Promoter/Homer(0.946) More Information \| Similar Motifs Found | motif file (matrix) |
| 4 | A G T C A G T C C G T A A T G C G A C T C T G A A C T G C T A G C A T G A C T G A C T G G T A C | 1e-89 | -2.064e+02 | 5.35% | 0.89% | 48.3bp (63.6bp) | BORIS(Zf)/K562-CTCFL-ChIP-Seq(GSE32465)/Homer(0.931) More Information \| Similar Motifs Found | motif file (matrix) |
| 5 | A G T C A C G T T C A G T G A C C T A G G T A C G T C A A G C T A T C G G T A C A C T G G A T C | 1e-83 | -1.932e+02 | 8.17% | 2.21% | 51.4bp (73.0bp) | NRF1(NRF)/MCF7-NRF1-ChIP-Seq(Unpublished)/Homer(0.980) More Information \| Similar Motifs Found | motif file (matrix) |
| 6 | A T C G C G A T T C G A C T G A A G T C A G T C C G T A G T A C C G T A T C A G | 1e-71 | -1.639e+02 | 11.25% | 4.29% | 53.3bp (60.4bp) | RUNX(Runt)/HPC7-Runx1-ChIP-Seq(GSE22178)/Homer(0.972) More Information \| Similar Motifs Found | motif file (matrix) |
| 7 | C G T A T C G A T A G C T G A C C T A G T A C G T G C A C T G A T A C G A G C T | 1e-68 | -1.584e+02 | 12.70% | 5.29% | 55.9bp (64.7bp) | ELF1(ETS)/Jurkat-ELF1-ChIP-Seq(SRA014231)/Homer(0.983) More Information \| Similar Motifs Found | motif file (matrix) |
| 8 | T G C A T A C G G A C T T A G C T C G A G A T C C T A G G A C T T C A G T A C G | 1e-59 | -1.371e+02 | 9.49% | 3.62% | 52.4bp (65.3bp) | USF1/MA0093.2/Jaspar(0.916) More Information \| Similar Motifs Found | motif file (matrix) |
| 9 | C T A G C T A G A C T G C G T A T C G A C G A T A G C T T A C G C A G T C T G A A C T G A C G T | 1e-58 | -1.350e+02 | 3.26% | 0.50% | 52.3bp (60.5bp) | GFY(?)/Promoter/Homer(0.994) More Information \| Similar Motifs Found | motif file (matrix) |
| 10 | T C G A C T A G T C G A A C T G G A T C A C G T C A T G G C A T | 1e-39 | -9.054e+01 | 47.40% | 36.99% | 56.6bp (64.2bp) | MSC/MA0665.1/Jaspar(0.711) More Information \| Similar Motifs Found | motif file (matrix) |
| 11 | G A T C G A T C G T C A T C A G C G A T G T C A C G T A T C G A G T A C C T G A | 1e-34 | -8.034e+01 | 18.05% | 11.26% | 57.5bp (61.6bp) | FOXI1/MA0042.2/Jaspar(0.886) More Information \| Similar Motifs Found | motif file (matrix) |
| 12 | A C T G C A G T A G C T C A T G A G T C G A T C C T G A C A G T C T A G T A C G | 1e-34 | -7.956e+01 | 7.40% | 3.29% | 55.0bp (62.9bp) | Rfx5(HTH)/GM12878-Rfx5-ChIP-Seq(GSE31477)/Homer(0.922) More Information \| Similar Motifs Found | motif file (matrix) |
| 13 | G A T C T A G C T C G A T A G C C T G A T A C G T A C G T C G A T G C A C G A T | 1e-24 | -5.564e+01 | 10.27% | 5.98% | 54.6bp (60.3bp) | SPIB/MA0081.1/Jaspar(0.743) More Information \| Similar Motifs Found | motif file (matrix) |
| 14 | G A C T G T A C C G A T G T A C C T A G A G T C C T A G C G T A | 1e-20 | -4.613e+01 | 7.71% | 4.34% | 54.0bp (72.1bp) | ZBTB33/MA0527.1/Jaspar(0.875) More Information \| Similar Motifs Found | motif file (matrix) |
| 15 | A C G T A G T C A G T C A G T C C T G A A T C G C T G A C G T A | 1e-20 | -4.607e+01 | 8.25% | 4.76% | 55.8bp (58.6bp) | STAT3/MA0144.2/Jaspar(0.841) More Information \| Similar Motifs Found | motif file (matrix) |
| 16 | A T C G C G A T T A C G T C G A T A G C A T C G A G C T T A G C G T C A A C G T A T C G A T G C | 1e-18 | -4.320e+01 | 1.29% | 0.26% | 48.5bp (51.5bp) | JDP2(var.2)/MA0656.1/Jaspar(0.895) More Information \| Similar Motifs Found | motif file (matrix) |
| 17 | G C A T T A C G A C G T A T C G G T C A A G C T G C A T C G T A G C T A T A G C G T A C G T C A | 1e-15 | -3.679e+01 | 1.32% | 0.32% | 53.8bp (58.7bp) | PH0073.1\_Hoxc9/Jaspar(0.637) More Information \| Similar Motifs Found | motif file (matrix) |
| 18 | A C T G G A T C G T A C C T A G A G T C A G T C C G T A A C G T T G A C A G C T A C G T A C T G | 1e-15 | -3.655e+01 | 1.34% | 0.33% | 62.2bp (70.8bp) | YY1(Zf)/Promoter/Homer(0.988) More Information \| Similar Motifs Found | motif file (matrix) |
| 19 | C T A G G A T C G C T A C T A G C T A G G T A C C G A T A C T G A G T C A T G C C T G A C T A G | 1e-14 | -3.431e+01 | 0.78% | 0.11% | 57.8bp (52.1bp) | Tlx?(NR)/NPC-H3K4me1-ChIP-Seq(GSE16256)/Homer(0.808) More Information \| Similar Motifs Found | motif file (matrix) |
| 20 | T A C G G T C A A T C G T A G C A G C T T G C A A C T G A T G C | 1e-13 | -3.161e+01 | 10.47% | 7.13% | 57.0bp (67.6bp) | POL010.1\_DCE\_S\_III/Jaspar(0.634) More Information \| Similar Motifs Found | motif file (matrix) |
| 21 \* | T C G A G A T C C A G T G C T A G A T C C A T G G T C A G A T C G T C A C G A T G C T A A C T G | 1e-10 | -2.492e+01 | 6.90% | 4.51% | 55.0bp (67.4bp) | Crem/MA0609.1/Jaspar(0.793) More Information \| Similar Motifs Found | motif file (matrix) |
| 22 \* | A C G T A C G T A G C T C A T G T A C G G A T C T C A G G T A C A C T G A C G T T C G A T C G A | 1e-9 | -2.078e+01 | 0.18% | 0.01% | 49.5bp (36.3bp) | E2F8/MA0865.1/Jaspar(0.772) More Information \| Similar Motifs Found | motif file (matrix) |
| 23 \* | G T A C A C T G A C T G A C T G A G T C A C T G C G T A G T A C A C G T A T G C C G T A C T G A | 1e-7 | -1.697e+01 | 0.13% | 0.00% | 14.2bp (0.0bp) | PB0143.1\_Klf7\_2/Jaspar(0.609) More Information \| Similar Motifs Found | motif file (matrix) |
| 24 \* | A G T C C G T A A G T C A C G T C T G A A G T C C G T A A G T C C G T A A G T C C T G A G T A C | 1e-4 | -9.217e+00 | 0.13% | 0.01% | 41.6bp (54.3bp) | PB0130.1\_Gm397\_2/Jaspar(0.686) More Information \| Similar Motifs Found | motif file (matrix) |
| 25 \* | C G T A A C G T C T A G A C T G C G T A C G T A A C G T A C T G A C T G C G T A | 1e-2 | -5.037e+00 | 0.34% | 0.15% | 54.3bp (58.3bp) | PB0098.1\_Zfp410\_1/Jaspar(0.751) More Information \| Similar Motifs Found | motif file (matrix) |
