## Supplementary File 8.2 for "Promoter G-quadruplexes and transcription factors cooperate to shape the cell type-specific transcriptome": SupplementaryFile8.2_GS430_HomerMotifs_knownResults.html

GS430\_IP\_BG4\_93T\_rep2\_m1\_peaks\_MotifOutput/ - Homer Known Motif Enrichment Results


### Homer Known Motif Enrichment Results (GS430\_IP\_BG4\_93T\_rep2\_m1\_peaks\_MotifOutput/)

Homer *de novo* Motif Results  
Gene Ontology Enrichment Results  
Known Motif Enrichment Results (txt file)  
Total Target Sequences = 1013, Total Background Sequences = 47338

|  |  |  |  |  |  |  |  |  |  |  |  |
| --- | --- | --- | --- | --- | --- | --- | --- | --- | --- | --- | --- |
| Rank | Motif | Name | P-value | log P-pvalue | q-value (Benjamini) | # Target Sequences with Motif | % of Targets Sequences with Motif | # Background Sequences with Motif | % of Background Sequences with Motif | Motif File | SVG |
| 1 | C A T G C T A G T C G A A C G T A C T G C G T A T A G C C G A T T G A C C G T A A G C T G A T C | Fra2(bZIP)/Striatum-Fra2-ChIP-Seq(GSE43429)/Homer | 1e-130 | -3.013e+02 | 0.0000 | 263.0 | 25.96% | 1888.8 | 3.99% | motif file (matrix) | svg |
| 2 | A C T G C T A G T C G A C G A T C A T G G C T A A T C G C G A T G T A C G C T A A G C T G T A C | Fra1(bZIP)/BT549-Fra1-ChIP-Seq(GSE46166)/Homer | 1e-124 | -2.866e+02 | 0.0000 | 275.0 | 27.15% | 2234.6 | 4.72% | motif file (matrix) | svg |
| 3 | C T A G T C G A A C G T A C T G C G T A A T G C A C G T G T A C C G T A A G C T G A T C G T A C | Atf3(bZIP)/GBM-ATF3-ChIP-Seq(GSE33912)/Homer | 1e-124 | -2.857e+02 | 0.0000 | 298.0 | 29.42% | 2696.6 | 5.69% | motif file (matrix) | svg |
| 4 | C T A G T C G A G C A T C A T G G C T A T A G C C G A T G T A C C T G A A G C T | JunB(bZIP)/DendriticCells-Junb-ChIP-Seq(GSE36099)/Homer | 1e-123 | -2.844e+02 | 0.0000 | 275.0 | 27.15% | 2255.6 | 4.76% | motif file (matrix) | svg |
| 5 | C T A G T C G A C G A T A C T G C G T A T A C G A G C T T G A C G C T A A C G T G A T C T A G C | Fosl2(bZIP)/3T3L1-Fosl2-ChIP-Seq(GSE56872)/Homer | 1e-121 | -2.792e+02 | 0.0000 | 214.0 | 21.13% | 1257.9 | 2.66% | motif file (matrix) | svg |
| 6 | C A G T T G C A A C G T A C T G C G T A A T C G C G A T T G A C C G T A A C G T | BATF(bZIP)/Th17-BATF-ChIP-Seq(GSE39756)/Homer | 1e-116 | -2.681e+02 | 0.0000 | 288.0 | 28.43% | 2680.5 | 5.66% | motif file (matrix) | svg |
| 7 | T C G A A C G T C A T G G C T A T A G C C G A T G T A C G C T A A C G T A T G C | AP-1(bZIP)/ThioMac-PU.1-ChIP-Seq(GSE21512)/Homer | 1e-110 | -2.539e+02 | 0.0000 | 298.0 | 29.42% | 3059.4 | 6.46% | motif file (matrix) | svg |
| 8 | C T A G T C G A A C G T A C T G C G T A T A G C C G A T G T A C C G T A A G C T G A T C G T A C | Jun-AP1(bZIP)/K562-cJun-ChIP-Seq(GSE31477)/Homer | 1e-107 | -2.468e+02 | 0.0000 | 173.0 | 17.08% | 873.7 | 1.84% | motif file (matrix) | svg |
| 9 | C G A T T A C G T G A C G A C T C A T G C G T A T A C G A C G T G T A C C T G A | Bach2(bZIP)/OCILy7-Bach2-ChIP-Seq(GSE44420)/Homer | 1e-59 | -1.360e+02 | 0.0000 | 112.0 | 11.06% | 702.9 | 1.48% | motif file (matrix) | svg |
| 10 | T G C A A G C T C T G A A T C G G A C T C T A G G T A C G A T C G T C A A G T C G T A C G A C T C T A G A T C G G C A T C A T G C A T G G A T C G T A C C T G A | CTCF(Zf)/CD4+-CTCF-ChIP-Seq(Barski\_et\_al.)/Homer | 1e-32 | -7.488e+01 | 0.0000 | 61.0 | 6.02% | 376.3 | 0.79% | motif file (matrix) | svg |
| 11 | T A C G T C G A C A G T A C T G G C T A A T G C C G A T G T A C C G T A A C T G T A G C C G T A | NF-E2(bZIP)/K562-NFE2-ChIP-Seq(GSE31477)/Homer | 1e-29 | -6.743e+01 | 0.0000 | 43.0 | 4.24% | 183.0 | 0.39% | motif file (matrix) | svg |
| 12 | G T C A G C A T A C T G G T A C G A C T A C T G G C T A A T C G C A G T G T A C C G T A A G C T | Nrf2(bZIP)/Lymphoblast-Nrf2-ChIP-Seq(GSE37589)/Homer | 1e-25 | -5.970e+01 | 0.0000 | 38.0 | 3.75% | 162.8 | 0.34% | motif file (matrix) | svg |
| 13 | T A G C A G T C T G A C A G T C C T A G A T C G A G T C C A T G T G A C A G T C G T A C A G T C A G T C G C A T C T A G A T C G G C A T A C T G A T C G G A T C | BORIS(Zf)/K562-CTCFL-ChIP-Seq(GSE32465)/Homer | 1e-23 | -5.482e+01 | 0.0000 | 72.0 | 7.11% | 773.7 | 1.63% | motif file (matrix) | svg |
| 14 | C G T A C G T A C G T A G C A T G C A T T A C G G T A C G A C T A C T G C G T A A T C G A C G T G T A C C G T A A G C T | Bach1(bZIP)/K562-Bach1-ChIP-Seq(GSE31477)/Homer | 1e-22 | -5.210e+01 | 0.0000 | 35.0 | 3.46% | 164.5 | 0.35% | motif file (matrix) | svg |
| 15 | A G T C C T G A A G T C C G A T C A G T G A T C A T G C A C T G A T C G G A C T | Fli1(ETS)/CD8-FLI-ChIP-Seq(GSE20898)/Homer | 1e-18 | -4.168e+01 | 0.0000 | 208.0 | 20.53% | 5199.3 | 10.98% | motif file (matrix) | svg |
| 16 | T C G A C T A G A G T C A G T C C G T A C G T A A C G T T A G C T C A G T A C G | NFY(CCAAT)/Promoter/Homer | 1e-17 | -4.139e+01 | 0.0000 | 153.0 | 15.10% | 3341.5 | 7.05% | motif file (matrix) | svg |
| 17 | T C A G T A G C G A C T C A T G C T G A A T C G G C A T G T A C C G T A A C T G T A G C T G C A | MafK(bZIP)/C2C12-MafK-ChIP-Seq(GSE36030)/Homer | 1e-16 | -3.813e+01 | 0.0000 | 64.0 | 6.32% | 856.9 | 1.81% | motif file (matrix) | svg |
| 18 | G C T A C T G A T C G A A G T C A G T C C T G A A G T C G T C A C T G A T G C A | RUNX1(Runt)/Jurkat-RUNX1-ChIP-Seq(GSE29180)/Homer | 1e-15 | -3.607e+01 | 0.0000 | 172.0 | 16.98% | 4201.3 | 8.87% | motif file (matrix) | svg |
| 19 | G A T C T C G A A G T C C G A T C G A T A G T C A T G C A C T G A T C G G A C T | Elk1(ETS)/Hela-Elk1-ChIP-Seq(GSE31477)/Homer | 1e-14 | -3.394e+01 | 0.0000 | 124.0 | 12.24% | 2686.3 | 5.67% | motif file (matrix) | svg |
| 20 | T C G A T A G C T G C A A C T G A C T G C G T A C G T A C T A G G A C T T A C G | ETS1(ETS)/Jurkat-ETS1-ChIP-Seq(GSE17954)/Homer | 1e-14 | -3.237e+01 | 0.0000 | 182.0 | 17.97% | 4735.1 | 10.00% | motif file (matrix) | svg |
| 21 | T A C G C T A G A T G C G A T C G T A C A G T C C T A G A G T C A G T C A G T C G T A C A G T C | Sp1(Zf)/Promoter/Homer | 1e-13 | -3.131e+01 | 0.0000 | 81.0 | 8.00% | 1461.1 | 3.08% | motif file (matrix) | svg |
| 22 | G A C T C T A G G A T C C A G T A C T G C T G A A T G C G C A T A T G C C T G A | MafA(bZIP)/Islet-MafA-ChIP-Seq(GSE30298)/Homer | 1e-13 | -3.011e+01 | 0.0000 | 135.0 | 13.33% | 3205.8 | 6.77% | motif file (matrix) | svg |
| 23 | A T G C A G T C C T G A A G T C C G A T A C G T A G T C A G T C A C G T A T C G G A C T A C G T | Etv2(ETS)/ES-ER71-ChIP-Seq(GSE59402)/Homer(0.967) | 1e-13 | -2.996e+01 | 0.0000 | 159.0 | 15.70% | 4041.9 | 8.53% | motif file (matrix) | svg |
| 24 | T C G A C T G A T A G C T G A C T C A G T C A G C G T A C G T A T C A G A G C T | ETV1(ETS)/GIST48-ETV1-ChIP-Seq(GSE22441)/Homer | 1e-12 | -2.981e+01 | 0.0000 | 218.0 | 21.52% | 6210.2 | 13.11% | motif file (matrix) | svg |
| 25 | C T G A T G C A T A G C T G A C T A C G T C A G C T G A G C T A T C A G G A C T | ELF1(ETS)/Jurkat-ELF1-ChIP-Seq(SRA014231)/Homer | 1e-12 | -2.907e+01 | 0.0000 | 112.0 | 11.06% | 2487.6 | 5.25% | motif file (matrix) | svg |
| 26 | G A T C C T G A A G T C C G A T C G A T G A T C A G T C A C T G A T C G A G C T | Elk4(ETS)/Hela-Elk4-ChIP-Seq(GSE31477)/Homer | 1e-10 | -2.530e+01 | 0.0000 | 112.0 | 11.06% | 2645.3 | 5.59% | motif file (matrix) | svg |
| 27 | T C G A T C G A T A G C G T A C T C A G T A C G C G T A C G T A T C A G A G C T | GABPA(ETS)/Jurkat-GABPa-ChIP-Seq(GSE17954)/Homer | 1e-9 | -2.281e+01 | 0.0000 | 149.0 | 14.71% | 4071.9 | 8.60% | motif file (matrix) | svg |
| 28 | C T G A T C A G C A G T C T A G A C T G C T A G G A T C A T C G A C T G C T G A T C A G G A T C | Sp5(Zf)/mES-Sp5.Flag-ChIP-Seq(GSE72989)/Homer | 1e-9 | -2.269e+01 | 0.0000 | 175.0 | 17.28% | 5043.9 | 10.65% | motif file (matrix) | svg |
| 29 | T C G A T A G C G T C A A C T G A C T G C G T A C G T A C T A G A G C T T C A G | ERG(ETS)/VCaP-ERG-ChIP-Seq(GSE14097)/Homer | 1e-9 | -2.217e+01 | 0.0000 | 226.0 | 22.31% | 7053.3 | 14.89% | motif file (matrix) | svg |
| 30 | T A G C G C T A T C G A C T G A A G T C A G T C C T G A A G T C C G T A C T A G | RUNX(Runt)/HPC7-Runx1-ChIP-Seq(GSE22178)/Homer | 1e-8 | -2.064e+01 | 0.0000 | 112.0 | 11.06% | 2869.4 | 6.06% | motif file (matrix) | svg |
| 31 | T G C A T C G A T A G C G T A C T C A G C T A G G T C A G C T A T C A G G A C T | ETS(ETS)/Promoter/Homer | 1e-8 | -1.968e+01 | 0.0000 | 72.0 | 7.11% | 1571.8 | 3.32% | motif file (matrix) | svg |
| 32 | C T A G T C A G C A G T T C A G A C T G A C T G G A T C C T A G A C T G C T A G T C A G A T G C | KLF14(Zf)/HEK293-KLF14.GFP-ChIP-Seq(GSE58341)/Homer | 1e-7 | -1.787e+01 | 0.0000 | 255.0 | 25.17% | 8600.1 | 18.16% | motif file (matrix) | svg |
| 33 | C G T A G A C T C G A T A T C G G T A C G C A T C A T G C G T A T A C G G C A T G T A C C G T A C A T G A T G C G C T A C T A G G C A T G C A T G C A T G A C T | MafB(bZIP)/BMM-Mafb-ChIP-Seq(GSE75722)/Homer | 1e-7 | -1.764e+01 | 0.0000 | 72.0 | 7.11% | 1654.3 | 3.49% | motif file (matrix) | svg |
| 34 | A T G C A G C T T C A G T G A C T C A G A T G C T G C A A C G T A T C G G A T C A C T G A G T C | NRF1(NRF)/MCF7-NRF1-ChIP-Seq(Unpublished)/Homer | 1e-7 | -1.750e+01 | 0.0000 | 41.0 | 4.05% | 712.2 | 1.50% | motif file (matrix) | svg |
| 35 | G C T A A G T C T A C G T G C A A T C G T C A G G C T A T C G A T C A G A G C T | ELF5(ETS)/T47D-ELF5-ChIP-Seq(GSE30407)/Homer | 1e-7 | -1.729e+01 | 0.0000 | 114.0 | 11.25% | 3128.2 | 6.60% | motif file (matrix) | svg |
| 36 | T A G C T A G C G C A T C A T G A C T G G C T A C G T A A C G T A C T G G A T C | TEAD4(TEA)/Tropoblast-Tead4-ChIP-Seq(GSE37350)/Homer | 1e-7 | -1.666e+01 | 0.0000 | 126.0 | 12.44% | 3609.7 | 7.62% | motif file (matrix) | svg |
| 37 | C T G A A T G C C G T A A C G T A G T C A G T C A C G T A C T G A T C G G C A T | SPDEF(ETS)/VCaP-SPDEF-ChIP-Seq(SRA014231)/Homer | 1e-6 | -1.601e+01 | 0.0000 | 142.0 | 14.02% | 4257.2 | 8.99% | motif file (matrix) | svg |
| 38 | C G T A T A G C T A G C T G C A A C T G C T A G C G T A C G T A T C A G G A C T | EHF(ETS)/LoVo-EHF-ChIP-Seq(GSE49402)/Homer | 1e-6 | -1.500e+01 | 0.0000 | 178.0 | 17.57% | 5744.9 | 12.13% | motif file (matrix) | svg |
| 39 | A T G C A T C G T A C G A G C T A T C G C T G A A G T C C T A G A G C T A T G C C T G A A T G C | CRE(bZIP)/Promoter/Homer | 1e-6 | -1.479e+01 | 0.0000 | 47.0 | 4.64% | 971.9 | 2.05% | motif file (matrix) | svg |
| 40 | T G C A C T G A A G T C G T C A A C T G A C T G C G T A C G T A C T G A A G C T | EWS:FLI1-fusion(ETS)/SK\_N\_MC-EWS:FLI1-ChIP-Seq(SRA014231)/Homer | 1e-6 | -1.436e+01 | 0.0000 | 91.0 | 8.98% | 2474.6 | 5.22% | motif file (matrix) | svg |
| 41 | G A T C G T A C C G A T A C T G A C T G C G T A C G T A A C G T A C T G G A T C | TEAD(TEA)/Fibroblast-PU.1-ChIP-Seq(Unpublished)/Homer | 1e-6 | -1.400e+01 | 0.0000 | 102.0 | 10.07% | 2897.3 | 6.12% | motif file (matrix) | svg |
| 42 | G T A C C A G T A C T G A C T G A C T G G A T C A C T G A C G T A C T G A C T G A G T C G A T C | KLF6(Zf)/PDAC-KLF6-ChIP-Seq(GSE64557)/Homer | 1e-6 | -1.396e+01 | 0.0000 | 157.0 | 15.50% | 5003.2 | 10.56% | motif file (matrix) | svg |
| 43 | C G T A C T A G A C T G A C T G G A C T C T A G C A G T C T A G C A T G G A T C | KLF5(Zf)/LoVo-KLF5-ChIP-Seq(GSE49402)/Homer | 1e-6 | -1.396e+01 | 0.0000 | 195.0 | 19.25% | 6525.5 | 13.78% | motif file (matrix) | svg |
| 44 | G A C T T C A G C T A G A G T C A G T C G T A C A G T C C T G A A G T C A G T C A G T C G A C T A G T C A C T G A T G C | KLF3(Zf)/MEF-Klf3-ChIP-Seq(GSE44748)/Homer | 1e-5 | -1.380e+01 | 0.0000 | 91.0 | 8.98% | 2508.6 | 5.30% | motif file (matrix) | svg |
| 45 | T G A C G C T A T C G A T G C A A G T C A G T C C G T A A G T C C G T A C T G A G C T A G T A C | RUNX2(Runt)/PCa-RUNX2-ChIP-Seq(GSE33889)/Homer | 1e-5 | -1.352e+01 | 0.0000 | 115.0 | 11.35% | 3415.7 | 7.21% | motif file (matrix) | svg |
| 46 | C T A G T C G A C G A T C T A G G C A T C A G T C T A G G A T C C G T A G T C A | CEBP:AP1(bZIP)/ThioMac-CEBPb-ChIP-Seq(GSE21512)/Homer | 1e-5 | -1.265e+01 | 0.0000 | 108.0 | 10.66% | 3215.9 | 6.79% | motif file (matrix) | svg |
| 47 | A T G C A T G C A T C G T A C G A G C T A G T C G C T A A G T C T C A G G A C T A C T G T C G A | E-box(bHLH)/Promoter/Homer | 1e-5 | -1.233e+01 | 0.0000 | 21.0 | 2.07% | 304.8 | 0.64% | motif file (matrix) | svg |
| 48 | A T G C T C G A A G T C A G C T A C G T G T A C A G T C G C T A C T A G C A T G G T C A C T G A T C A G A G T C | Stat3+il21(Stat)/CD4-Stat3-ChIP-Seq(GSE19198)/Homer | 1e-5 | -1.200e+01 | 0.0000 | 92.0 | 9.08% | 2662.2 | 5.62% | motif file (matrix) | svg |
| 49 | A G T C C G A T A C T G A T C G T G A C G C T A C A T G A T C G T G A C C G A T A C T G T A G C G T A C G T C A | Tlx?(NR)/NPC-H3K4me1-ChIP-Seq(GSE16256)/Homer | 1e-5 | -1.161e+01 | 0.0001 | 62.0 | 6.12% | 1604.6 | 3.39% | motif file (matrix) | svg |
| 50 | T C A G A G C T A T G C C G T A A G C T T C A G C A G T A C T G C T G A A G T C | MITF(bHLH)/MastCells-MITF-ChIP-Seq(GSE48085)/Homer | 1e-4 | -1.129e+01 | 0.0001 | 128.0 | 12.64% | 4096.3 | 8.65% | motif file (matrix) | svg |
| 51 | C T A G A G C T G A C T C A T G A G T C A G T C G T C A C A G T C T A G T C A G G T A C C T G A T C G A G A T C T G A C | Rfx2(HTH)/LoVo-RFX2-ChIP-Seq(GSE49402)/Homer | 1e-4 | -1.107e+01 | 0.0001 | 21.0 | 2.07% | 331.8 | 0.70% | motif file (matrix) | svg |
| 52 | A C T G C A T G G C T A T C G A G C T A A G C T A G C T G T A C A G T C T G A C | NFkB-p65-Rel(RHD)/ThioMac-LPS-Expression(GSE23622)/Homer | 1e-4 | -1.004e+01 | 0.0003 | 14.0 | 1.38% | 178.6 | 0.38% | motif file (matrix) | svg |
| 53 | T C A G C G T A A G T C A G C T C G T A A G T C C T G A C G T A A G T C G C A T A G T C A G T C A G T C C T G A A C T G T G C A T C G A C A T G A T C G G A T C | Ronin(THAP)/ES-Thap11-ChIP-Seq(GSE51522)/Homer | 1e-4 | -1.004e+01 | 0.0003 | 14.0 | 1.38% | 178.5 | 0.38% | motif file (matrix) | svg |
| 54 | A C T G G A T C G A C T A C T G A C G T C A T G A C T G A C G T A G C T C G A T | RUNX-AML(Runt)/CD4+-PolII-ChIP-Seq(Barski\_et\_al.)/Homer | 1e-4 | -9.818e+00 | 0.0004 | 97.0 | 9.58% | 3016.4 | 6.37% | motif file (matrix) | svg |
| 55 | A T C G A G C T A C T G A G T C A C T G A G T C C G T A A C G T A C T G A G T C A C T G A G T C | NRF(NRF)/Promoter/Homer | 1e-4 | -9.799e+00 | 0.0004 | 39.0 | 3.85% | 913.3 | 1.93% | motif file (matrix) | svg |
| 56 | C T A G A C T G C T A G T C A G T C A G T A C G C T A G A C T G | Maz(Zf)/HepG2-Maz-ChIP-Seq(GSE31477)/Homer | 1e-4 | -9.540e+00 | 0.0005 | 164.0 | 16.19% | 5725.1 | 12.09% | motif file (matrix) | svg |
| 57 | T C A G A G C T A C G T A C G T G T A C G A T C C G T A C T A G C A T G G T C A C G T A T C G A | STAT4(Stat)/CD4-Stat4-ChIP-Seq(GSE22104)/Homer | 1e-3 | -9.166e+00 | 0.0007 | 116.0 | 11.45% | 3819.9 | 8.06% | motif file (matrix) | svg |
| 58 | T C G A A G C T A C G T A C G T A G T C A G T C A C G T A T C G G A C T A T C G | EWS:ERG-fusion(ETS)/CADO\_ES1-EWS:ERG-ChIP-Seq(SRA014231)/Homer | 1e-3 | -8.954e+00 | 0.0008 | 96.0 | 9.48% | 3051.8 | 6.44% | motif file (matrix) | svg |
| 59 | A T G C G A C T A C G T C T A G A C G T A C G T A C G T C T G A G A T C G C T A A G C T C G T A | Foxa2(Forkhead)/Liver-Foxa2-ChIP-Seq(GSE25694)/Homer | 1e-3 | -8.733e+00 | 0.0010 | 102.0 | 10.07% | 3306.2 | 6.98% | motif file (matrix) | svg |
| 60 | A G T C G A C T C A G T A C T G C T A G T G A C G C T A A T G C G C A T A T C G C G A T A C T G G A T C G T A C G T C A C T G A | NF1(CTF)/LNCAP-NF1-ChIP-Seq(Unpublished)/Homer | 1e-3 | -8.690e+00 | 0.0010 | 58.0 | 5.73% | 1634.5 | 3.45% | motif file (matrix) | svg |
| 61 | A G T C G A C T C A G T G T A C A G T C A T C G T C A G A C T G G T C A C G T A | Stat3(Stat)/mES-Stat3-ChIP-Seq(GSE11431)/Homer | 1e-3 | -8.651e+00 | 0.0010 | 66.0 | 6.52% | 1931.6 | 4.08% | motif file (matrix) | svg |
| 62 | A G T C A T C G C T A G A G C T G A C T C T A G A G T C A G T C G C T A C A G T T C A G T C A G G A T C C T G A T C G A G A T C | RFX(HTH)/K562-RFX3-ChIP-Seq(SRA012198)/Homer | 1e-3 | -8.527e+00 | 0.0012 | 17.0 | 1.68% | 285.7 | 0.60% | motif file (matrix) | svg |
| 63 | C T A G T A C G G A T C G T A C G C T A A G C T A G C T G T C A T C G A T A G C | Nanog(Homeobox)/mES-Nanog-ChIP-Seq(GSE11724)/Homer | 1e-3 | -8.282e+00 | 0.0015 | 472.0 | 46.59% | 19486.6 | 41.14% | motif file (matrix) | svg |
| 64 | T C A G T G A C G T A C T G C A G T A C C T A G G T A C A T G C A G T C G T C A A G T C G A C T | Klf9(Zf)/GBM-Klf9-ChIP-Seq(GSE62211)/Homer | 1e-3 | -8.208e+00 | 0.0015 | 58.0 | 5.73% | 1666.3 | 3.52% | motif file (matrix) | svg |
| 65 | A G T C A C G T A C T G A G C T A C G T A C G T G T C A A G T C | Foxo1(Forkhead)/RAW-Foxo1-ChIP-Seq(Fan\_et\_al.)/Homer | 1e-3 | -8.159e+00 | 0.0016 | 223.0 | 22.01% | 8394.4 | 17.72% | motif file (matrix) | svg |
| 66 | T A C G T C A G A G C T A T G C C G T A A G T C T C A G A C G T A C T G T C G A | USF1(bHLH)/GM12878-Usf1-ChIP-Seq(GSE32465)/Homer | 1e-3 | -8.122e+00 | 0.0016 | 65.0 | 6.42% | 1932.5 | 4.08% | motif file (matrix) | svg |
| 67 | A T G C A G T C G C A T A G C T A C G T T C A G C G A T A G C T G A T C A T C G | Sox10(HMG)/SciaticNerve-Sox3-ChIP-Seq(GSE35132)/Homer | 1e-3 | -7.931e+00 | 0.0020 | 195.0 | 19.25% | 7230.7 | 15.27% | motif file (matrix) | svg |
| 68 | G T C A T G C A G C T A A G T C C G T A A C T G T G A C G C A T T C A G C A G T | Ap4(bHLH)/AML-Tfap4-ChIP-Seq(GSE45738)/Homer | 1e-3 | -7.905e+00 | 0.0020 | 149.0 | 14.71% | 5300.6 | 11.19% | motif file (matrix) | svg |
| 69 | T G C A C T G A A T G C G T C A A C T G A C T G C G T A C G T A C T A G A G C T | Ets1-distal(ETS)/CD4+-PolII-ChIP-Seq(Barski\_et\_al.)/Homer | 1e-3 | -7.742e+00 | 0.0023 | 44.0 | 4.34% | 1189.5 | 2.51% | motif file (matrix) | svg |
| 70 | A T G C G A T C C G A T C T A G A C T G G C T A C G T A A G C T A C T G A G C T | TEAD2(TEA)/Py2T-Tead2-ChIP-Seq(GSE55709)/Homer | 1e-3 | -7.546e+00 | 0.0027 | 72.0 | 7.11% | 2243.3 | 4.74% | motif file (matrix) | svg |
| 71 | C G T A T G A C T A G C T G C A A C T G A C T G C G T A C G T A T C A G G A C T | ELF3(ETS)/PDAC-ELF3-ChIP-Seq(GSE64557)/Homer | 1e-3 | -7.298e+00 | 0.0035 | 93.0 | 9.18% | 3088.9 | 6.52% | motif file (matrix) | svg |
| 72 | T A C G T C A G T G C A A G C T T G A C A G C T A G T C A C T G G A T C A C T G T C G A A C T G C T G A C T G A A T G C | ZBTB33(Zf)/GM12878-ZBTB33-ChIP-Seq(GSE32465)/Homer | 1e-3 | -7.257e+00 | 0.0036 | 12.0 | 1.18% | 182.9 | 0.39% | motif file (matrix) | svg |
| 73 | C A T G G T A C C G T A A G T C C T A G A C G T A C T G G T A C A G T C A G C T | bHLHE40(bHLH)/HepG2-BHLHE40-ChIP-Seq(GSE31477)/Homer | 1e-3 | -7.034e+00 | 0.0044 | 46.0 | 4.54% | 1305.3 | 2.76% | motif file (matrix) | svg |
| 74 | A G C T G C A T A C T G A C G T A G T C A C G T C T A G T A C G | Smad3(MAD)/NPC-Smad3-ChIP-Seq(GSE36673)/Homer | 1e-2 | -6.835e+00 | 0.0053 | 319.0 | 31.49% | 12834.4 | 27.10% | motif file (matrix) | svg |
| 75 | T C G A G A C T A G C T T G A C A G C T G T A C T C A G G A T C A T C G T G C A A C T G C T G A | GFX(?)/Promoter/Homer | 1e-2 | -6.769e+00 | 0.0056 | 7.0 | 0.69% | 73.3 | 0.15% | motif file (matrix) | svg |
| 76 | T A G C C G A T A C G T A G C T A G C T A G T C A T G C A G T C A C T G A T G C A T G C G C T A | E2F7(E2F)/Hela-E2F7-ChIP-Seq(GSE32673)/Homer | 1e-2 | -6.751e+00 | 0.0056 | 21.0 | 2.07% | 459.0 | 0.97% | motif file (matrix) | svg |
| 77 | A T G C C T G A G A C T A C G T A C G T G T A C G A T C C G A T C T A G C A T G C G T A C G T A C T G A G A C T | STAT1(Stat)/HelaS3-STAT1-ChIP-Seq(GSE12782)/Homer | 1e-2 | -6.748e+00 | 0.0056 | 39.0 | 3.85% | 1070.6 | 2.26% | motif file (matrix) | svg |
| 78 | C T G A T C A G G T A C G C T A A C T G T G A C G C A T C A T G | SCL(bHLH)/HPC7-Scl-ChIP-Seq(GSE13511)/Homer | 1e-2 | -6.635e+00 | 0.0061 | 488.0 | 48.17% | 20568.6 | 43.43% | motif file (matrix) | svg |
| 79 | T C G A C G T A A G T C A G C T C G T A A G T C T C G A G C T A G A C T C G A T A G T C A G T C A G T C C T G A T C A G T G C A T C G A C A G T A T C G A G T C | GFY-Staf(?,Zf)/Promoter/Homer | 1e-2 | -6.559e+00 | 0.0065 | 15.0 | 1.48% | 282.0 | 0.60% | motif file (matrix) | svg |
| 80 | A G C T A C G T A C T G A T G C A G T C C G T A C T G A T A C G | NF1-halfsite(CTF)/LNCaP-NF1-ChIP-Seq(Unpublished)/Homer | 1e-2 | -6.532e+00 | 0.0066 | 204.0 | 20.14% | 7830.1 | 16.53% | motif file (matrix) | svg |
| 81 | C T A G G T A C A G T C T G C A A G T C C T G A A G T C A G T C A G T C G C T A | Klf4(Zf)/mES-Klf4-ChIP-Seq(GSE11431)/Homer | 1e-2 | -6.397e+00 | 0.0075 | 60.0 | 5.92% | 1878.9 | 3.97% | motif file (matrix) | svg |
| 82 | T C A G G A C T C A G T C T G A A G C T C T A G G A C T T G C A C T G A A G T C | HLF(bZIP)/HSC-HLF.Flag-ChIP-Seq(GSE69817)/Homer | 1e-2 | -6.341e+00 | 0.0078 | 104.0 | 10.27% | 3636.4 | 7.68% | motif file (matrix) | svg |
| 83 | G A C T C G A T T C A G G A T C G A C T A G C T A G C T A G T C G A T C C G T A C T A G C T A G T C G A T C G A C T G A | Bcl6(Zf)/Liver-Bcl6-ChIP-Seq(GSE31578)/Homer | 1e-2 | -6.308e+00 | 0.0080 | 143.0 | 14.12% | 5260.0 | 11.11% | motif file (matrix) | svg |
| 84 | A G C T A G T C A G T C A C G T C T A G A C G T A C G T A C G T C G T A A G T C G A T C C G T A | FOXP1(Forkhead)/H9-FOXP1-ChIP-Seq(GSE31006)/Homer | 1e-2 | -6.174e+00 | 0.0090 | 56.0 | 5.53% | 1743.5 | 3.68% | motif file (matrix) | svg |
| 85 | C A G T T C A G G A T C A C T G A C G T C T A G A C T G A C T G G A C T C T A G | Egr1(Zf)/K562-Egr1-ChIP-Seq(GSE32465)/Homer | 1e-2 | -6.148e+00 | 0.0092 | 85.0 | 8.39% | 2887.0 | 6.10% | motif file (matrix) | svg |
| 86 | C T G A C A G T C T G A A G T C C T A G G A C T A T C G G T A C | HIF-1b(HLH)/T47D-HIF1b-ChIP-Seq(GSE59937)/Homer | 1e-2 | -6.025e+00 | 0.0102 | 140.0 | 13.82% | 5174.2 | 10.92% | motif file (matrix) | svg |
| 87 | T C G A A C G T A C T G C T G A A G T C T C A G A G C T G T A C C G T A A G C T G A T C T C G A | JunD(bZIP)/K562-JunD-ChIP-Seq/Homer | 1e-2 | -6.013e+00 | 0.0102 | 17.0 | 1.68% | 361.0 | 0.76% | motif file (matrix) | svg |
| 88 | T G C A T A G C G A C T T G C A T G A C T G C A C G T A A G C T A G C T A G T C A G T C G T A C | GFY(?)/Promoter/Homer | 1e-2 | -5.959e+00 | 0.0107 | 16.0 | 1.58% | 331.6 | 0.70% | motif file (matrix) | svg |
| 89 | T C G A T A G C G T C A A C T G C T A G C G T A C G A T A C T G A C G T A C T G A C T G A C G T | ETS:RUNX(ETS,Runt)/Jurkat-RUNX1-ChIP-Seq(GSE17954)/Homer | 1e-2 | -5.941e+00 | 0.0108 | 18.0 | 1.78% | 394.9 | 0.83% | motif file (matrix) | svg |
| 90 | C T A G C A T G T G C A A C G T A G T C C G T A C A T G T C A G A C G T A C G T G C T A A G T C | Six1(Homeobox)/Myoblast-Six1-ChIP-Chip(GSE20150)/Homer | 1e-2 | -5.641e+00 | 0.0144 | 35.0 | 3.46% | 995.5 | 2.10% | motif file (matrix) | svg |
| 91 | T C A G A C G T A G T C T C G A A G T C T C A G G C A T C T A G C T A G A G C T | Usf2(bHLH)/C2C12-Usf2-ChIP-Seq(GSE36030)/Homer | 1e-2 | -5.635e+00 | 0.0144 | 51.0 | 5.03% | 1596.1 | 3.37% | motif file (matrix) | svg |
| 92 | T G C A A G C T A C G T C T A G G A T C C T A G G A T C G T C A C T G A A G T C | CEBP(bZIP)/ThioMac-CEBPb-ChIP-Seq(GSE21512)/Homer | 1e-2 | -5.524e+00 | 0.0158 | 98.0 | 9.67% | 3490.0 | 7.37% | motif file (matrix) | svg |
| 93 | T C A G A G C T A T G C C G T A A G T C T C A G A C G T A T C G T C G A A G T C G A T C T G A C | TFE3(bHLH)/MEF-TFE3-ChIP-Seq(GSE75757)/Homer | 1e-2 | -5.477e+00 | 0.0164 | 15.0 | 1.48% | 317.8 | 0.67% | motif file (matrix) | svg |
| 94 | A T G C G A T C C G A T A C G T A C G T A C T G C A G T A G C T | Sox3(HMG)/NPC-Sox3-ChIP-Seq(GSE33059)/Homer | 1e-2 | -5.435e+00 | 0.0169 | 196.0 | 19.35% | 7673.3 | 16.20% | motif file (matrix) | svg |
| 95 | T G A C C G A T A C T G A C T G A C T G G A C T A C T G A C G T A C T G A C T G G A T C G A T C | EKLF(Zf)/Erythrocyte-Klf1-ChIP-Seq(GSE20478)/Homer | 1e-2 | -5.328e+00 | 0.0186 | 32.0 | 3.16% | 906.9 | 1.91% | motif file (matrix) | svg |
| 96 | C G T A T A C G T C G A A C T G A C T G C G T A C G T A T A C G A G C T T A C G | PU.1(ETS)/ThioMac-PU.1-ChIP-Seq(GSE21512)/Homer | 1e-2 | -5.102e+00 | 0.0231 | 62.0 | 6.12% | 2076.9 | 4.38% | motif file (matrix) | svg |
| 97 | C G T A C A T G C A T G A C T G C T A G T C G A G C A T C G A T A G C T A G T C G A T C G T A C | NFkB-p65(RHD)/GM12787-p65-ChIP-Seq(GSE19485)/Homer | 1e-2 | -4.926e+00 | 0.0272 | 57.0 | 5.63% | 1896.0 | 4.00% | motif file (matrix) | svg |
| 98 | G C T A C G A T C A T G A T G C A G T C A G T C G A C T T A C G T C G A C T A G A C T G T A G C | AP-2alpha(AP2)/Hela-AP2alpha-ChIP-Seq(GSE31477)/Homer | 1e-2 | -4.923e+00 | 0.0272 | 100.0 | 9.87% | 3651.3 | 7.71% | motif file (matrix) | svg |
| 99 | T C G A T G A C A G T C C G T A A G T C C T A G A C G T A C T G A C T G A G C T A G T C G C A T | Max(bHLH)/K562-Max-ChIP-Seq(GSE31477)/Homer | 1e-2 | -4.838e+00 | 0.0291 | 80.0 | 7.90% | 2834.9 | 5.99% | motif file (matrix) | svg |
| 100 | T C G A G C A T A C T G C T G A A G T C T C A G G A C T G T A C C G T A A G C T A G T C G A T C | c-Jun-CRE(bZIP)/K562-cJun-ChIP-Seq(GSE31477)/Homer | 1e-2 | -4.757e+00 | 0.0313 | 50.0 | 4.94% | 1635.1 | 3.45% | motif file (matrix) | svg |
| 101 | G C T A T C G A C G T A C T A G A G C T G T C A G T C A C G T A A G T C C G T A | FOXA1(Forkhead)/LNCAP-FOXA1-ChIP-Seq(GSE27824)/Homer | 1e-2 | -4.727e+00 | 0.0319 | 143.0 | 14.12% | 5505.8 | 11.62% | motif file (matrix) | svg |
| 102 | T A C G T A C G G T A C A T C G A C T G T A C G T C G A C T G A T C G A A T C G | E2F6(E2F)/Hela-E2F6-ChIP-Seq(GSE31477)/Homer | 1e-2 | -4.679e+00 | 0.0332 | 66.0 | 6.52% | 2282.5 | 4.82% | motif file (matrix) | svg |
