## Supplementary File 8.2.1 for "Promoter G-quadruplexes and transcription factors cooperate to shape the cell type-specific transcriptome": SupplementaryFile8.2.1_GS430_homerResults_denovoMotifs.html

GS430\_IP\_BG4\_93T\_rep2\_m1\_F2L0\_peaks\_MotifOutput// - Homer de novo Motif Results


### Homer *de novo* Motif Results (GS430\_IP\_BG4\_93T\_rep2\_m1\_F2L0\_peaks\_MotifOutput//)

Known Motif Enrichment Results  
Gene Ontology Enrichment Results  
If Homer is having trouble matching a motif to a known motif, try copy/pasting the matrix file into
STAMP  
More information on motif finding results: HOMER
| Description of Results
| Tips
  
Total target sequences = 1334  
Total background sequences = 47678  
\* - possible false positive  

|  |  |  |  |  |  |  |  |  |
| --- | --- | --- | --- | --- | --- | --- | --- | --- |
| Rank | Motif | P-value | log P-pvalue | % of Targets | % of Background | STD(Bg STD) | Best Match/Details | Motif File |
| 1 | T A G C T G A C T C G A A C G T A C T G C G T A A T C G A C G T A G T C C G T A | 1e-83 | -1.926e+02 | 15.37% | 2.86% | 48.2bp (63.6bp) | FOS/MA0476.1/Jaspar(0.961) More Information \| Similar Motifs Found | motif file (matrix) |
| 2 | C A T G C T A G C A T G A C T G G T A C C T A G A C T G C A T G T A C G G T A C G A C T T C G A | 1e-22 | -5.219e+01 | 6.07% | 1.61% | 49.7bp (62.1bp) | Sp1(Zf)/Promoter/Homer(0.951) More Information \| Similar Motifs Found | motif file (matrix) |
| 3 | C T A G A G T C A G T C C G T A C G T A A C G T T G A C T C A G | 1e-20 | -4.745e+01 | 11.17% | 4.76% | 50.2bp (63.5bp) | NFY(CCAAT)/Promoter/Homer(0.909) More Information \| Similar Motifs Found | motif file (matrix) |
| 4 | A G T C C A G T G A C T G T A C T G A C A T C G T A C G A G T C G T C A A G C T | 1e-18 | -4.329e+01 | 5.77% | 1.71% | 53.4bp (64.0bp) | Elk1(ETS)/Hela-Elk1-ChIP-Seq(GSE31477)/Homer(0.879) More Information \| Similar Motifs Found | motif file (matrix) |
| 5 | G T A C G T A C C T G A T A G C G C T A T C G A A T C G T C G A A C T G A C T G A C T G T G A C | 1e-17 | -4.015e+01 | 2.70% | 0.41% | 45.7bp (58.9bp) | BORIS(Zf)/K562-CTCFL-ChIP-Seq(GSE32465)/Homer(0.872) More Information \| Similar Motifs Found | motif file (matrix) |
| 6 | A C T G C G A T C G T A C G T A A C T G A T G C C G T A C T G A A C G T A C T G C T G A A G T C | 1e-14 | -3.361e+01 | 0.52% | 0.00% | 43.7bp (0.0bp) | FOXO3/MA0157.2/Jaspar(0.640) More Information \| Similar Motifs Found | motif file (matrix) |
| 7 | T C G A A G T C T C A G G C A T A C T G C G A T A T G C G T C A A G T C C G A T | 1e-14 | -3.268e+01 | 2.55% | 0.47% | 49.4bp (68.8bp) | HEY2/MA0649.1/Jaspar(0.682) More Information \| Similar Motifs Found | motif file (matrix) |
| 8 | T A C G A G T C T G A C G C T A A T G C T C A G A C G T A T C G T C G A A T G C G A C T A T C G | 1e-13 | -3.216e+01 | 2.10% | 0.31% | 45.9bp (65.4bp) | USF1/MA0093.2/Jaspar(0.853) More Information \| Similar Motifs Found | motif file (matrix) |
| 9 | C A T G T A G C G T C A C G T A C T G A G A T C T G A C G C T A T G A C C T G A | 1e-13 | -3.054e+01 | 35.31% | 26.05% | 55.1bp (61.0bp) | RUNX(Runt)/HPC7-Runx1-ChIP-Seq(GSE22178)/Homer(0.869) More Information \| Similar Motifs Found | motif file (matrix) |
| 10 | T A G C C G T A A G T C C T G A C T G A A G T C C T G A A C T G G T A C A C G T | 1e-12 | -2.992e+01 | 3.15% | 0.78% | 53.5bp (64.6bp) | MyoG(bHLH)/C2C12-MyoG-ChIP-Seq(GSE36024)/Homer(0.770) More Information \| Similar Motifs Found | motif file (matrix) |
| 11 \* | T G C A A C T G C T A G T G C A C G T A C T G A A C G T T A C G G T C A A G T C C T A G A C G T | 1e-11 | -2.713e+01 | 1.57% | 0.20% | 55.8bp (60.0bp) | PU.1-IRF(ETS:IRF)/Bcell-PU.1-ChIP-Seq(GSE21512)/Homer(0.706) More Information \| Similar Motifs Found | motif file (matrix) |
| 12 \* | A G C T G T A C C A G T T G A C C G A T A C T G G C A T C G T A A G C T T G A C T G C A G T A C | 1e-11 | -2.713e+01 | 2.55% | 0.57% | 50.4bp (60.1bp) | DMRT3/MA0610.1/Jaspar(0.583) More Information \| Similar Motifs Found | motif file (matrix) |
| 13 \* | G T C A C A T G A T C G C G T A G T A C A C G T A G T C C A G T A G C T A C G T | 1e-11 | -2.641e+01 | 4.42% | 1.56% | 58.2bp (61.1bp) | PROX1/MA0794.1/Jaspar(0.702) More Information \| Similar Motifs Found | motif file (matrix) |
| 14 \* | G T A C C T A G G T A C T G C A A C G T C T A G A G T C C A T G A G T C T G C A T C G A G C T A | 1e-11 | -2.557e+01 | 2.32% | 0.51% | 52.9bp (68.6bp) | NRF1(NRF)/MCF7-NRF1-ChIP-Seq(Unpublished)/Homer(0.868) More Information \| Similar Motifs Found | motif file (matrix) |
| 15 \* | C G A T C G T A A C T G G T C A C A G T C A G T C A T G T C A G G C T A T C G A C G A T T C A G | 1e-10 | -2.440e+01 | 2.02% | 0.41% | 53.3bp (62.3bp) | TEAD3/MA0808.1/Jaspar(0.717) More Information \| Similar Motifs Found | motif file (matrix) |
| 16 \* | A C T G T A G C A G T C C G T A A G T C C G T A C G T A C G T A A C T G A G T C C G T A C G T A | 1e-10 | -2.394e+01 | 0.45% | 0.00% | 53.7bp (53.9bp) | SOX10/MA0442.1/Jaspar(0.684) More Information \| Similar Motifs Found | motif file (matrix) |
| 17 \* | A C G T A G T C C G T A A C T G A C T G A G T C A T G C C T A G A G T C A G T C A C G T C T G A | 1e-9 | -2.270e+01 | 0.37% | 0.00% | 59.2bp (0.0bp) | Srebp2(bHLH)/HepG2-Srebp2-ChIP-Seq(GSE31477)/Homer(0.584) More Information \| Similar Motifs Found | motif file (matrix) |
| 18 \* | C G T A G T C A A C T G A C T G C G T A A T C G C G T A C T G A A G T C C G T A A T G C A T C G | 1e-9 | -2.226e+01 | 0.67% | 0.03% | 69.7bp (60.1bp) | PB0122.1\_Foxk1\_2/Jaspar(0.582) More Information \| Similar Motifs Found | motif file (matrix) |
| 19 \* | T G C A T A G C G T A C A T C G A G T C C G T A A T G C G T C A C A G T C G A T A G T C A G T C | 1e-9 | -2.208e+01 | 1.35% | 0.19% | 49.9bp (59.5bp) | TEAD1/MA0090.2/Jaspar(0.776) More Information \| Similar Motifs Found | motif file (matrix) |
| 20 \* | G A T C A T C G A C T G G C A T A C G T C A G T G T A C A G T C A G C T G C A T | 1e-9 | -2.183e+01 | 1.80% | 0.36% | 61.2bp (62.5bp) | PB0033.1\_Irf3\_1/Jaspar(0.744) More Information \| Similar Motifs Found | motif file (matrix) |
| 21 \* | A C T G C G A T T A G C G T C A C T G A C G T A A C T G A T C G G T A C T A G C | 1e-8 | -2.064e+01 | 2.70% | 0.81% | 50.2bp (60.3bp) | Nr2f6/MA0677.1/Jaspar(0.756) More Information \| Similar Motifs Found | motif file (matrix) |
| 22 \* | T G A C G C A T C A T G C T A G T C A G C G T A C T G A G C A T A C G T A C G T | 1e-8 | -2.045e+01 | 5.32% | 2.41% | 58.0bp (59.2bp) | NFkB-p65-Rel(RHD)/ThioMac-LPS-Expression(GSE23622)/Homer(0.781) More Information \| Similar Motifs Found | motif file (matrix) |
| 23 \* | A G T C A C T G A T G C C T G A G T C A A C G T A G T C C T A G A G T C A T G C | 1e-8 | -1.999e+01 | 1.05% | 0.12% | 56.7bp (62.9bp) | NRF1/MA0506.1/Jaspar(0.643) More Information \| Similar Motifs Found | motif file (matrix) |
| 24 \* | A C T G A C T G A C T G A G T C A C G T C G T A C G T A C G T A A G T C A C G T | 1e-8 | -1.966e+01 | 0.60% | 0.03% | 45.8bp (57.9bp) | BARHL2/MA0635.1/Jaspar(0.603) More Information \| Similar Motifs Found | motif file (matrix) |
| 25 \* | T A C G C A G T A T C G G T A C A T C G A C G T A T G C T A C G T C G A A T G C T A C G A C G T | 1e-8 | -1.926e+01 | 0.37% | 0.01% | 60.0bp (90.2bp) | EGR2/MA0472.2/Jaspar(0.727) More Information \| Similar Motifs Found | motif file (matrix) |
| 26 \* | A C G T C G A T G C A T C G T A C T A G C T A G C G A T A G T C G C T A A C G T | 1e-8 | -1.922e+01 | 2.55% | 0.78% | 55.5bp (60.0bp) | RORA/MA0071.1/Jaspar(0.760) More Information \| Similar Motifs Found | motif file (matrix) |
| 27 \* | C T G A C G T A C T A G A T C G A G T C C G A T A C G T C A G T A T C G G A T C | 1e-8 | -1.910e+01 | 4.80% | 2.14% | 50.5bp (61.4bp) | Nr2e3/MA0164.1/Jaspar(0.720) More Information \| Similar Motifs Found | motif file (matrix) |
| 28 \* | C T A G A T C G A C T G T G A C T G A C A G T C G A C T A G T C | 1e-7 | -1.769e+01 | 14.09% | 9.40% | 53.8bp (58.1bp) | ZNF692(Zf)/HEK293-ZNF692.GFP-ChIP-Seq(GSE58341)/Homer(0.708) More Information \| Similar Motifs Found | motif file (matrix) |
| 29 \* | A C T G T C A G A C G T A C T G C G T A A G T C A C T G A C G T G T A C G T C A | 1e-7 | -1.750e+01 | 1.12% | 0.17% | 52.7bp (65.5bp) | CRE(bZIP)/Promoter/Homer(0.920) More Information \| Similar Motifs Found | motif file (matrix) |
| 30 \* | T C G A C T G A T C A G G T C A T C A G A T G C A T G C C T A G A C T G C G A T | 1e-7 | -1.732e+01 | 1.05% | 0.15% | 52.7bp (64.6bp) | TFCP2/MA0145.3/Jaspar(0.680) More Information \| Similar Motifs Found | motif file (matrix) |
| 31 \* | A G C T C G A T A G T C A C T G A G T C C G A T T G A C C T G A | 1e-6 | -1.394e+01 | 13.94% | 9.81% | 59.4bp (65.2bp) | Gmeb1/MA0615.1/Jaspar(0.645) More Information \| Similar Motifs Found | motif file (matrix) |
